# Endogenous single-cell experiments and stochastic modeling reveal control mechanisms of glucocorticoid receptor dynamics and DUSP1 transcription

**DOI:** 10.64898/2026.07.01.735884

**Authors:** Eric Ron, Alex Popinga, Jack Forman, Luis U. Aguilera, Linda S. Forero Quintero, Brian Munsky

**Affiliations:** Colorado State University, School of Biomedical and Chemical Engineering, Fort Collins, 80523, USA

## Abstract

Glucocorticoids activate the glucocorticoid receptor (GR) to suppress inflammation, yet it remains unclear how GR transport dynamics and downstream gene regulation are coordinated within single cells. We combine immunocytochemistry (ICC) and single-molecule fluorescence in situ hybridization (smFISH) to quantify endogenous GR transport and *DUSP1* transcription dynamics across thousands of individual cells following dexamethasone (Dex) stimulation. Performing multiple rounds of statistical inference based on Chemical Master Equations (CME), we determine the most likely mechanisms and reaction rates for Dex-driven GR nuclear import; compartment-specific GR degradation; GR-dependent control of the *DUSP1* promoter; and DUSP1 transcription, elongation, transport, and degradation. Our inferred model suggests that nuclear GR degradation is the dominant mechanism of receptor clearance, that GR primarily regulates promoter activation, and that time-dependent AU-rich element (ARE)–mediated mRNA degradation contributes heavily to *DUSP1* clearance. With these mechanisms, the fully-parameterized model quantitatively predicts joint distributions of GR translocation and decay dynamics, *DUSP1* transcription site activity, and nuclear and cytoplasmic DUSP1 mRNA heterogeneity among clonal cells as functions of time and across seven orders of magnitude for Dex induction concentrations. Our results establish an integrated quantitative framework to link receptor dynamics to gene expression heterogeneity and predict single-cell hormone-responsive transcription programs.

## Introduction

Glucocorticoids (GCs) are cornerstone therapeutics for the treatment of inflammatory and autoimmune diseases, acting through the glucocorticoid receptor (GR) to regulate hundreds of target genes across nearly all tissues^1–4^. Dexamethasone (Dex) is a high-affinity, long-acting synthetic GC that potently activates GR and serves as a benchmark ligand for mechanistic studies^5^. We focus on HeLa cells, which despite a problematic history^6^, are a well-characterized epithelial model with consistent nuclear–cytoplasmic morphology to resolve the subcellular and transcriptional processes underlying Dex-dependent anti-inflammatory responses^5–7^. Among GR-induced genes, *Dual Specificity Phosphatase 1 (DUSP1)* encodes MAPK phosphatase 1 (MKP1), which dephosphorylates and inactivates the stress kinases p38 and JNK, forming a key negative-feedback loop on inflammatory MAPK signaling (Fig. 1a)^8,9^.

**Figure 1.**
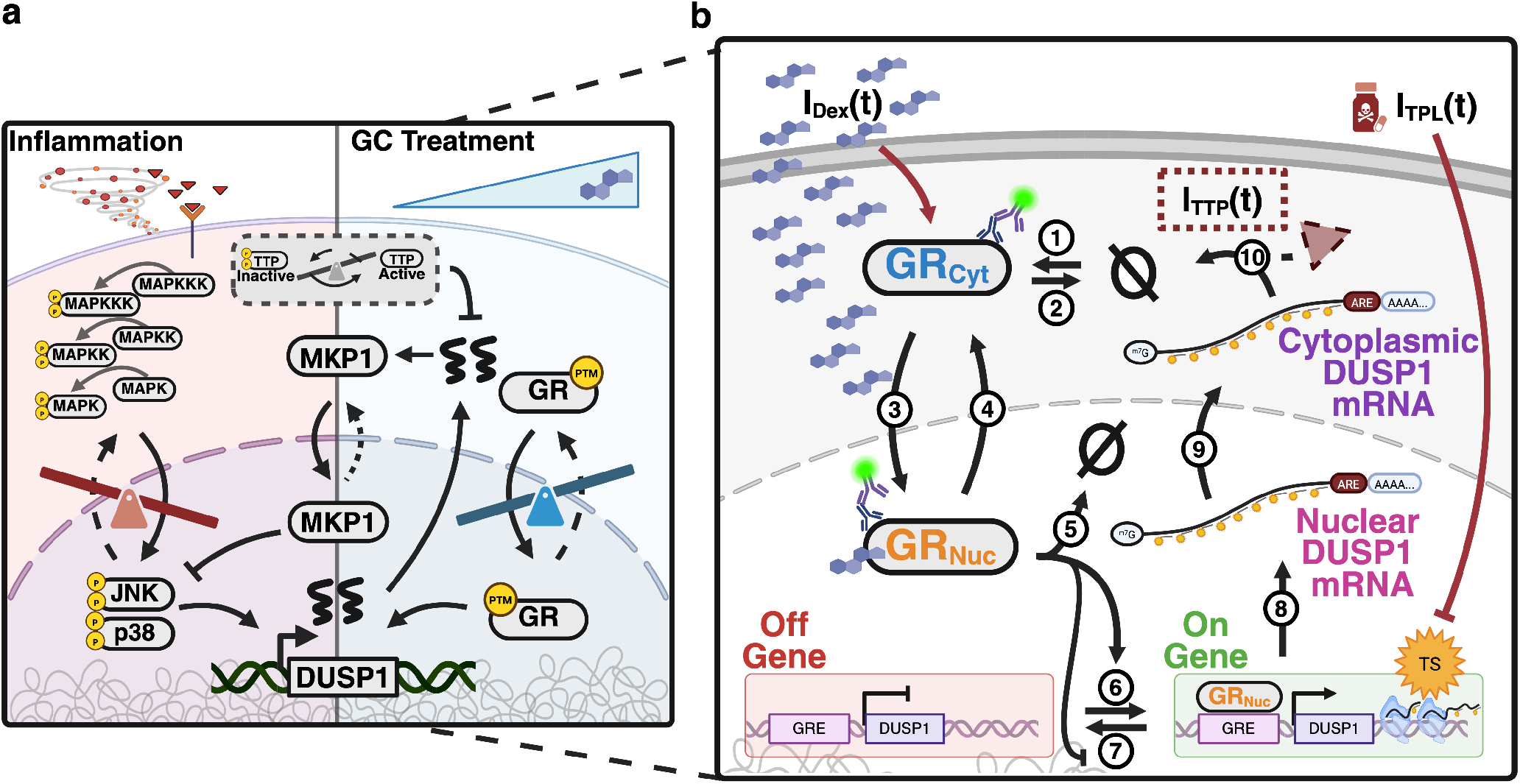
**(a)** Inflammatory signaling activates JNK and p38, inducing *DUSP1* transcription and p38-dependent TTP phosphorylation that stabilizes ARE containing mRNAs. MKP1 dephosphorylates JNK and p38, forming a negative feedback loop. GCs up regulate *DUSP1* via GR to suppress MAPK signaling. **(b)** Model schematic. HeLa cells are stimulated with multiple concentrations of Dex over three hours to generate a time-dependent Dex signal *I*_Dex_(*t*). The model accounts for: **(1)** GR synthesis and **(2)** cytoplasmic GR degradation; **(3)** Dex-induced GR ranslocation to the nucleus; **(4)** nuclear–cytoplasmic shuttling of GR; and **(5)** nuclear GR degradation. Within the nucleus, GR binds to GREs to activate the *DUSP1* promoter, **(6**,**7)** modeled as stochastic switching between OFF and ON states. **(8)** Transcription produces nascent nuclear DUSP1 mRNA at transcription sites (TS), which are detected by smFISH, followed by **(9)** export of *DUSP1* mRNA to the cytoplasm and **(10)** ARE-mediated degradation through *I*_TTP_(*t*). In transcription inhibition experiments, a secondary signal *I*_TPL_(*t*) represents TPL-mediated suppression of mRNA synthesis during dual Dex–TPL stimulation. Red arrows indicate drug stimulation.

The GR is a modular transcription factor whose structure dictates both localization and activity. The ligand-binding domain anchors unliganded GR in cytoplasmic chaperone complexes with HSP90 and FKBP51; the DNA-binding domain recognizes Glucocorticoid Response Elements (GREs); and the intrinsically disordered N-terminal domain harbors activation function-1 and post-translational modification sites that modulate transcriptional potency, nuclear residency, and receptor stability^10–15^. Upon ligand binding, GR undergoes conformational changes that expose nuclear localization sequences (NLS1 and NLS2), enhancing importin-*α*/*β* mediated nuclear import and suppressing CRM1-dependent export^16,17^. Once in the nucleus, GR binds GREs to activate target genes, but is also subject to ongoing nucleocytoplasmic shuttling and ligand-dependent proteasomal degradation^7,18–20^. The compartment in which GR is cleared remains unresolved, and the rates of nuclear and cytoplasmic turnover have rarely been quantified. Although export-mediated cytoplasmic turnover has been proposed^16,21^, more recent evidence supports proteasomal degradation within the nucleus^20,22,23^, and nuclear turnover has been reported for estrogen and androgen receptors^24–26^. Despite previous efforts, substantial gaps preclude predictive understanding for the balance of nuclear import, receptor recycling, chromatin engagement, and compartment-specific degradation to determine the magnitude, duration and heterogeneity of transcriptional response dynamics in endogenous cells.

The human *DUSP1* gene resides at 5q35.1, spans approximately 3.1 kb on the reverse strand of chromosome 5 (GRCh38), and comprises four exons and three introns^27^ (Supplementary Fig. S1). At GR-regulated promoters, single-molecule and biochemical studies of the mouse mammary tumor virus (MMTV) model system have shown that GR can activate transcription through several non-exclusive mechanisms, including increased burst frequency^28^, stabilization of the active promoter state^12,29^, and enhanced transcription rates within bursts^12,30^. Consistent with these mechanisms, multiple candidate GREs have been mapped upstream of the *DUSP1* transcription start site in A549 cells^31^, and at least one site is enriched for both GR and the coactivator p300 alongside increased chromatin accessibility and histone acetylation^32^, though GR binding and chromatin context can vary across cell types^33^. Nascent *DUSP1* transcripts accumulate at transcription sites (TS) and appear as bright nuclear foci under smFISH imaging (Supplementary Fig. S2), with intensities brighter than multiples of the average single diffraction-limited mRNA^34^ (Supplementary Fig. S1). Following processing and export through nuclear pore complexes, mature *DUSP1* mRNAs enter the cytoplasm, where they are translated into MKP1 or targeted for degradation. The *DUSP1* mRNA contains AU-rich elements (AREs) in its 3’ UTR (Supplementary Fig. S1), making it a substrate for ARE-binding proteins, such as tristetraproline (TTP), which can accelerate mRNA turnover^9,35–38^. Consequently, ligand-induced *DUSP1* expression arises from a balance between GR-driven transcriptional activation, receptor down-regulation, and cytoplasmic mRNA decay.

Previous modeling efforts have captured important aspects of GR-mediated transcriptional regulation using a range of approaches. Equilibrium thermodynamic models of ligand–GR–coregulator–DNA interactions have described how allosteric coupling gives rise to ligand- and gene-specific transactivation at steady state^39^. ODE models have simulated the kinetics of GR autoregulation and induction of target genes such as *BIM* and *GILZ* in GC-sensitive and -resistant leukemia cell lines^40^. Multivariate statistical approaches have demonstrated that GR Ser211 phosphorylation is a stronger predictor of endogenous target gene responses than conventional reporter assays^41^, and Boolean models of the GR interactome have provided qualitative predictions of signaling states associated with GC sensitivity and resistance^42^. However, these approaches either operate at steady state or on population-averaged quantities, and none integrates subcellular spatial resolution with stochastic single-cell dynamics to mechanistically link GR activation to the heterogeneous transcriptional responses observed at the single-cell level.

To complement and expand upon previous investigations of GR and DUSP1, our approach integrates single-cell imaging of endogenous targets with stochastic modeling to capture the spatial and temporal dynamics of GR signaling coupled with downstream regulation of DUSP1 transcription. Using fixed-cell ICC and smFISH measurements of GR localization and *DUSP1* mRNA abundance at many time points during responses to different stimulation dosages (see Methods), we quantify the time-varying distributions of inter- and intracellular heterogeneity that are inaccessible to bulk assays, live-cell reporters, or exogenous constructs. Previous work in other systems has demonstrated that systematic integration of measured and modeled temporal and spatial cell-to-cell variability can provide sensitive *fluctuation fingerprints* to more precisely interrogate gene regulatory mechanisms and their parameters^43–49^. Inspired by these efforts, we use a chemical master equation (CME) framework to explicitly model the probabilistic nature of transcription, transport, and degradation with a framework that reproduces not only means but the full distributions of these measured responses. By statistically comparing many different model hypotheses against the data, we infer the most probable mechanisms and parameters to achieve a highly-predictive quantitative model for GR nuclear import and degradation, GR-modulated promoter activation, DUSP1 transcriptional elongation, and ARE-mediated mRNA decay (Fig. 1b). This integrative modeling approach clarifies how GR-dependent transcriptional and post-transcriptional processes jointly shape the heterogeneous single-cell responses observed across Dex stimulation and triptolide (TPL)-mediated transcription inhibition conditions.

## Results

We begin by proposing a semi-mechanistic stochastic model abstraction for the coupled GR–DUSP1 system as summarized in Fig. 1b. The model is comprised of three modules: (*i*) GR transport regulation defined by **reactions 1-5**, (*ii*) DUSP1 transcription and transport regulation defined by **reactions 6-9**, and (*iii*) DUSP1 degradation regulation defined by **reaction 10**. The mechanisms and parameters of these modules are sequentially inferred from single-cell ICC (Module *i*) and smFISH data (Modules *ii* and *iii*) as described in the following sub-sections.

### Subcellular ICC measurements reveal heterogeneous GR translocation dynamics

To quantify endogenous GR dynamics in response to GC stimulation, HeLa cells were treated with 1, 10, or 100 nM Dex for six different time periods (10, 30, 50, 75, 120, 180) min in addition to an unstimulated control. Representative ICC images (Fig. 2a), which were verified against isotype and secondary control experiments (Supplementary Fig. S3), illustrate three key features of GR localization. First, unstimulated HeLa cells exhibit a mean estimated nuclear-to-cytoplasmic (N/C) mass ratio of 0.32 ± 0.10 (STD), reflecting moderate nuclear GR in most cells (Fig. 2a,d,e, Supplementary Table 1) and consistent with prior reports of nuclear GR localization in unstimulated cells^16,50,51^. Second, at 1 nM Dex, the population N/C ratio rises slowly and reaches only 0.90 ± 0.22 by 180 min, whereas 10 and 100 nM produce faster and significantly larger shifts, with peak N/C ratios of 5.23 ± 3.99 at 180 min for 10 nM and 6.63 ± 3.49 at 30 min for 100 nM (Supplementary Fig. S4, Supplementary Table S1). Third, the total amount of GR decreases across all conditions, with faster rates of depletion for higher concentrations of Dex (Fig. 2e).

**Figure 2.**
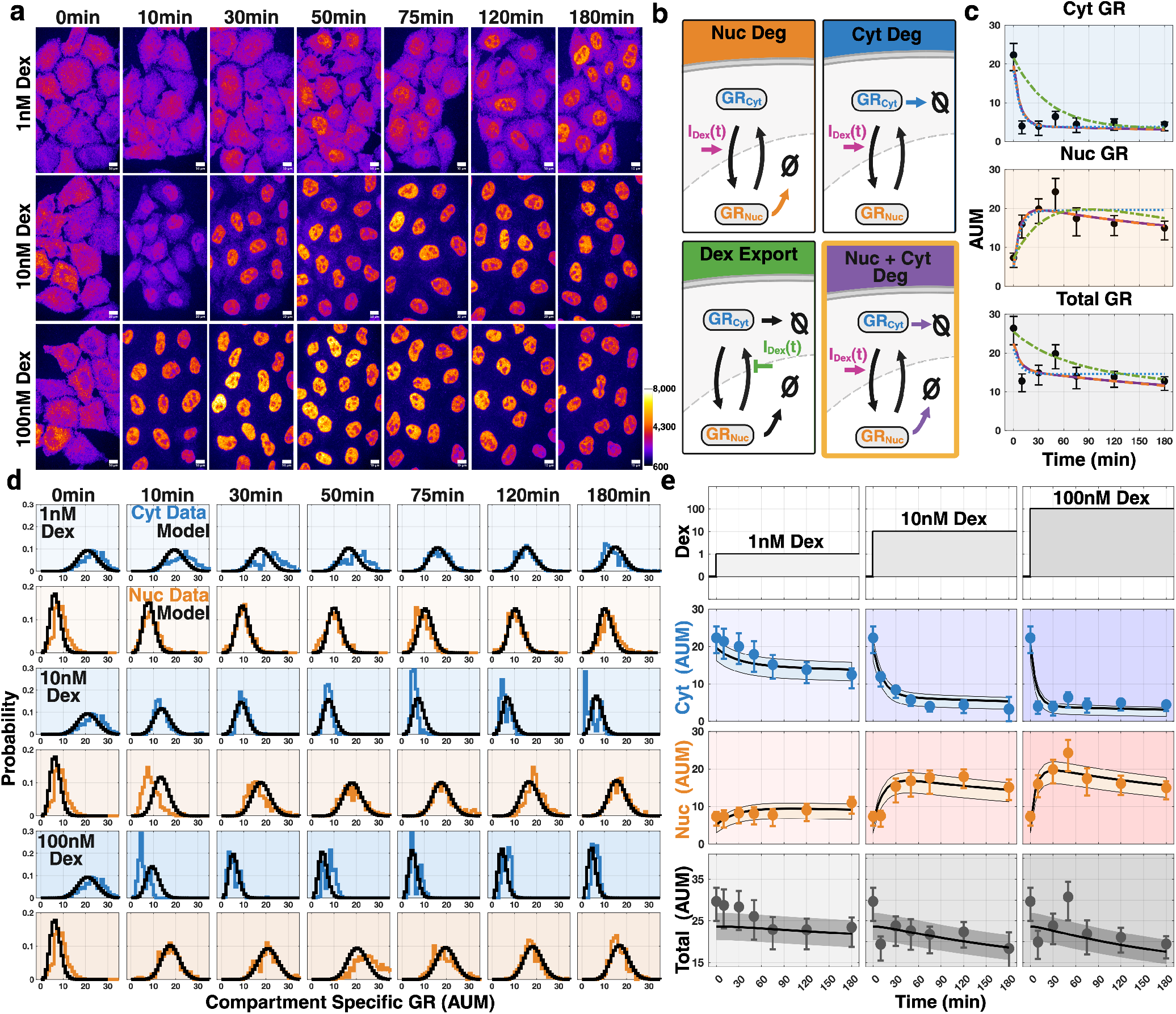
Measurement and modeling of GR production, transport, and degradation dynamics. **(a)** Representative HeLa cells stained for GR by ICC at selected times following continuous Dex stimulation (1 nM, 10 nM, 100 nM). Intensities scaled by background. All scale bars are 10 µm. **(b)** Candidate GR models, each comprising cytoplasmic and nuclear GR species with Dex stimulation *I*_Dex_(*t*) driving nuclear export (green) or nuclear import with GR degradation in the nucleus (Nuc Deg, orange), cytoplasm (Cyt Deg, blue), or both nucleus and cytoplasm (Nuc + Cyt Deg, purple). The Nuc + Cyt Deg model provides the best fit. **(c)** Comparison of model fits (colored as in (b)) to cytoplasmic (Cyt GR), nuclear (Nuc GR), and total (Total GR) GR versus time for 100 nM Dex stimulation compared measured medians and upper/lower quartile ranges (black, circles and bars). **(d)** Measured distributions of cytoplasmic (blue) and nuclear (orange) GR concentration at each time point and Dex dose (1 nM, 10 nM, 100 nM), compared with FSP model fits (black). **(e)** Dex input profiles (top) and temporal trajectories of 25/50/75 percentiles for cytoplasmic GR (AUM, blue), nuclear GR (AUM, orange), and total GR mass (AUM, gray) at each dose. Measured medians and upper/lower quartile ranges are shown by circles and bars, and the corresponding model fits are shown by solid lines and shading.

#### A stochastic model consisting of Dex-driven nuclear import and compartment-specific degradation captures the heterogeneous response of GR spatial and temporal dynamics

To interpret the measured GR spatial dynamics (i.e., ***CytGR*** and ***NucGR***) as functions of time and Dex concentration, we constructed a flexible GR model (Module *i*) and used it to compare competing hypotheses for how Dex controls GR transport and where GR is degraded (Fig. 2b, Supplementary Fig. S5).

The model couples cytoplasmic and nuclear GR through nucleocytoplasmic transport, with Dex supplied as a time-varying signal 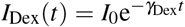, where *I*_0_ is the supplied Dex concentration and *γ*_Dex_ is its first order decay rate, and with Dex-stimulated transport following Michaelis-Menten kinetics to allow for saturation at higher concentrations. We evaluated four candidate models (Fig. 2b, Supplementary Fig. S5): three in which Dex drives nuclear import while GR degradation is (*i*) restricted to the nucleus, (*ii*) restricted to the cytoplasm, or (*iii*) active in both compartments (with independent rates), and (*iv*) one in which Dex suppresses nuclear export instead of import.

To distinguish among these candidates, we fit each model independently to the single-cell ICC data simultaneously for all three Dex concentrations. By solving the CME using Finite State Projection (FSP)^52,53^ and searching over parameter space, we found the maximum a posterior (MAP) estimate for each model, and we accounted for differences in model complexity using the Bayesian Information Criteria (BIC, Supplementary Table S2). The Dex export model gave the poorest agreement and was unable to match the rapid rate of cytoplasm to nucleus GR transport (Supplementary Fig. S5), while the cytoplasm-only degradation model overestimated nuclear GR at later times and could not reproduce the decrease in total GR (Fig. 2c, Supplementary Fig. S5). The nucleus-only and compartment-specific degradation models produced reasonable fits to the cytoplasmic, nuclear, and total GR dynamics (Fig. 2c, Supplementary Fig. S5), but the compartment-specific model achieved the highest MAP of the four candidate models and the lowest BIC after penalizing for its additional degradation parameter (Supplementary Table S2). An approximate Bayes factor (BF) reached the same conclusion, favoring the compartment-specific model by over e^100^-fold over the next-best nucleus-only model (Supplementary Table S2).

Figure 2d shows the resulting fits to the nuclear and cytoplasmic GR distributions as functions of time and Dex concentration, and Figure 2e summarizes these distributions to show the measured and model-fitted median, 25^th^ and 75^th^ percentile levels for cytoplasmic, nuclear, and total GR concentrations for all Dex concentrations and times. Supplementary Table S3 shows the inferred maximum a posterior (MAP) estimates and inferred parameter uncertainties inferred using Metropolis Hastings (MH) assuming broad lognormal prior distributions with standard deviations of two orders of magnitude for each parameter (see methods), while Supplementary Figures S6 and S7 show the corresponding joint and marginal posterior distributions, respectively.

By comparing the rates of nuclear and cytoplasmic GR degradation inferred from the ICC data (*γ*_GRnuc_ and *γ*_*GRcyt*_ in Supplementary Table S3), we found that nuclear degradation was faster by approximately 3-fold, supporting ligand-induced nuclear proteasomal degradation as a key mechanism of GR deactivation^7,19,54,55^. Consistent with this interpretation, the cytoplasmic GR antibody intensity did not reappear at later times, and total GR levels declined more rapidly at higher Dex concentrations, when a greater fraction of GR was in the nucleus (Fig. 2d,e). Inferred transport rates also recapitulated the expected pattern of ligand-dependent shuttling^16,17^: under unstimulated conditions, the nuclear export rate (*k*_nc_ = 0.0188 ± 0.00066 min^*−*1^) exceeded the basal import rate (*k*_cn0_ = 0.0060 ± 0.00016 min^*−*1^), favoring cytoplasmic localization, while 100 nM Dex-stimulated import (*k*_cn0_ + *k*_cn1_*I*_Dex_(*t*) *≈* 11.5 min^*−*1^) was much higher (Supplementary Table S3), driving rapid and nearly complete nuclear accumulation. Together these findings reproduced the observations that 1 nM stimulation maintained the highest total GR (Nuc + Cyto) across all stimulatory conditions, while 100 nM produced the fastest and most uniform transport induction followed by the most rapid decline in ***NucGR, CytGR***, and ***Total GR*** (Fig. 2e).

#### Single-cell-single-mRNA measurements reveal stochastic DUSP1 transcription dynamics

To quantify early transcription of endogenous *DUSP1* mRNA, we designed a smFISH probe set targeting exonic regions of the human *DUSP1* transcript (Supplementary Fig. S1)^56^. *GAPDH* smFISH probes served as a cytoplasmic marker to enable automated single-cell segmentation (Fig. 3a, Supplementary Fig. S2). Images were analyzed using a custom implementation of BigFISH^34^ (see Methods). A combined compartment-based and local signal-to-noise ratio (SNR) thresholding strategy yielded the most reliable spot detection, as verified visually (SNR Methods) (Supplementary Fig. S2). This approach substantially reduced false positives and enabled consistent identification of transcription sites (magenta), nuclear mRNAs (blue), and cytoplasmic mRNAs (purple) (Fig. 3a), with minor expected imperfections (Supplementary Fig. S2). Once identified, candidate spots were post-processed and separated into TS’s and mature mRNA (Fig. 3a, Supplementary Fig. S2).

**Figure 3.**
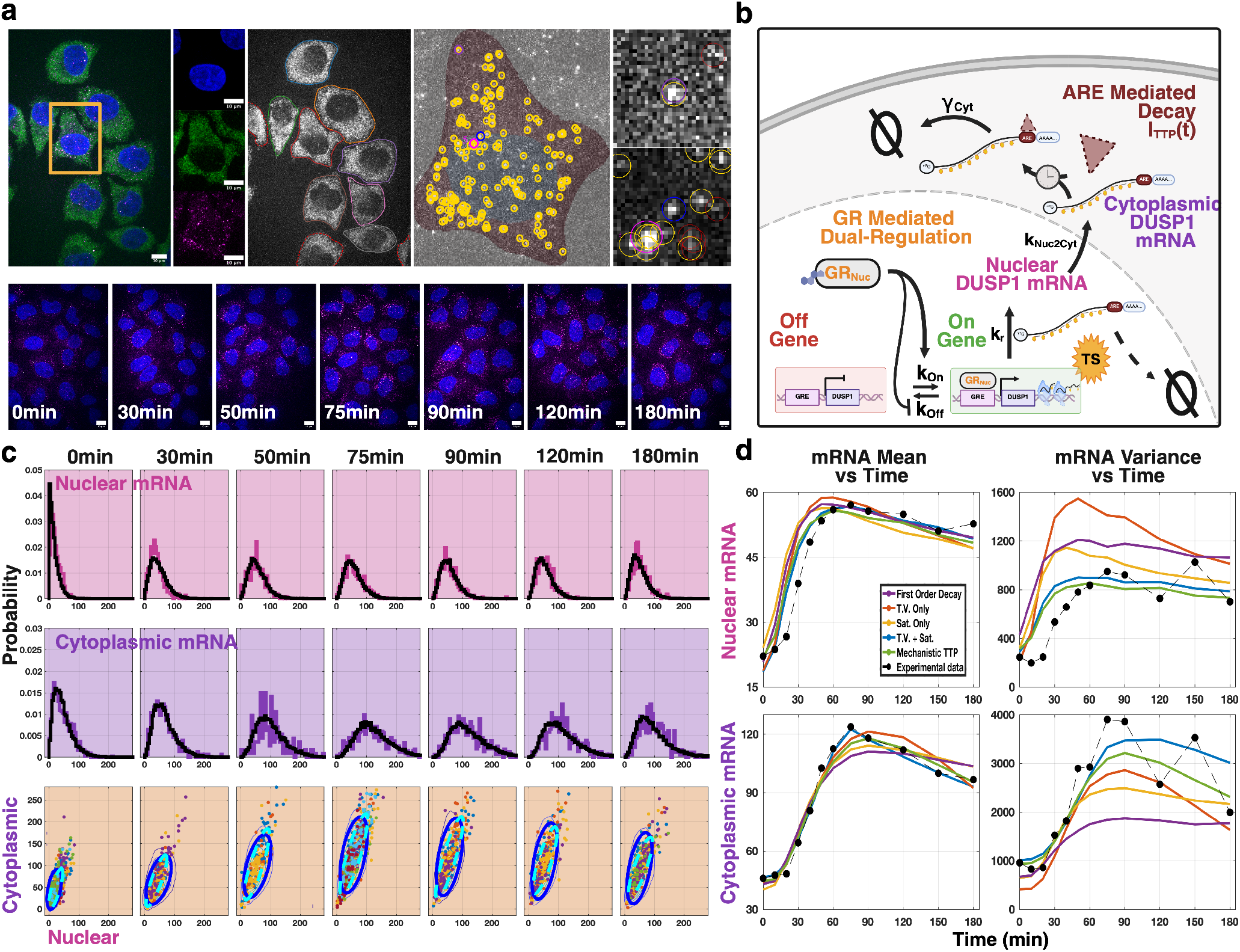
Measurement and modeling of nuclear DUSP1 activation. **(a)** Single-cell segmentation and smFISH spot detection workflow. From left to right: merged three-color image of a representative field of view; zoomed view of a selected cell (gold box) with individual DAPI, *GAPDH* smFISH, and *DUSP1* smFISH channels; cytoplasmic segmentation with partial peripheral cells excluded; nuclear and cytoplasmic segmentation overlaid with detected *DUSP1* mRNA; example spot crops with cytoplasmic (top) and nuclear (bottom) detections. Gold circles indicate accepted mRNA detections, magenta circles denote transcription sites, and red circles mark low-SNR candidates removed during filtering. Bottom: representative merged *DUSP1* smFISH and DAPI images following 100 nM Dex stimulation at the indicated time points. All scale bars are 10 µm. **(b)** Schematic of the stochastic model for GR-dependent control of to *DUSP1* mRNA production, nuclear export, and ARE-mediated cytoplasmic degradation. **(c)** Distributions of nuclear (top) and cytoplasmic (middle) and join (bottom) *DUSP1* mRNA at the indicated time points and compared to corresponding model fits. Ellipses in the bottom row show 68% confidence intervals as predicted by the model (solid blue) or measured in the data (dashed cyan). **(d)** Mean (left column) and variance (right column) of nuclear (top row) and cytoplasmic (bottom row) *DUSP1* mRNA over time, comparing experimental measurements to candidate model fits (First Order Decay, T.V. Only, Sat. Only, T.V. + Sat., Mechanistic TTP).

At 100nM stimulation, and across all time points, *DUSP1* expression remained heterogeneous at the single-cell level (Fig. 3c,d, Supplementary Fig. S8). The fraction of cells that had one or more active TS increased rapidly after stimulation and peaked at roughly 30–40 minutes, while the intensity of active TS remained relatively constant (Fig. 4a). TS activation was followed by the rise in nuclear mRNA, which reached a maximum near 75 minutes (Fig. 4a). Nuclear and cytoplasmic mRNA levels then peaked between 75–90 minutes (Fig. 3d, Fig. 4a) before declining, with a more pronounced decrease observed in the cytoplasm (Fig. 3c,d). DUSP1 mRNA distributions were highly variable with Fano factors (i.e., variance/mean) of 11.1 ± 2.9 (at 0 min) to 16.6 ± 2.5 (at 90 min) for nuclear mRNA and with much higher cytoplasmic Fano factors of 20.9 ± 3.5 (at 0 min) to 32.7 ± 6.4 (at 90 min).

**Figure 4.**
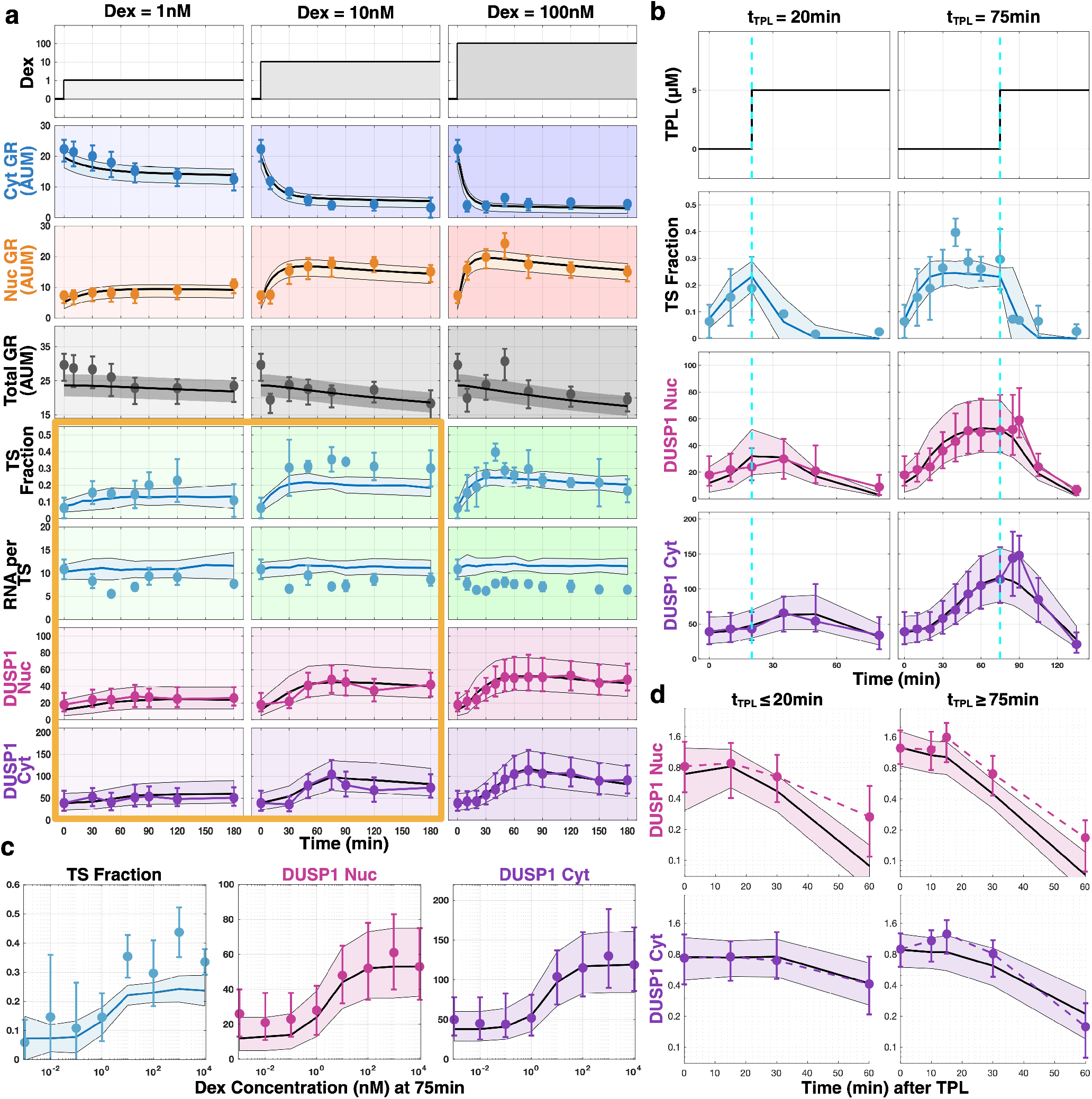
Model predictions for *DUSP1* transcriptional dynamics. (a)GR and *DUSP1* dynamics at 1, 10, and 100 nM Dex. The top three rows show fits to cytoplasmic GR concentration (AUM, blue), nuclear GR concentration (AUM, orange), and total GR mass (AUM, gray). The bottom four rows show *DUSP1* TS fraction, RNA per TS, nuclear mRNA (*DUSP1* Nuc), and cytoplasmic mRNA (*DUSP1* Cyt). The semi-mechanistic model was fit to the GR data and the 100 nM Dex column; the gold outline marks the 1 and 10 nM columns, which are predictions generated without refitting. **(b)** Dex–TPL dual-stimulation predictions for early (*t*_TPL_ = 20 min) and late (*t*_TPL_ = 75 min) transcriptional inhibition. Cells were first stimulated with 100 nM Dex, followed by 5 µM TPL added at *t*_TPL_; cyan dashed lines mark TPL addition. **(c)** Dex-titration predictions at *t* = 75 min spanning 1 pM to 10 µM. **(d)** Compartment-specific *DUSP1* mRNA decay after TPL, grouped into early (*t*_TPL_ *≤* 20 min) and late (*t*_TPL_ *≥* 75 min) inhibition. The y-axis is compartment-specific mRNA normalized to one at the predicted average time for the combined process of TPL onset, *DUSP1* elongation, and nuclear-to-cytoplasmic export. Cytoplasmic decay rates increase substantially at later times while nuclear rates remain similar, consistent with time-dependent ARE-mediated turnover. Panels (b)–(d) are predictions from the semi-mechanistic model without refitting. In (a) and (c), TS fraction and RNA per TS are mean ± standard deviation; all other data show median ± interquartile range (25th/75th percentiles) for data (markers/bars) and model (lines and shading).

#### Stochastic modeling shows that nuclear GR increases DUSP1 transcription burst frequencies

To build a quantitative understanding for the control of *DUSP1* regulation, we extended the GR model from above (Module *i*) with a transcription and nuclear-transport module (Module *ii*) comprising three additional biochemical species (Fig. 1b): inactive gene copies (***offGene***), active gene copies (***onGene***), and nuclear mRNA transcripts (***rNuc***). To capture the stochastic dynamics of these species, we adopt the standard two-state bursting gene expression model^43^. In this model, ***offGene, onGene***, and ***rNuc*** populations are controlled through four additional reactions: allele activation (**reaction 6**: *→* ***offGene onGene***), allele deactivation (**reaction 7**: ***onGene*** *→****offGene***), transcription (**reaction 8**: *→* ***onGene onGene*** + ***rNuc***), and nuclear degradation or transport of mRNA from the nucleus to the cytoplasm (**reaction 9**: ***rNuc****→* ***∅***, or ***rNuc → rCyt***). For (**reaction 8**), the model allows for different transcription rates for when alleles are in the ***offGene*** or ***onGene*** states. The model was assumed to have a total of two *DUSP1* alleles per cell, consistent with observations that the vast majority (98.9%) of cells were detected to contain zero, one, or two observable TS per cell.

We reasoned that the finding that application of 100 nM Dex leads to increased *DUSP1* activity could be explained by multiple distinct mechanisms in the context of the bursting gene expression model. GR could increase the promoter activation rate (*k*_on_ in **reaction 6**), decrease the promoter deactivation rate (*k*_off_ in **reaction 7**), or increase the transcription rate for the ON state (*k*_r_ in **reaction 8**), or act through some combination of these effects (Supplementary Fig. S10). Depending upon model parameters, these mechanisms can predict distinct and quantitatively revealing relationships between the fraction of cells with active TS and the distributions of nascent mRNA per active TS or mRNA counts per cell^43,45^.

To select among the proposed models, we searched over parameter space to maximize the a posterior probability of the smFISH data for the number of mRNA per cell and the distribution for the total intensity of the transcription sites (or in the rare 1.1% of cases where more than two TS are detected, the total of the two brightest TS). The resulting MAP-based comparisons strongly favored GR regulation of the promoter activation rate *k*_on_ as the dominant mechanism: the Bayes factor in favor of *k*_on_ over *k*_r_ alone was *LR* = *e*^334^, and over *k*_off_ alone was *LR* = *e*^128^ (Supplementary Table S4). Consistent with this, a model in which GR acts exclusively through *k*_on_ achieved the best MAP for TS abundance among all single-action models, while the combined *k*_on&_*k*_off_ model marginally better overall MAP fits and BIC/BF values (Supplementary Table S4). In the smFISH data, we found that Dex stimulation produces a substantial increase in the fraction of *DUSP1* alleles exhibiting detectable TS, while the mean and the distributions of TS intensity show relatively small changes over time (Fig. 4a), a pattern that is most consistent with increased burst frequency through increases in *k*_on_ rather than increased burst size through decreases in *k*_off_ or increases in *k*_r_^45^.

Although the single strongest regulatory interaction was estimated to be an increase in *k*_on_, models that also included GR-dependent decreases in *k*_off_ yielded modest improvements (Supplementary Table S4). Models incorporating GR-dependent changes in *k*_r_ provided no meaningful improvement in fit, with the full three-parameter model (*k*_on_*/k*_off_*/k*_r_) performing comparably to the two-parameter model and carrying a higher BIC penalty. Our final model therefore retains GR regulation of *k*_on_ as the dominant mechanism (thick line), with potential for weaker modulation of *k*_off_ (thin line) (Fig. 1b, Supplementary Fig. S9). The final inferred set of semi-mechanistic model parameters and uncertainties is presented in Supplementary Table S5.

#### Subcellular measurements reveal complex dynamics of DUSP1 mRNA transport and degradation

To extend our GR and *DUSP1* promoter-level analyses to include the full *DUSP1* mRNA life cycle, we next examined the time-varying relationship between nuclear and cytoplasmic mRNA distributions (Fig. 3c,d, Supplementary Fig. S8). At early times, following a short delay after addition of Dex, the average level of cytoplasmic mRNA increases rapidly to reach a maximum level shortly after that of the nuclear mRNA. These observations were expected since cytoplasmic mRNA accumulation follows behind that of the nucleus with a delay set by the nucleus to cytoplasm transport rate. However, upon comparing the distributions of nuclear and cytoplasmic mRNA at later times (Fig 3c,d, Supplementary Fig. S8), we made two surprising key observations: First, after reaching its maximum value, the average level of cytoplasmic mRNA rapidly decreases, which is surprising given that the average level of nuclear mRNA decreases only slightly below its maximal value. Second, at later times, the relative level of heterogeneity in the number of cytoplasmic mRNA (e.g., Fano factor *FF*_cyt_ = 32.7 6 ±.4 at *t* = 90 min) is much wider than that of the nuclear mRNA (*FF*_nuc_ = 16.6 ± 2.51), which was surprising given that the cytoplasmic mean number (*µ*_cyt_ = 118.0) is more than twice as large as that of nuclear mRNA (*µ*_nuc_ = 55.7) at the same time.

#### Spatial stochastic modeling suggests that ARE-mediated decay is the dominant mechanism for cytoplasmic DUSP1 mRNA clearance

To extend our model to capture these observations, we added a cytoplasmic degradation module (Module *iii*) with an additional species for cytoplasmic mRNA (***rCyt***), which increases via the previous transport reaction (**reaction 9**) and decays according to **reaction 10** (*rCyt →* ∅). During preliminary parameter inference, we found that the nuclear mRNA degradation pathway (Fig. 3b, dashed line) was consistently driven to negligible values, and thus we only consider cytoplasmic decay from this point onward.

We proposed and evaluated four candidate cytoplasmic decay models of varying levels of complexity (Supplementary Fig. S10): (i) a standard model of constant first-order decay, (ii) a semi-mechanistic model where the degradation rate saturates at high levels of mRNA, (iii) a model where the degradation rate increases over time, and (iv) a model incorporating both saturation and time-dependence in the degradation rate. Mechanism (ii) was inspired by the work of Hansen et al^49^, who demonstrated that higher variability in the cytoplasm could be explained by a biphasic degradation mechanism, where distinct subsets of mRNA exhibit different degradation rates. For example, ARE–dependent cytoplasmic degradation could lead to increased degradation of mRNA but only if sufficient quantities of ARE-binding proteins are available. Similarly, mechanism (iii) was inspired by the knowledge that the ARE-binding protein TTP is known to be induced by Dex^37,57,58^.

Upon fitting the extended models to simultaneously capture TS intensities, nuclear mRNA, and cytoplasmic mRNA (see methods), we found that all degradation models could reproduce the average levels of nuclear and (to a lesser extent) the cytoplasmic mRNA as functions of time (Fig. 3d, top left and bottom left, Supplementary Fig. S10). The models diverged sharply, however, in their ability to reproduce the heterogeneity of the mRNA distributions. Without a saturation term to constrain the degradation rate, the simpler models could not capture the joint variability in both compartments, and their best fits substantially over-estimated nuclear variability (Fig. 3d, top right) or under-estimated cytoplasmic variability (Fig. 3d, bottom right). Only the semi-mechanistic model incorporating both time-varying and saturating degradation (T.V. + Sat.) and the full mechanistic DUSP1-TTP model (Supplementary Fig. S11) captured the nuclear and cytoplasmic variances while simultaneously fitting both means (Fig. 3d, Supplementary Fig. S10). These qualitative observations were supported by quantitative comparisons of the models’ fits to the data (Supplementary Table S6), showing that the best fits consistently and overwhelmingly favored the selection of a degradation mechanisms that includes both time dependence and saturation with respect to cytoplasmic mRNA abundance. This result matches closely to our key qualitative observations above and illustrated in Figure 3b. First, the time dependent increase in the cytoplasmic decay rate is needed to achieve the rapid average reduction in cytoplasmic mRNA that is observed at later times. Second, the saturation effect on the degradation rate is needed to allow mRNA to survive longer when they are already more abundant, which leads to a longer tail in the cytoplasmic mRNA distribution (Fig. 3c, Supplementary Fig. S8).

Together, these results suggest that ARE-mediated, time-varying, and saturable cytoplasmic degradation is the dominant mechanism governing *DUSP1* mRNA clearance following GC stimulation (Fig. 3b, Supplementary Fig. S10). This Dex-dependent, but variable, acceleration of cytoplasmic mRNA turnover is consistent with recent studies demonstrating enhanced *DUSP1* degradation through ARE-binding proteins such as TTP^37^.

#### The inferred Dex-GR-DUSP1 model accurately predicts *DUSP1* transcriptional dynamics across Dex concentrations and transcriptional inhibition conditions

After fitting to the single-cell GR ICC experimental measurements at 1, 10, and 100 nM Dex to constrain GR mechanisms and parameters and the smFISH measurements at 100 nM Dex to constrain *DUSP1* regulation mechanisms and parameters, we next asked if our fully-parameterized model could quantitatively predict *DUSP1* transcriptional outputs across other Dex concentrations. To extend the model to the new levels of Dex without introducing new parameters or mechanisms, we assumed that the impact of Dex on the time variation of cytoplasmic degradation should scale linearly with the increase of GR in the nucleus (see Methods). Model predictions for TS fraction, RNA per TS, and nuclear and cytoplasmic *DUSP1* mRNA were validated against experimental measurements at both 1 nM and 10 nM Dex across all sampled time points (Fig. 4a). Importantly, these predictions capture both mean behaviors and observed variances, indicating that the stochastic structure of the model accurately extrapolates to predict cell-to-cell heterogeneity (Supplementary Figs. S12,S13).

To further challenge the model’s ability to predict promoter activity, we performed a Dex concentration sweep spanning seven orders of magnitude and measured *DUSP1* mRNA abundance at a fixed stimulation time of 75 min. Without refitting, the model accurately predicted both nuclear and cytoplasmic *DUSP1* mRNA counts across concentrations ranging from 1 pM to 10 µM (Fig. 4c). The resulting dose–response relationship exhibits a characteristic sigmoidal shape, reflecting saturation of GR transport, GR-dependent promoter activation, and downstream transcriptional output. We confirmed that our final model consistently made more accurate predictions than any of the other discarded models (Fig. 3d, Supplementary Fig. S10, and Table S6).

To validate the inferred cytoplasmic decay mechanisms, we next examined the model’s ability to predict *DUSP1* dynamics under transcriptional inhibition. Dual-stimulation experiments were performed in which cells were first treated with 100 nM Dex and subsequently exposed to 5 µM TPL at defined times following Dex addition. TPL is known to block transcriptional initiation, providing a stringent perturbation of promoter activity^59^. Consistent with model predictions, TS frequency decreased rapidly to near zero following TPL addition at all tested time points, indicating transcriptional shutoff in 10 minutes of less. Model predictions accurately captured the subsequent temporal evolution of nuclear and cytoplasmic *DUSP1* mRNA following transcriptional arrest (Fig. 4d), with nuclear and cytoplasmic mRNA distributions and joint nuclear–cytoplasmic distributions for each TPL timing shown in Supplementary Figure S14.

To focus directly on the model’s prediction of a temporal change in the cytoplasmic degradation rate, Figure 4d shows the predicted and measured post-TPL decay rates for pooled and normalized data before and after 75 min following Dex. At early inhibition times, the observed decay rate was 0.0140 ± 0.0036 min^*−*1^, and at later times the decay measured to be significantly higher at 0.0376 ± 0.0073 (one sided T-test, p = 0.0066). These relative decay rates were correctly estimated by the model, which predicted an effective decay rate of 0.0131 ± 0.0045 min^*−*1^ at early times and 0.0316 ± 0.0019 min^*−*1^ at later times.

Together, these results demonstrate that the inferred models for GR transport and degradation and GR-mediated transcriptional activation, and nonlinear time-varying cytoplasmic decay mechanisms are sufficient to predict temporal and spatial aspects of *DUSP1* mRNA dynamics across Dex concentrations and under acute transcriptional inhibition.

## Discussion

Here we combined single-cell ICC, smFISH, and stochastic modeling to quantitatively link GR transport with transcriptional and post-transcriptional regulation of the endogenous anti-inflammatory gene *DUSP1*. By fitting discrete stochastic models directly to joint distributions of nuclear and cytoplasmic GR, transcription site activity, and mRNA abundance, we moved beyond mean responses to resolve how variability and timing emerge from the underlying regulatory processes. This integrative framework captures not only the magnitude but also the heterogeneity and temporal ordering of GR-dependent transcriptional responses, providing a quantitative description of how GC signaling propagates from receptor activation to mRNA clearance at the single-cell level.

Our analysis supports a model in which Dex primarily drives GR nuclear accumulation through enhanced import, followed by concentration-dependent nuclear degradation that modulates *DUSP1* transcriptional output. This compartment-specific turnover provides a natural explanation for the observed dose-dependent rise and decline of nuclear GR, particularly at higher Dex concentrations where nuclear GR peaks early and decays rapidly. Such dynamics are consistent with proteasome-mediated clearance^7,54,55,60^ acting as a built-in deactivation mechanism that constrains high nuclear flux, rather than simple nuclear export or cytoplasmic recycling^23^. Importantly, this interpretation does not require assuming multiple transcriptionally distinct GR species, but instead emerges from fitting population-wide distributions under a minimal transport and degradation scheme. Future studies examining single-cell distributions of GR phosphorylation states (e.g., S211) over time could further resolve the relationship between ligand-induced activation, transcriptional competence, and proteasomal targeting^61,62^.

At the level of transcriptional regulation, our single-cell measurements indicate that GR activation increases the fraction of *DUSP1* alleles engaged in transcription without producing a strong or sustained increase in TS intensity. We systematically tested models in which GR modulates the promoter activation rate *k*_on_, the promoter de-activation rate *k*_off_, the transcription rate *k*_r_, or combinations thereof. Models acting through *k*_on_ alone or through dual regulation of *k*_on_ and *k*_off_ provided the strongest fits; however, the dual regulation models predicted increases in TS intensity that were not observed in the data, favoring the simpler *k*_on_-only mechanism. General activation models including modulation of *k*_r_ did not improve fits or predictions. Within a two-state bursting framework, this pattern is most consistent with an effective increase in promoter activation frequency, rather than a large change in transcriptional output per active allele^48,63^. Together with work using MMTV reporter constructs^12,28,54,64^, which found that GR primarily increases burst frequency, this pattern strongly supports a dominant effect of GR on the effective activation rate *k*_on_. Chromatin studies at the endogenous *DUSP1* locus further show GR-dependent recruitment of p300, increases in histone H3/H4 acetylation, and localized chromatin remodeling at the *DUSP1* GRE, features that are naturally associated with both facilitating promoter activation and stabilizing the active chromatin state^32^. While additional mechanisms such as stabilization of the active promoter state cannot be excluded, particularly given chromatin remodeling at the *DUSP1* locus, these effects are not uniquely identifiable from our current fixed-cell transcription site intensity measurements alone. Accordingly, our model represents GR action through dual regulation of the promoter, in which a GR-dependent increase in the effective activation rate *k*_on_ dominates over a smaller effect on the de-activation rate *k*_off_, capturing the dominant but not exclusive contribution of GR to transcriptional control.

Beyond transcription initiation, the results of our semi-mechanistic models highlight cytoplasmic mRNA decay as a key mechanism in the Dex-GR-*DUSP1* expression dynamics. Our models required a saturable, time-dependent degradation process to capture *DUSP1* both nuclear and cytoplasmic mRNA dynamics, particularly the rapid decline of cytoplasmic mRNA following peak expression that could not be reconciled with a fixed degradation rate given comparatively sustained transcriptional activity in the nucleus. This is consistent with Muazzen et al., who showed that Dex treatment reduces *DUSP1* mRNA half-life even as steady-state levels rise, and more broadly that GCs universally destabilize ARE-containing mRNAs like DUSP1^37^. The heterogeneity of this response has consequential implications: comparative transcriptomic analyses between Dex treatment and SARS-CoV-2 infection revealed that ARE-mRNAs tend to regulate in opposite directions^37,65^, suggesting that ARE-dependent post-transcriptional control is a clinically relevant factor in GC efficacy during COVID-19^66^.

To further our understanding of the observed cytoplasmic mRNA decline, we extended from our final semi-mechanistic model to a fully mechanistic model that explicitly incorporates Dex-induced *TTP* gene activation via nuclear GR, *TTP* mRNA production and nuclear export, TTP protein translation, and reversible binding of TTP to ARE-containing mRNAs including *DUSP1* and *TTP* itself (Supplementary Fig. S11). Although this extended model is not uniquely identifiable leading to far greater parameter uncertainties (Supplementary Figs. 15-18), it achieved stronger fits and (more importantly) substantially improved predictions for unseen experimental data (Supplementary Fig. S19, Table S6), offering an interpretable explanation for the faster clearance of cytoplasmic relative to nuclear *DUSP1* mRNA. In this framework, we have assumed that Dex-induced TTP is unphosphorylated and therefore active, since Dex does not activate p38 but instead induces *DUSP1*/MKP-1, which suppresses p38 activity and maintains TTP in its decay-promoting state^37,57,67^. Thus, ligand-induced *DUSP1* expression is shaped not only by transcriptional activation but also by an actively regulated mRNA clearance circuit that is itself a product of GR signaling.

Several limitations frame the scope of our conclusions. All measurements were obtained using ICC and smFISH measurements that allowed us to obtain endogenous expression measurements without perturbing the cells’ genetic or biochemical processes, but the reliance on fixed cells restricts direct observation of temporal correlations within individual cells and limits identifiability of closely related regulatory mechanisms. In addition, GR localization and *DUSP1* transcription were quantified in separate experiments, such that nuclear GR abundance and *DUSP1* mRNA levels were not measured simultaneously within the same cells. As a result, the coupling between GR transport and transcriptional output is inferred statistically across matched populations rather than directly observed at the single-cell level. Importantly, *TTP* mRNA and protein levels have not been measured directly in this study; TTP dynamics were inferred through model fitting and are consistent with established biochemistry^36–38^ but remain to be validated experimentally in this system. Future work should directly quantify *TTP* expression, TTP phosphorylation status, p38 activity, and MKP-1 protein levels to confirm the proposed regulatory circuit and further constrain model parameters. Our analysis further focuses on a single endogenous target gene in a single epithelial cell line, and regulatory architectures may differ across GR targets or cell types. Nevertheless, the experimental and computational framework developed here is extensible to additional genes, perturbations, and signaling context. As such, by demonstrating how integrating endogenous single-cell measurements with stochastic modeling can yield predictive, mechanistic insight into hormone-driven gene regulation, this work provides a foundation for future studies to rationally modulate inflammatory signaling through model-guided perturbations.

## Methods

### Experimental Methods

#### Cell Culture

HeLa Kyoto cells (a gift from the DeLuca Lab) were maintained in Dulbecco’s modified Eagle medium (DMEM, Thermo Fisher Scientific, 11,960–044) supplemented with 10% fetal bovine serum (FBS, Atlas Biologicals, F-0050-A), 10 U/mL penicillin/streptomycin (P/S, Thermo Fisher, 15140122), 1 mM L-glutamine (L-glut, Thermo Fisher Scientific, 25030081) in a humidified incubator at 37 °C with 5% CO_2_.

For smFISH and ICC experiments, HeLa Kyoto cells were seeded on 18 mm cover glasses in a 12-well plate at approximately 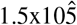 cells per well, in DMEM supplemented with 10% Charcoal Stripped fetal bovine serum (csFBS, Sigma, F6765-500ML), 10 U/mL penicillin/streptomycin (P/S, Thermo Fisher, 15140122), 1 mM L-glutamine) 24 hours prior to the experiment.

#### Drug Stimulation

Dexamethasone (Dex, Sigma, D2915) was reconstituted in nuclease free water to a stock concentration of 67.3 *×* 10^3^ nM. For experimentation, dilutions were performed such that a volume of 1.6 µL in 1000 µL had the desired concentration. All samples were incubated at 37 °C with 5% CO_2_ during the experiment and were only temporarily removed for the addition and mixing of the stimulant(s) (approx. 1.5 min).

For transcription inhibition, cells were first stimulated with 100nM Dex as above. Following Dex stimulation, 5mM of Triptolide (TPL, Sigma, 645900) was added to the corresponding wells at appropriate times. All samples were incubated at 37 °C with 5% CO_2_ during the experiment and were only temporarily removed for the addition and mixing of the stimulant(s) (aprox. 1.5 min).

#### Immunocytochemistry

Immunocytochemistry was performed using a modified protocol described here:

https://www.abcam.com/en-us/technical-resources/protocols/icc-protocol

Briefly, the samples were rinsed twice with RNase-free 1X PBS and fixed with 4% PFA at room temperature for 10 minutes. Samples were permeabilized with 0.1% Triton-X100 in PBS for 20 min and washed with 1 mL of PBS three times for ten minutes each wash. Next, samples were blocked for nonspecific binding with 10% Normal Goat Serum (Thermo Fisher, 50062Z) for 1 hr at 20 °C. Then samples were incubated overnight in a 1:500 dilution of primary antibody (Abcam, AB183127) containing 1% BSA(Thermo Fisher, A14289SA) in PBST (PBS with 0.1% Tween-20 (Thermo Fisher, 85113)). The next day the primary antibody solution was aspirated, and the samples were washed with 1 mL of PBS three times for ten minutes each wash. The samples were incubated with the secondary antibody solution 0.5 µg/mL Goat anti-Rabbit IgG (H+L) Cross-Adsorbed Secondary Antibody, Alexa Fluor 488 (Thermo Fisher, A11008), 5% Normal Goat Serum, and 50 ng/mL DAPI in PBST) in the dark for 1hr at 20 °C. Samples then were washed 4 times. Once with 1 mL of PBST for 10 min, and three times with 1 mL of 1X PBS for 10 min each wash. Finally, the samples are mounted on a 15 µL droplet of Vectashield mounting medium (Vector Laboratories, H-1000-10), and sealed with clear nail polish. (Abcam 1 ^*°*^ Ab: Rb mAb to Glucocorticoid Receptor [EPR19621] (Ab183127) [0.549 mg/mL])

#### Single-Molecule Inexpensive FISH

##### Probe Design

To design smFISH probes for *DUSP1* and GAPDH, the gene’s sequence was obtained from the Ensembl genome browser by using the transcript ID (DUSP1:ENST00000239223.4, GAPDH:ENST00000229239.10) and exported as a FASTA file. The RepeatMasker Web Server was used to identify any repetitive sequences within the *DUSP1* and GAPDH exonic sequences and the Oligostan script in R-studio is used to generate potential probes, ensuring at least 21-24 probes for high-quality results. Candidate probes were filtered based on criteria from relevant literature to select those with optimal thermodynamic properties and specificity, avoiding probes that bind non-target regions. Probe specificity was assessed by searching the sequences against genomic and transcript databases using BLAST. Probes were selected if they aligned perfectly to *DUSP1* and showed no significant similarity to other genes. The selected probe binding sites were visualized and verified using vector mapping software, and the final probes, including the FLAP sequences, were ordered from IDT at 100 µM. The final DUSP1 probe set is provided in Supplementary Figure S1.

##### Sample Labeling

smFISH was performed following a protocol previously described^56,68^. Briefly, samples were rinsed twice with RNase-free 1 *×* PBS and fixed with 4% PFA at room temperature for 10 min. This was followed by permeabilization with 70% ethanol at 4 °C for a minimum of 1 hr. Samples were washed with Wash A buffer (Biosearch Technologies, SMF-WA1-60) for 5 min. Each coverslip was placed cell-side down on a droplet containing 40 µL of hybridization buffer (Biosearch Technologies, SMF-HB1-10) and 1 µL of each duplexed smFISH probe set (*DUSP1* probes annealed to FLAP-Y-Cy5 and GAPDH exon probes annealed to FLAP-Y-Cy3), and incubated overnight at 37 °C in a sealed humidified chamber.

The following day, samples were transferred to a new 12-well plate and washed twice in Wash A buffer containing 10% formamide at 37 °C for 30 min, first without additives and then with 50 ng/mL DAPI included. Finally, samples were treated with Wash B buffer (Biosearch Technologies, SMF-WB1-20) at room temperature for 5 min, mounted on a 15 µL droplet of Vectashield mounting medium (Vector Laboratories, H-1000-10), and sealed with clear nail polish.

#### Microscopy Imaging

Fluorescent images were acquired on an Olympus IX81 inverted spinning-disk confocal microscope (CSU22 head with quad dichroic and an additional emission filter wheel to minimize spectral crossover) using a 60*×*/1.42 NA oil-immersion objective. Confocal z-stacks were collected with a step size of 0.5 µm, with 27 optical sections acquired per channel.

ICC experiments were imaged using two high-power diode lasers with rapid microsecond switching (405 nm for DAPI, 100 ms exposure; 488 nm for Alexa Fluor 488, 100 ms exposure; Thermo Fisher, A11008). smFISH fields of view were imaged using three high-power diode lasers with rapid microsecond switching (405 nm for DAPI, 100 ms; 561 nm for smFISH of GAPDH exons, 100 ms; and 647 nm for smFISH of *DUSP1* mRNA, 300 ms). Combined smFISH/ICC fields of view were imaged using four high-power diode lasers (405 nm for DAPI, 100 ms; 488 nm for Alexa Fluor 488, 100 ms; 561 nm for smFISH of GAPDH exons, 100 ms; and 647 nm for smFISH of *DUSP1* mRNA, 300 ms).

The system was equipped with differential interference contrast (DIC) optics, built-in spherical aberration correction for all objectives, and a wide-field xenon light source. Images were acquired using an EMCCD camera (iXon Ultra 888, Andor) controlled with SlideBook software, yielding images with an effective pixel size of 160 nm. The imaging area was 624 *×* 928 pixels.

### GR ICC Image Processing and Quantification

To obtain accurate and consistent subcellular single-cell GR measurements, raw image stacks were processed through the following steps: illumination correction, pseudo-cytoplasmic mask generation, intensity normalization by imaging date, intensity discretization, and conversion to concentrations using per-cell nuclear-to-cytoplasmic area ratios. Although images were acquired as z-stacks of shape [*P, T,C, Z,Y, X* ], all subsequent intensity quantification and mask generation were performed on 2D maximum-intensity projections along the Z-axis.

#### Illumination Correction

Illumination correction was performed using a data-driven, background-based approach. Image z-stacks were first projected along the Z-axis using the 10th percentile intensity to generate a background estimate per field of view, then aggregated across fields of view via a median projection to produce a single profile per channel. Each profile was smoothed with a Gaussian filter (channel-specific *σ*) to suppress noise and remove cellular morphology artifacts. A 2D Gaussian was then fit to each smoothed profile using nonlinear least squares to produce a parametric correction map. Each fitted profile was normalized to its maximum value and inverted to form a per-pixel correction factor, which was applied multiplicatively to the original image stack. Corrected images were re-projected and visualized to confirm uniformity (Supplementary Fig. S3).

#### Pseudo-Cytoplasmic Mask from DAPI Dilation

In datasets without a cytoplasmic stain, pseudo-cytoplasmic regions were derived from the nuclear (DAPI) mask. Nuclear masks were dilated by 20 pixels, and a watershed algorithm was applied to prevent overlaps between neighboring cells. The resulting cell territories were matched to their corresponding nuclei, and the pseudo-cytoplasmic mask for each cell was defined as the difference between the dilated territory and the nuclear mask. Mean intensity within this pseudo-cytoplasmic ring was calculated for each cell, providing non-overlapping cytoplasmic measurements. Cells touching the image border were excluded from analysis.

#### Intensity Normalization and Cell Size Gating

Raw GR nuclear `(nucGRint)` and cytoplasmic `(cytGRint)` intensities were normalized for between-day imaging variability by a replica-scaling procedure, yielding illumination- and day-corrected intensities `(NucGRCorrected, CytGRCorrected)`. To reduce extrinsic variability due to variations in cell size, both GR and *DUSP1* datasets were gated to include only cells with nuclear areas between the 25th and 75th percentiles of their respective nuclear area distributions. The *DUSP1* datasets were then subjected to a second gate corresponding to the 25th to 75th percentile for cytoplasm areas.

#### Conversion between Units of Fluorescence Intensity and Sub-Cellular GR Mass

Mean fluorescent intensities within the nuclear and pseudo-cytoplasmic masks, `NucGRCorrected`, `CytGRCorrected`, were assumed to be proportional to the compartmental GR concentrations in arbitrary units of concentration, AUC. To determine GR levels in consistent units of mass, a nuclear-to-cytoplasmic ratio (NCR) was computed as the mean nuclear area divided by the mean cytoplasmic area in the gated *DUSP1* dataset. Arbitrary units of mass (AUM) in the nucleus was estimated as the `NucGRCorrected` intensity rescaled by the NCR, and the `CytGRCorrected` intensity was used directly as a measurement of mass in the cytoplasm. For comparison to the discrete-valued GR transport model, the resulting quantities of mass were discretized into 30 uniformly-spaced bins spanning the 1st to 99th percentile range of the `CytGRCorrected` intensity. Bins were unbounded above to avoid clipping high-expressing cells, and the same bin definitions were used for both nucleus and cytoplasm. The resulting `normGRnuc` and `normGRcyt` represent sub-cellular GR concentrations in arbitrary units of mass (AUM).

### *DUSP1* smFISH mRNA Quantification

All image processing codes for smFISH mRNA quantification can be found at: https://github.com/MunskyGroup/Ron_et_al_2026_ImageProcessing

#### Data Conversion

Raw microscopy image stacks and segmentation masks were converted into a standardized HDF5 format for downstream processing. For each experimental folder, multi-channel 3D TIFF images and associated log files were parsed to identify fields of view, time points, and channels. Masks for nuclei and cytoplasm, if precomputed, were unpacked and aligned to their respective channels. All image data were stored as `float32` Dask arrays, rechunked for efficient processing, and saved to disk along with acquisition metadata in JSON format embedded within the HDF5 file.

#### Cellpose Segmentation

Segmentation of nuclei and cell boundaries was performed using the Cellpose^69^ deep learning framework, with custom pretrained models for nuclear DAPI and cytoplasmic smFISH GAPDH-Cy3 channels^69^. Nuclear masks were generated from the DAPI channel, while cytoplasmic masks were segmented from the corresponding GAPDH smFISH channel (3a). Model parameters (diameter, flow threshold, probability threshold, and minimum object size) were optimized for each sub-cellular compartment. Segmentation results were post-processed to ensure one-to-one alignment between nuclear and cytoplasmic masks (Supplementary Fig. S2).

#### Preliminary Spot Detection

Initial RNA spot detection was performed using BigFISH^34^, operating on the *DUSP1*-Cy5 smFISH channel. Detected coordinates were annotated with per-spot properties including voxel-based size, signal intensity, and a computed signal-to-noise ratio (SNR) using the `compute_snr_spots` function. For each field of view, spot coordinates were mapped to individual cells and classified as nuclear or cytoplasmic using segmentation masks. Clusters corresponding to transcription sites or large cytoplasmic foci were identified and later included in single-molecule counts (Supplementary Fig. S2).

#### BigFISH Threshold Selection

For each smFISH channel, an empirical detection threshold was selected by analyzing a random subset of up to 50 images. Spot intensity distributions were used to compute descriptive statistics (minimum, maximum, median, mean, and quartiles), from which a per-dataset threshold was chosen. In most datasets, the mean threshold value was applied; for specific noisier datasets, fixed percentile values (25th, 75th, or 90th) were used instead to improve robustness.

#### SNR Analysis and Thresholding

Spot quality control was based on two complementary criteria:

- **Absolute SNR**: A hard cutoff requiring SNR *≥* `abs_threshold`.
- **MG SNR**: A compartment-specific SNR defined as

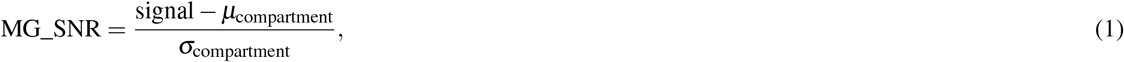

where *µ*_compartment_ and *σ*_compartment_ are the mean and standard deviation of intensity within the nuclear or cytoplasmic mask of the same cell.

For all analyses, the `mg_abs` method was used, retaining spots that passed either the MG SNR threshold (`mg_threshold`) = 3 or the absolute SNR threshold (`abs_threshold` = 6). Spots failing both criteria were excluded from downstream quantification (Supplementary Fig. S2).

#### Cell-Level Quantification

Raw spot, cluster, and cell-property data were loaded and concatenated across all imaging slides for each experimental replicate using a combined dataset manager. SNR metrics were computed per spot via `SNRAnalysis` (`abs_threshold` = 6, `mg_threshold = 3`), annotating each spot with an absolute SNR flag and a compartment-specific MG SNR value (Equation 1).

Spot filtering used the `mg_abs` method, which retains a spot if it passes the MG SNR threshold `(MG_pass)` or, for spots with 1 *≤* MG_SNR *≤* `mg_threshold`, if their absolute SNR exceeds `abs_threshold`. This fallback prevents rejection of real spots in cells with highly variable background. Spots failing both criteria were recorded separately for quality review. Cells touching image borders were excluded at the filtering stage. To ensure consistent cell identity across replicates, unique cell, spot, and cluster identifiers were offset by a replicate-specific numeric prefix prior to concatenation.

Using the filtered spot list, the `DUSP1Measurement` pipeline aggregated single-molecule counts to the cell level across two quality filters: absolute (SNR *≥* `abs_threshold)`, and `MG_SNR` (compartment-specific SNR *≥* `mg_threshold)`. Each filter yielded total, nuclear, and cytoplasmic spot counts per cell.

TS and cytoplasmic foci were identified from nuclear and cytoplasmic clusters, respectively, subject to a minimum cluster size of four spots. Clusters falling below this threshold were excluded from TS and foci counts but tracked via diagnostic fields `(n_ts_lt_min, n_foci_lt_min)`. For valid TS, the largest and second-largest cluster sizes were also recorded `(largest_ts, second_largest_ts)`.

Additional per-cell metrics included:

- nuclear and cytoplasmic area,
- mean and standard deviation of nuclear, cytoplasmic, and whole-cell intensities, and
- experimental metadata (time, Dex concentration, replicate, FOV).

This pipeline was applied identically to all *DUSP1* experimental replicates. Cell-level results were saved alongside the filtered spot, cluster, and cell property tables for downstream analysis.

### Stochastic Modeling

We used the Stochastic System Identification Toolkit (SSIT) (https://github.com/MunskyGroup/SSIT/ ^53^) to define, solve, and fit the GR-mediated *DUSP1* transcription (Dex-GR-*DUSP1*) model. Scripts to reproduce all analyses and generate all figures are found at [https://github.com/MunskyGroup/Ron_et_al_2026], and are briefly described below.

The full model consists of six species: inactive gene copies (‘offGene’), active gene copies (‘onGene’), cytoplasmic GR (‘cytGR’), nuclear GR (‘nucGR’), nuclear mRNA transcripts (‘rNuc’), and cytoplasmic mRNA (‘rCyt’). The precise details of these reactions vary depending on the specific model and the three stages of the identification procedure:

#### GR Model

The model of GR transport and decay consists of five reactions: GR production in the cytoplasm (**reaction 1**: *→* ∅ cytGR with rate *k*_GR_), GR degradation in the cytoplasm (**reaction 2**: cytGR *→* ∅ with rate *γ*_GRcyt_), GR transport from cytoplasm to nucleus (**reaction 3**: cytGR *→* nucGR with rate *k*_cn_), GR transport from nucleus to cytoplasm (**reaction 4**: nucGR *→* cytGR with rate *k*_nc_), and GR degradation in the nucleus (**reaction 5**: nucGR *→* ∅ with rate *γ*_GCnuc_). The influence of Dex on GR was modeled as a time-varying deterministic signal:

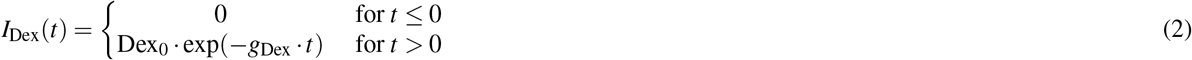

Dex controlled either import 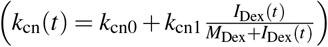 or export 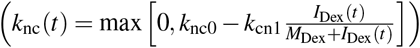, depend-ing on the model. Four alternative models were considered (Fig. 2b, Supplementary Fig S5) with different combinations of degradation locations (i.e., with or without reactions 2 or 5) and different mechanisms of transport control (i.e., variation in either *k*_nc_ or *k*_cn_).

#### *DUSP1* Model

Nascent and nuclear mRNA were assumed to be controlled by the upstream GR process and four additional reactions: gene activation (**reaction 6**: offGene *→* onGene with rate *k*_on_), gene deactivation (**reaction 7**: onGene *→* offGene with rate *k*_off_), transcription initiation (**reaction 8**: (offGene,onGene) *→* (offGene,onGene) + rNuc with rate *k*_r*−*OFF_ or *k*_r*−*ON_, depending on the state of the gene), and transport to the cytoplasm (**reaction 9**: rNuc *→* rCyt with rate *k*_transport_).

Three mechanisms were considered for nuclear GR to control DUSP1 mRNA expression (Supplementary Fig S9): by increasing activation 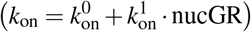, by decreasing deactivation (*k*_off_ = *k*_off_*/*(1 + *m*_koff_ *·* nucGR)), or by increasing transcription initiation in the ON state 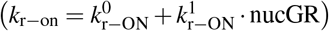, Transcription elongation was assumed to occur over a fixed deterministic time following initiation, *τ*_elong_, which defined the accumulation time for nascent mRNA on the TS and the time delay between the GR input and the release of mature mRNA into the nucleus (i.e., mature mRNA release at time *t* is assumed to be controlled by nucGR at time *t −τ*_elong_).

Cytoplasmic mRNA were assumed to be degraded (**reaction 10**: rCyt *→* ∅ with rate *γ*). Four alternate models were proposed for the control of this degradation (Supplementary Fig S10): In the *first order decay* model, *γ* was assumed to be constant (*γ* = *γ*_0_). In the *time variation only* model, *γ* was assumed to vary according to *γ*(*t*) = *γ*_0_ *· I*_TTP_(*t*), where

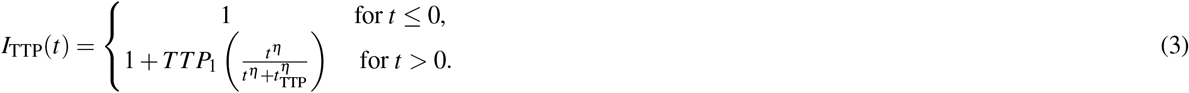

In the *saturation only* model, the rate of decay was modeled assuming biphasic decay rates according to *γ*(*r*_Cyt_) = *γ*_0_ *· r*_Cyt*−*F_ + *γ*_1_ *r*_Cyt*−*B_, where the fraction of total cytoplasmic mRNA in the free state (*r*_Cyt*−*F_) and bound state (*r*_Cyt*−*B_) were determined by rapid binding and unbinding between *r*_Cyt_ and a finite amount of *TT P*_tot_ (see supplemental text for full derivation). In the combined *time variation and saturation* model, the level of *TT P*_tot_ evolves according to Eq. 3.

TPL perturbations were modeled by assuming that the transcription initiation rates (*k*_r*−*ON_ and _r*−*OFF_) decreased from their original value to zero according to linear exponential decay (*k*_r*−*X_(*t*) = *k*_r*−*X_ exp(*− k*_TPL_ (*t· − t*_TPL_))), where ‘X’ is replaced with ‘ON’ or ‘OFF’ as needed.

##### Chemical Master Equation Solutions

Solutions to the time varying Chemical Master Equation were obtained using either Finite State Projection analyses^52^ or Stochastic Simulations^70^ using the SSIT.^53^ For comparison to GR data, the model for nuclear and cytoplasmic GR was solved using the FSP. For preliminary calculations and comparisons to nascent and nuclear DUSP1 mRNA distribution data, an approximate hybrid analysis was used where nucGR and cytGR were analyzed using ODEs that fed into an FSP model for the gene states and mRNA distributions. All ODE and FSP analyses assumed probability distributions corresponding to steady state under *Dex* = 0*nM* at time *t* = 0. SSA analyses were initiated at the steady state ODE mean (rounded to the nearest integer) at a time *t* = *−* 500 min and allowed to evolve to approximate steady state at *t* = 0. The hybrid ODE-FSP approximation was verified to match reasonably well to SSA simulations for nuclear mRNA (p = 0.07, median KS test, for 1000 simulations, Supplemental Fig. S20), allowing for an efficient mid-fidelity means to constrain parameter guesses. However, because deviations were observed at the tails due to ignoring variability in the GR signal, final decisions on model selection were made based on SSA analysis of the full model.

##### Parameter Inference and Prediction Quantification

After solving the chemical master equation using FSP or hybrid ODE–FSP approximations, the probability distributions were used to estimate model parameters using likelihood functions and sampling of their Bayesian posterior distributions. For a model (*M*) with parameter vector (*θ*), the solution gives the probability (*P*_*M*_(**x**|*t, θ*)) of each discrete molecular state (**x**) at time (*t*). Experimental observations were compared to the corresponding model-predicted distributions for the measured species. Thus, for cells indexed by (*c*) at time points indexed by (*t* _*j*_), the likelihood of the collected data points *D ≡ {***d**_c,j_*}* was computed as the product of the model probabilities assigned to the observed counts,

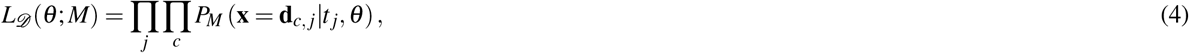

or equivalently as the summed log-likelihood,

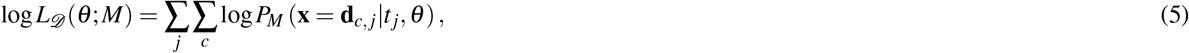

In the model inference, priors (*P*(*θ*))were assumed to compute the log posterior as:

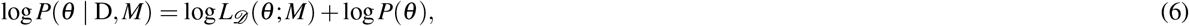

where independent Gaussian priors were placed on log_10_-transformed parameters.

For the full model in the final estimation phase, direct likelihood maximization using the FSP was not feasible, and approximate loss factors were used. For each parameter guess, a set of 10,000 independent SSA trajectories for *r*_Cyt_ and *r*_Nuc_ and a corresponding FSP analyses for nascent RNA were combined to define the loss function

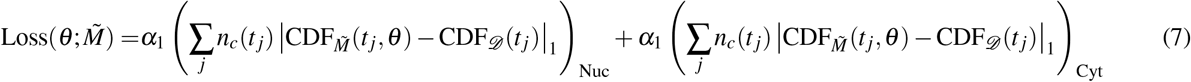

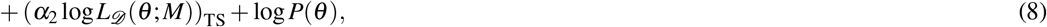

where 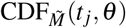 and CDF_*D*_ (*t* _*j*_) are the cumulative density functions of model-generated simulations and the data at the time *t* _*j*_, and *n*_*c*_(*t* _*j*_) is the number of cells measured at each time. To combine metrics for different model analyses and data sets, the heuristic factors *α*_1_ and *α*_2_ were chosen to achieve consistent variation in the goodness of fit for nuclear distributions. Specifically, *α*_1_, which scales the first two terms, was defined as

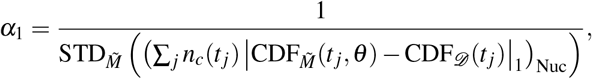

and *α*_2_, which scales the third term, was defined as

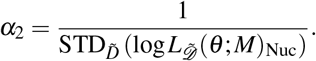

Here, the first STD is taken over independent sets of SSA trajectories for the model 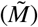 while the real data is fixed, and the second STD is taken over independent samples of simulated data 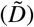 while the deterministic FSP analysis for *M* is fixed. For consistency, definition of these parameters considers only the nuclear DUSP1 mRNA distributions, analyzed using SSA for *α*_1_ and using the hybrid ODE-FSP for *α*_2_.

Parameter estimation was performed in four stages. First, the eight parameters involved in GR translocation, creation, and degradation in the nucleus and cytoplasm were identified using full FSP-based maximum a posterior estimation (MAP, Eq. 6) compared to GR measurements at 1, 10, and 100 nM Dex. Next, preliminary values for the nine nuclear *DUSP1* parameters related to promoter switching, transcription, and mRNA transport were estimated using hybrid ODE analysis of GR and FSP analysis of Nascent and Nuclear DUSP1 and estimated using MAP estimation from measurements at 100 nM Dex. Third, all but the GR parameters were updated using 10,000 SSA trajectories per parameter set for approximate Bayesian computing on the marginal distributions of nascent, nuclear, and cytoplasmic DUSP1 mRNA using measurements at 100 nM Dex. Finally, the TPL inhibition rate *k*_TPL_ was estimated using the nuclear mRNA distributions at 0 nM Dex following application of TPL.

### Other metrics for model comparison

When likelihood calculations were possible, model complexity was summarized using Akaike and Bayesian Information Criteria,

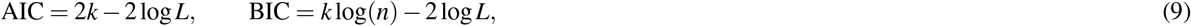

where *k* is the number of fitted parameters, and *n* is the number of single-cell observations used in the comparison. For models scored using MAP values rather than pure log likelihoods, the same expressions were used as posterior-score-based information criteria, and these values were interpreted as relative model-comparison summaries rather than exact frequentist likelihood criteria. Differences in AIC and BIC were computed relative to the best-scoring model, with lower values indicating better support after penalizing the number of parameters. We also reported approximate Bayes factors (BF) derived from BIC differences,

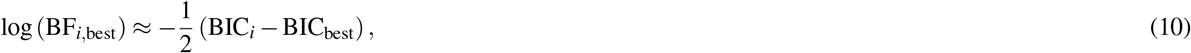

so that values less than zero indicate weaker support for model (i) relative to the best-BIC model. These likelihood, posterior-score, AIC, BIC, and approximate log Bayes factor summaries were tabulated for the GR and nuclear *DUSP1* model comparisons in Supplementary Tables S2,S4.

### Model predictions

Model predictions for unseen concentrations of Dex or TPL repression experiments at different times were evaluated without refitting. However, to account for the secondary effect of Dex on the time varying signal *I*_TTP_(*t*) in the semi-mechanistic models, the maximum value of the TTP signal (*TT P*_1_) was scaled by the relative area under the nucGR curve compared to that at 100 nM dex:

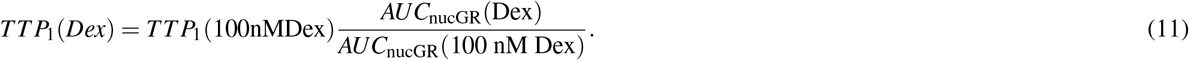

Predictions using the fully mechanistic model used exclusively the parameters fit at 100 nM Dex without any such adjustments. Prediction performance is quantified as likelihood of new data sets given the previously identified parameters (when exclusively using the FSP or hybrid ODE-FSP analyses) or by the ABC objective function (when using SSA simulations). In this latter case, prediction scores were used as simulation-based measures of generalization rather than formal likelihood-ratio statistics. For TPL predictions, only previously unseen post-treatment time points were considered to evaluate prediction performance.

## Acknowledgements

We thank all past and current members of the Munsky Group, members of the Stasevich Lab, and Gregor Neuert for helpful discussions, suggestions, and microscope time, and the de’Luca Lab for providing the HeLa cells. All main and Supplementary figures were created or compiled using BioRender (https://BioRender.com).

## Author contributions statement

Conceptualization: E.R., B.M.

simFISH Probe design: L.F.S.Q.

Experimental Procedure design: E.R., L.F.S.Q.

Performed experiments/collected data: E.R.

Designed analysis software: J.F., E.R., L.U.A., B.M.

Analyzed results: E.R., B.M.

Computational Modeling: B.M., A.P., E.R.

Wrote the original draft: E.R., B.M., A.P.

All authors reviewed the manuscript

Resources, supervision, and funding acquisition: B.M.

## Additional information

Image quantification codes: https://github.com/MunskyGroup/AngelFISH

SSIT specific analyses GR-*DUSP1*: https://github.com/MunskyGroup/Ron_et_al_2026

## Data availability

The processed data files for GR and *DUSP1* used in this study are available in the SSIT repository. Full data, including the raw microscopy images, are available from the corresponding author upon request.

## References

1. Barnes, P. J. Glucocorticosteroids: current and future directions. Br. J. Pharmacol. 163, 29–43, DOI: 10.1111/j.1476-5381.2010.01199.x (2011). https://bpspubs.onlinelibrary.wiley.com/doi/pdf/10.1111/j.1476-5381.2010.01199.x.

2. Schäcke, H., Döcke, W. D. & Asadullah, K. Mechanisms involved in the side effects of glucocorticoids. Pharmacol. & Ther. 96, 23–43, DOI: 10.1016/S0163-7258(02)00297-8 (2002).

3. Newton, R., Shah, S., Altonsy, M. O. & Gerber, A. N. Glucocorticoid and cytokine crosstalk: Feedback, feedforward, and co-regulatory interactions determine repression or resistance. J. Biol. Chem. 292, 7163–7172, DOI: 10.1074/JBC.R117.777318 (2017).

4. Zhao, Y., Yao, Z., Xu, S., Yao, L. & Yu, Z. Glucocorticoid therapy for acute respiratory distress syndrome: Current concepts. J. Intensive Medicine DOI: 10.1016/j.jointm.2024.02.002 (2024).

5. Lasa, M., Abraham, S. M., Boucheron, C., Saklatvala, J. & Clark, A. R. Dexamethasone Causes Sustained Expression of Mitogen-Activated Protein Kinase (MAPK) Phosphatase 1 and Phosphatase-Mediated Inhibition of MAPK p38. Mol. Cell. Biol. 22, 7802–7811, DOI: 10.1128/MCB.22.22.7802-7811.2002 (2002).

6. Lyapun, I. N., Andryukov, B. G. & Bynina, M. P. HeLa Cell Culture: Immortal Heritage of Henrietta Lacks. Mol. Genet. Microbiol. Virol. 34, 195–200, DOI: 10.3103/S0891416819040050 (2019).

7. Wallace, A. D. & Cidlowski, J. A. Proteasome-mediated Glucocorticoid Receptor Degradation Restricts Transcriptional Signaling by Glucocorticoids. J. Biol. Chem. 276, 42714–42721, DOI: 10.1074/jbc.M106033200 (2001).

8. Hoppstädter, J. & Ammit, A. J. Role of dual-specificity phosphatase 1 in glucocorticoid-driven antiinflammatory responses. Front. Immunol. 10, DOI: 10.3389/FIMMU.2019.01446 (2019).

9. Clark, A. R. & Dean, J. L. The control of inflammation via the phosphorylation and dephosphorylation of tristetraprolin: a tale of two phosphatases. Biochem. Soc. Transactions 44, 1321–1337, DOI: 10.1042/bst20160166 (2016).

10. Vettorazzi, S., Nalbantoglu, D., Gebhardt, J. C. M. & Tuckermann, J. A guide to changing paradigms of glucocorticoid receptor function—a model system for genome regulation and physiology. The FEBS J. 289, 5718–5743, DOI: 10.1111/FEBS.16100 (2022).

11. Timmermans, S., Souffriau, J. & Libert, C. A general introduction to glucocorticoid biology. Front. Immunol. 10, DOI: 10.3389/FIMMU.2019.01545 (2019).

12. Stavreva, D. A. et al. Transcriptional Bursting and Co-bursting Regulation by Steroid Hormone Release Pattern and Transcription Factor Mobility. Mol. cell 75, 1161–1177.e11, DOI: 10.1016/j.molcel.2019.06.042 (2019).

13. Garcia, D. A. et al. Power-law behavior of transcription factor dynamics at the single-molecule level implies a continuum affinity model. Nucleic Acids Res. 49, 6605–6620, DOI: 10.1093/nar/gkab072 (2021).

14. Garcia, D. A. et al. An intrinsically disordered region-mediated confinement state contributes to the dynamics and function of transcription factors. Mol. cell 81, 1484–1498.e6, DOI: 10.1016/j.molcel.2021.01.013 (2021).

15. Robertson, S., Hapgood, J. P. & Louw, A. Glucocorticoid receptor concentration and the ability to dimerize influence nuclear translocation and distribution. Steroids 78, 182–194, DOI: 10.1016/J.STEROIDS.2012.10.016 (2013).

16. Vandevyver, S., Dejager, L. & Libert, C. On the Trail of the Glucocorticoid Receptor: Into the Nucleus and Back. Traffic 13, 364–374, DOI: 10.1111/J.1600-0854.2011.01288.X (2012).

17. Haché, R. J. G., Tse, R., Reich, T., Savory, J. G. A. & Lefebvre, Y. A. Nucleocytoplasmic Trafficking of Steroid-free Glucocorticoid Receptor *. J. Biol. Chem. 274, 1432–1439, DOI: 10.1074/jbc.274.3.1432 (1999).

18. Vandevyver, S., Dejager, L. & Libert, C. Comprehensive overview of the structure and regulation of the glucocorticoid receptor. Endocr. Rev. 35, 671–693, DOI: 10.1210/er.2014-1010 (2014).

19. Wagh, K., Stavreva, D. A., Upadhyaya, A. & Hager, G. L. Transcription Factor Dynamics: One Molecule at a Time. Annu. Rev. Cell Dev. Biol. 39, 277–305, DOI: 10.1146/annurev-cellbio-022823-013847 (2023).

20. Iyer-Bierhoff, A. et al. Acetylation-induced proteasomal degradation of the activated glucocorticoid receptor limits hormonal signaling. iScience 27, 108943, DOI: 10.1016/j.isci.2024.108943 (2024).

21. Itoh, M. et al. Nuclear export of glucocorticoid receptor is enhanced by c-Jun N-terminal kinase-mediated phosphorylation. Mol. Endocrinol. 16, 2382–2392, DOI: 10.1210/ME.2002-0144 (2002).

22. Kinyamu, H. K., Chen, J. & Archer, T. K. Linking the ubiquitin–proteasome pathway to chromatin remodeling/modification by nuclear receptors. J. Mol. Endocrinol. 34, 281–297, DOI: 10.1677/jme.1.01680 (2005).

23. Chen, G., Lv, S., Pascal, L. E. & Wang, Z. Regulation of glucocorticoid receptor nuclear localization in prostate cancer cells. The J. Pharmacol. Exp. Ther. 392, 103577, DOI: 10.1016/j.jpet.2025.103577 (2025).

24. Lonard, D. M., Nawaz, Z., Smith, C. L. & O’Malley, B. W. The 26S Proteasome Is Required for Estrogen Receptor- and Coactivator Turnover and for Efficient Estrogen Receptor-Transactivation. Mol. Cell 5, 939–948, DOI: 10.1016/S1097-2765(00)80259-2 (2000).

25. Reid, G. et al. Cyclic, Proteasome-Mediated Turnover of Unliganded and Liganded ER on Responsive Promoters Is an Integral Feature of Estrogen Signaling. Mol. Cell 11, 695–707, DOI: 10.1016/S1097-2765(03)00090-X (2003).

26. Lv, S. et al. Regulation and targeting of androgen receptor nuclear localization in castration-resistant prostate cancer. J. Clin. Investig. 131, e141335, DOI: 10.1172/JCI141335 (2021).

27. National Center for Biotechnology Information. DUSP1 dual specificity phosphatase 1 [Homo sapiens] (Gene ID: 1843).

28. Larson, D. R. et al. Direct observation of frequency modulated transcription in single cells using light activation. eLife 2, e00750, DOI: 10.7554/eLife.00750 (2013).

29. Fryer, C. J. & Archer, T. K. Chromatin remodelling by the glucocorticoid receptor requires the BRG1 complex. Nature 393, 88–91, DOI: 10.1038/30032 (1998).

30. Nagaich, A. K., Walker, D. A., Wolford, R. & Hager, G. L. Rapid Periodic Binding and Displacement of the Glucocorticoid Receptor during Chromatin Remodeling. Mol. Cell 14, 163–174, DOI: 10.1016/S1097-2765(04)00178-9 (2004).

31. Tchen, C. R. et al. Glucocorticoid Regulation of Mouse and Human Dual Specificity Phosphatase 1 (DUSP1) Genes. The J. Biol. Chem. 285, 2642–2652, DOI: 10.1074/jbc.M109.037309 (2010).

32. Shipp, L. E. et al. Transcriptional Regulation of Human Dual Specificity Protein Phosphatase 1 (DUSP1) Gene by Glucocorticoids. PLOS ONE 5, e13754, DOI: 10.1371/JOURNAL.PONE.0013754 (2010).

33. John, S. et al. Chromatin accessibility pre-determines glucocorticoid receptor binding patterns. Nat. Genet. 43, 264–268, DOI: 10.1038/ng.759 (2011).

34. Imbert, A. et al. FISH-quant v2: a scalable and modular tool for smFISH image analysis. RNA 28, 786–795, DOI: 10.1261/RNA.079073.121/-/DC1 (2022).

35. Khabar, K. S. Post-transcriptional control during chronic inflammation and cancer: A focus on AU-rich elements. Cell. Mol. Life Sci. 67, 2937–2955, DOI: 10.1007/S00018-010-0383-X/TABLES/3 (2010).

36. Brooks, S. A. & Blackshear, P. J. Tristetraprolin (TTP): Interactions with mRNA and proteins, and current thoughts on mechanisms of action. Biochimica et Biophys. Acta (BBA) - Gene Regul. Mech. 1829, 666–679, DOI: 10.1016/j.bbagrm.2013.02.003 (2013).

37. Muazzen, Z. et al. Global analysis of the abundance of AU-rich mRNAs in response to glucocorticoid treatment. Sci. Reports 14, DOI: 10.1038/s41598-024-51301-6 (2024).

38. Tiedje, C. et al. The RNA-binding protein TTP is a global post-transcriptional regulator of feedback control in inflammation. Nucleic Acids Res. 44, 7418–7440, DOI: 10.1093/nar/gkw474 (2016).

39. Monczor, F. et al. A Model of Glucocorticoid Receptor Interaction With Coregulators Predicts Transcriptional Regulation of Target Genes. Front. Pharmacol. 10, DOI: 10.3389/fphar.2019.00214 (2019).

40. Wei-Chen Chen, D., Lynch, J. T., Demonacos, C., Krstic-Demonacos, M. & Schwartz, J. M. Quantitative analysis and modeling of glucocorticoid-controlled gene expression. Pharmacogenomics 11, 1545–1560, DOI: 10.2217/PGS.10.125 (2010).

41. Van Moortel, L. et al. Novel assays monitoring direct glucocorticoid receptor protein activity exhibit high predictive power for ligand activity on endogenous gene targets. Biomed. & Pharmacother. 152, 113218, DOI: 10.1016/J.BIOPHA.2022.113218 (2022).

42. Bakker, E. et al. Insight into glucocorticoid receptor signalling through interactome model analysis. PLOS Comput. Biol. 13, e1005825, DOI: 10.1371/journal.pcbi.1005825 (2017).

43. Munsky, B., Neuert, G. & van Oudenaarden, A. Using gene expression noise to understand gene regulation. Science 336, 183–187, DOI: 10.1126/science.1216379 (2012).

44. Neuert, G. et al. Systematic identification of signal-activated stochastic gene regulation. Science 339, 584–587, DOI: 10.1126/science.1231456 (2013).

45. Senecal, A. et al. Transcription factors modulate c-Fos transcriptional bursts. Cell Reports 8, 75–83, DOI: 10.1016/j.celrep.2014.05.053 (2014).

46. Munsky, B., Fox, Z. & Neuert, G. Integrating single-molecule experiments and discrete stochastic models to understand heterogeneous gene transcription dynamics. Methods 85, 12–21, DOI: 10.1016/j.ymeth.2015.06.009 (2015).

47. Munsky, B., Li, G., Fox, Z. R., Shepherd, D. P. & Neuert, G. Distribution shapes govern the discovery of predictive models for gene regulation. Proc. Natl. Acad. Sci. 115, 7533–7538, DOI: 10.1073/pnas.1804060115 (2018).

48. Raj, A., Peskin, C. S., Tranchina, D., Vargas, D. Y. & Tyagi, S. Stochastic mRNA synthesis in mammalian cells. PLoS Biol. 4, 1707–1719, DOI: 10.1371/JOURNAL.PBIO.0040309 (2006).

49. Hansen, M. M., Desai, R. V., Simpson, M. L. & Weinberger, L. S. Cytoplasmic amplification of transcriptional noise generates substantial cell-to-cell variability. Cell systems 7, 384–397 (2018).

50. Matthews, L. et al. Cell Cycle Phase Regulates Glucocorticoid Receptor Function. PLOS ONE 6, e22289, DOI: 10.1371/journal.pone.0022289 (2011).

51. Oakley, R. H., Webster, J. C., Sar, M., Parker, C. R., Jr. & Cidlowski, J. A. Expression and Subcellular Distribution of the -Isoform of the Human Glucocorticoid Receptor*. Endocrinology 138, 5028–5038, DOI: 10.1210/endo.138.11.5501 (1997).

52. Munsky, B. & Khammash, M. The finite state projection algorithm for the solution of the chemical master equation. J. Chem. Phys. 124, 44104, DOI: 10.1063/1.2145882/561868 (2006).

53. Popinga, A. N., Forman, J., Svetlov, D., Vo, H. & Munsky, B. The stochastic system identification toolkit (ssit) to model, fit, predict, and design experiments. bioRxiv DOI: 10.64898/2026.02.20.707039 (2026). https://www.biorxiv.org/content/early/2026/02/22/2026.02.20.707039.full.pdf.

54. Stavreva, D. A., Müller, W. G., Hager, G. L., Smith, C. L. & McNally, J. G. Rapid Glucocorticoid Receptor Exchange at a Promoter Is Coupled to Transcription and Regulated by Chaperones and Proteasomes. Mol. Cell. Biol. 24, 2682–2697, DOI: 10.1128/MCB.24.7.2682-2697.2004 (2004).

55. Wallace, A. D., Cao, Y., Chandramouleeswaran, S. & Cidlowski, J. A. Lysine 419 Targets Human Glucocorticoid Receptor for Proteasomal Degradation. Steroids 75, 1016–1023, DOI: 10.1016/j.steroids.2010.06.015 (2010).

56. Tsanov, N. et al. smiFISH and FISH-quant – a flexible single RNA detection approach with super-resolution capability. Nucleic Acids Res. 44, e165, DOI: 10.1093/NAR/GKW784 (2016).

57. Smoak, K. & Cidlowski, J. A. Glucocorticoids Regulate Tristetraprolin Synthesis and Posttranscriptionally Regulate Tumor Necrosis Factor Alpha Inflammatory Signaling. Mol. Cell. Biol. 26, 9126–9135, DOI: 10.1128/MCB.00679-06 (2006).

58. Ishmael, F. T. et al. Role of the RNA-Binding Protein Tristetraprolin in Glucocorticoid-Mediated Gene Regulation. The J. Immunol. 180, 8342–8353, DOI: 10.4049/jimmunol.180.12.8342 (2008).

59. Titov, D. V. et al. XPB, a subunit of TFIIH, is a target of the natural product triptolide. Nat. Chem. Biol. 7, 182–188, DOI: 10.1038/nchembio.522 (2011).

60. Deroo, B. J. et al. Proteasomal Inhibition Enhances Glucocorticoid Receptor Transactivation and Alters Its Subnuclear Trafficking. Mol. Cell. Biol. 22, 4113–4123, DOI: 10.1128/MCB.22.12.4113-4123.2002 (2002).

61. Avenant, C., Ronacher, K., Stubsrud, E., Louw, A. & Hapgood, J. P. Role of ligand-dependent GR phosphorylation and half-life in determination of ligand-specific transcriptional activity. Mol. Cell. Endocrinol. 327, 72–88, DOI: 10.1016/j.mce.2010.06.007 (2010).

62. Galliher-Beckley, A. J. & Cidlowski, J. A. Emerging roles of glucocorticoid receptor phosphorylation in modulating glucocorticoid hormone action in health and disease. IUBMB Life 61, 979–986, DOI: 10.1002/IUB.245 (2009).

63. Hebenstreit, D. & Karmakar, P. Transcriptional bursting: from fundamentals to novel insights. Biochem. Soc. Transactions 52, 1695–1702, DOI: 10.1042/BST20231286 (2024).

64. Stasevich, T. J. & McNally, J. G. Assembly of the transcription machinery: Ordered and stable, random and dynamic, or both? Chromosoma 120, 533–545, DOI: 10.1007/S00412-011-0340-Y (2011).

65. Goel, S. et al. SARS-CoV-2 Switches ‘on’ MAPK and NFB Signaling via the Reduction of Nuclear DUSP1 and DUSP5 Expression. Front. Pharmacol. 12, 631879, DOI: 10.3389/FPHAR.2021.631879/BIBTEX (2021).

66. The RECOVERY Collaborative Group. Dexamethasone in Hospitalized Patients with Covid-19. New Engl. J. Medicine 384, 693–704, DOI: 10.1056/NEJMoa2021436 (2021).

67. Stoecklin, G. et al. MK2-induced tristetraprolin:14-3-3 complexes prevent stress granule association and ARE-mRNA decay. The EMBO J. 23, 1313–1324, DOI: 10.1038/sj.emboj.7600163 (2004).

68. Haimovich, G. & Gerst, J. E. Single-molecule Fluorescence in situ Hybridization (smFISH) for RNA Detection in Adherent Animal Cells. Bio-protocol 8, e3070, DOI: 10.21769/BioProtoc.3070 (2018).

69. Stringer, C., Wang, T., Michaelos, M. & Pachitariu, M. Cellpose: a generalist algorithm for cellular segmentation. Nat. Methods 2020 18:1 18, 100–106, DOI: 10.1038/s41592-020-01018-x (2020).

70. Gillespie, D. T. A general method for numerically simulating the stochastic time evolution of coupled chemical reactions. J. Comput. Phys. 22, 403–434, DOI: 10.1016/0021-9991(76)90041-3 (1976).

