## Supplemental Information for "Endogenous single-cell experiments and stochastic modeling reveal control mechanisms of glucocorticoid receptor dynamics and DUSP1 transcription"

### 1 Stochastic modeling

#### 1.1 Overview of the Dex-GR-*DUSP1* expression model fit-and-predict pipeline in the SSIT

1. GR model selection and fit
2. *DUSP1* nuclear/transcription-site model selection and fit (using GR model results)
   (a) Build nuclear model
   (b) Compute FSP solutions for nuclear model
   (c) Build TS-analysis model
   (d) Compute hybrid ODE-FSP solutions for TS model
   (e) Define free parameters of nuclear model and log-priors over them
   (f) Compute nuclear likelihood and TS likelihood
3. *DUSP1* cytoplasmic model selection and fit (using GR results and *DUSP1* nuclear results)
   (a) Build cytoplasmic model, using fitted GR and nuclear *DUSP1* parameters
   (b) Define free parameters of the cytoplasmic *DUSP1* model
   (c) Run SSA and fit trajectories using ABC
   (d) Use a combined objective function from nuclear + cytoplasmic *DUSP1* marginals, TS likelihood, and prior
   (e) Fit cytoplasmic model using Metropolis-Hastings sampling
4. Fit TPL data to *DUSP1* model(s) and select final *DUSP1* model
5. Test predictive power of final *DUSP1* model by holding out Dex/TPL conditions

**Notation.** Each model is defined by its species, its reactions, and, for every reaction $j$, a propensity $w_j$ and a stoichiometry vector $\boldsymbol{s}_j$ (the net change in species counts when reaction $j$ fires). Reactions are written reactants $\to$ products, with $\varnothing$ a zeroth-order source or a degradation sink; the stoichiometry column lists only the nonzero entries of $\boldsymbol{s}_j$ as "species $\pm n$."

#### 1.2 Time-varying input signals

Three deterministic signals enter the propensities. They are evaluated at the current time $t$ (in minutes, with $t = 0$ the moment of dexamethasone addition).

The dexamethasone signal driving transport of GR is defined:

$$I_{\text{Dex}}(t) = \begin{cases} 0 & \text{for } t \leq 0 \\ \text{Dex}_0 \cdot \exp(-g_{\text{Dex}} \cdot t) & \text{for } t > 0 \end{cases} \tag{1}$$

$$I_{\text{TTP}}(t) = \begin{cases} 1 & \text{for } t \leq 0, \\ 1 + TTP_1 \left( \frac{t^\eta}{t^\eta + t_{\text{TTP}}^\eta} \right) & \text{for } t > 0. \end{cases} \tag{2}$$

The post triptolide shut off of transcription is defined by:

$$I_{\text{TPL}}(t) = \begin{cases} 1 & \text{for } t \leq t_{\text{TPL}}, \\ \exp(-k_{\text{TPL}} \cdot (t - t_{\text{TPL}})) & \text{for } t > t_{\text{TPL}}. \end{cases} \tag{3}$$

To represent the time delay between transcription initiation and appearance of mature nuclear mRNA, the signals $I_{\text{Dex}}$ and $I_{\text{TPL}}$ are delayed by an amount $\tau_{\text{elong}}$ when calculating their effects on nuclear mRNA. In the absence of triptolide $t_{\text{TPL}} \to \infty$ so that $I_{\text{TPL}} \equiv 1$.

##### 1.3 Parameter glossary

Each parameter is listed with its symbol, description (units), and the corresponding reaction in the mechanistic-model schematic (Fig. S11).

**Glucocorticoid receptor (GR).**

- $\gamma_{\mathrm{GRcyt}}$ cytoplasmic GR degradation rate ($\mathrm{min}^{-1}$) — reaction 1
- $k_{\mathrm{g}}$ GR (cytoplasmic) synthesis rate ($\mathrm{min}^{-1}$) — reaction 2
- $k_{\mathrm{cn0}}$ basal cyto→nuc import rate ($\mathrm{min}^{-1}$) — reaction 3
- $k_{\mathrm{cn1}}$ Dex-dependent import rate ($\mathrm{min}^{-1}$) — reaction 3
- $M_{\mathrm{Dex}}$ half-saturation of Dex import (nM) — reaction 3
- $k_{\mathrm{nc}}$ GR nuclear export rate ($\mathrm{min}^{-1}$) — reaction 4
- $\gamma_{\mathrm{GRnuc}}$ nuclear GR degradation rate ($\mathrm{min}^{-1}$) — reaction 5
- $D_0$ Dex dose / input amplitude (nM) — input $I_{\mathrm{Dex}}(t)$
- $\gamma_{\mathrm{Dex}}$ Dex signal decay rate ($\mathrm{min}^{-1}$) — input $I_{\mathrm{Dex}}(t)$
- $k_{\mathrm{nc1}}$ Dex modulation of export ($\mathrm{min}^{-1}$) — GR Dex-export model only

***DUSP1* transcription.**

- $k_{\mathrm{on},0}$ basal gene activation rate ($\mathrm{min}^{-1}$) — reaction 6
- $k_{\mathrm{on},1}$ nucGR-dependent activation rate ($\mathrm{min}^{-1}$) — reaction 6
- $k_{\mathrm{off}}$ basal gene inactivation rate ($\mathrm{min}^{-1}$) — reaction 7
- $m_{\mathrm{koff}}$ nucGR modulation of inactivation ($\mathrm{molecules}^{-1}$) — reaction 7
- $k_{\mathrm{r,off}}$ leak / off-state transcription rate ($\mathrm{min}^{-1}$) — reaction 8
- $k_{\mathrm{r,on},0}$ basal on-state transcription rate ($\mathrm{min}^{-1}$) — reaction 8
- $k_{\mathrm{r,on},1}$ nucGR-dependent transcription rate ($\mathrm{min}^{-1}$) — reaction 8

**Cytoplasmic *DUSP1* mRNA.**

- $k_{\mathrm{Nuc2Cyt}}$ *DUSP1* mRNA nuclear export rate ($\mathrm{min}^{-1}$) — reaction 9
- $k_{\mathrm{degCyt},0}$ basal cytoplasmic degradation rate ($\mathrm{min}^{-1}$) — reaction 10a
- $k_{\mathrm{degCyt},1}$ TTP-bound (enhanced) degradation rate ($\mathrm{min}^{-1}$) — reaction 10c
- $k_{\mathrm{DTTP}}$ TTP–mRNA binding saturation (molecules) — semi-mechanistic $I_{\mathrm{TTP}}$ model
- $TTP_1$ max fold-change in TTP activity (unitless) — input $I_{\mathrm{TTP}}(t)$
- $t_{\mathrm{TTP}}$ half-max time for TTP activity (min) — input $I_{\mathrm{TTP}}(t)$
- $\eta$ Hill coefficient, TTP time-dependence (unitless) — input $I_{\mathrm{TTP}}(t)$

**Mechanistic TTP module (extended model).**

- $k_{\mathrm{on},1}^{\mathrm{TTP}}$ nucGR-dependent TTP-gene activation ($\mathrm{min}^{-1}$) — reaction 11
- $k_{\mathrm{off}}^{\mathrm{TTP}}$ TTP-gene inactivation ($\mathrm{min}^{-1}$) — reaction 12
- $k_{\mathrm{r,on}}^{\mathrm{TTP}}$ TTP transcription rate ($\mathrm{min}^{-1}$) — reaction 13
- $k_{\mathrm{Nuc2Cyt}}^{\mathrm{TTP}}$ TTP mRNA nuclear export ($\mathrm{min}^{-1}$) — reaction 14
- $k_{\mathrm{tl}}$ TTP translation rate ($\mathrm{min}^{-1}$) — reaction 15
- $\gamma_{\mathrm{p}}$ TTP protein degradation ($\mathrm{min}^{-1}$) — reaction 18
- $k_{\mathrm{b}}$ TTP protein–mRNA binding rate ($\mathrm{min}^{-1}$) — reactions 10b, 16
- $k_{\mathrm{u}}$ TTP protein–mRNA unbinding rate ($\mathrm{min}^{-1}$) — reactions $10\mathrm{b}^{-1}$, $16^{-1}$
- $\eta_{\mathrm{TTP}}$ Hill coefficient, TTP–mRNA binding (unitless) — reactions 10b, 16
- $\gamma_0^{\mathrm{TTP}}$ basal TTP mRNA degradation ($\mathrm{min}^{-1}$) — reaction 17

#### 1.4 Alternative models for Glucocorticoid Receptor translocation

**Species.** $GR_{\text{cyt}}$ (cytoplasmic GR) and $GR_{\text{nuc}}$ (nuclear GR). the initial condition for all species is the steady state distribution under the condition of no Dex.

Reactions are numbered 1–5 as in the graphical abstract (Fig. 1, right). Each reaction transfers, creates, or removes a single molecule, so its stoichiometry is given directly by the reaction arrow. The best-fit model ("Nuc + Cyt Deg", Fig. 3) contains all five reactions:

$$
\begin{aligned}
&(1)\quad GR_{\text{cyt}} \to \varnothing: \; w_1 = \gamma_{\text{GRcyt}} GR_{\text{cyt}} && \text{(cytoplasmic GR degradation)}\\
&(2)\quad \varnothing \to GR_{\text{cyt}}: \; w_2 = k_g && \text{(GR synthesis)}\\
&(3)\quad GR_{\text{cyt}} \to GR_{\text{nuc}}: \; w_3 = \left(k_{cn0} + k_{cn1}\frac{I_{\text{Dex}}(t)}{M_{\text{Dex}}+I_{\text{Dex}}(t)}\right) GR_{\text{cyt}} && \text{(Dex-induced import)}\\
&(4)\quad GR_{\text{nuc}} \to GR_{\text{cyt}}: \; w_4 = k_{nc}\, GR_{\text{nuc}} && \text{(nuclear export)}\\
&(5)\quad GR_{\text{nuc}} \to \varnothing: \; w_5 = \gamma_{\text{GRnuc}} GR_{\text{nuc}} && \text{(nuclear GR degradation)}
\end{aligned}
$$

Here, the Dex-dependent import term is active only after Dex addition as defined in Eq. 1.

The four candidate models (Fig. 3) differ only in their degradation/translocation structure:

- **Nuc + Cyt Deg** (best fit; Import, $\gamma_{\text{GRnuc}}$, $\gamma_{\text{GRcyt}}$; 8 parameters): all reactions 1–5.
- **Cyt Deg** (Import, $\gamma_{\text{GRcyt}}$; 7 parameters): set $\gamma_{\text{GRnuc}} = 0$, removing reaction 5.
- **Nuc Deg** (Import, $\gamma_{\text{GRnuc}}$; 7 parameters): set $\gamma_{\text{GRcyt}} = 0$, removing reaction 1.
- **Dex Export** (Export, $\gamma_{\text{GRnuc}}$, $\gamma_{\text{GRcyt}}$; 8 parameters): keep reactions 1, 2, 5 and replace reactions 3–4 below.

In the Dex Export model, Dex inhibits nuclear export rather than enhancing import. Reactions 3–4 become:

$$
\begin{aligned}
&(3)\quad GR_{\text{cyt}} \to GR_{\text{nuc}}: \; w_3 = k_{cn0}\, GR_{\text{cyt}}\\
&(4)\quad GR_{\text{nuc}} \to GR_{\text{cyt}}: \; w_4 = \max\left(0,\; k_{nc} - k_{nc1}\frac{I_{\text{Dex}}(t)}{M_{\text{Dex}}+I_{\text{Dex}}(t)}\right) GR_{\text{nuc}}
\end{aligned}
$$

#### 1.5 Alternative models for nuclear *DUSP1* transcriptional activation

**Species.** $Gene_{\text{Off}}$ and $Gene_{\text{On}}$ (the inactive and active *DUSP1* promoter states) and $DUSP1_{\text{nuc}}$ (nuclear *DUSP1* mRNA). Nuclear GR enters as the time-varying input $GR_{\text{nuc}}(t)$ taken from the fitted GR model.

All five candidate models (Fig. S6, M1–M5) share the same four reactions (numbered 6-9 as shown in graphical abstract Fig. 1, right) and differ only in which terms couple $GR_{\text{nuc}}$ to the promoter and to transcription. The full (superset) propensities are:

$$
\begin{aligned}
&(6)\quad Gene_{\text{Off}} \to Gene_{\text{On}}: && w_1 = \left(k_{on,0} + k_{on,1}\, GR_{\text{nuc}}\right) Gene_{\text{Off}}\\
&(7)\quad Gene_{\text{On}} \to Gene_{\text{Off}}: && w_2 = \frac{k_{off}}{1 + m_{koff}\, GR_{\text{nuc}}} Gene_{\text{On}}\\
&(8)\quad \varnothing \to DUSP1_{\text{nuc}}: && w_3 = \left(k_{r,off} + \left(k_{r,on,0} + k_{r,on,1}\, GR_{\text{nuc}}\right) Gene_{\text{On}}\right) I_{\text{TPL}}(t)\\
&(9)\quad DUSP1_{\text{nuc}} \to \varnothing: && w_4 = k_{Nuc2Cyt}\, DUSP1_{\text{nuc}}
\end{aligned}
$$

Reactions 6–9 are promoter activation, promoter inactivation, transcription, and nuclear export, respectively. Here $GR_{\text{nuc}}$ can act at three points: promoter activation ($k_{on,1}$), promoter inactivation ($m_{koff}$), and the on-state transcription rate ($k_{r,on,1}$). Each model retains a subset of these couplings, holding the others at zero:

- **M1 – Repression:** $GR_{\text{nuc}}$ slows promoter inactivation only ($m_{koff}$ free; $k_{on,1} = k_{r,on,1} = 0$).
- **M2 – Activation:** $GR_{\text{nuc}}$ accelerates promoter activation only ($k_{on,1}$ free; $m_{koff} = k_{r,on,1} = 0$).
- **M3 – Production:** $GR_{\text{nuc}}$ raises the transcription rate only ($k_{r,on,1}$ free; $k_{on,1} = m_{koff} = 0$).
- **M4 – General Activation:** $GR_{\text{nuc}}$ accelerates activation, slows inactivation, and raises transcription ($k_{on,1}$, $m_{koff}$, $k_{r,on,1}$ all free).
- **M5 – Dual Regulation:** $GR_{\text{nuc}}$ accelerates activation and slows inactivation ($k_{on,1}$, $m_{koff}$ free; $k_{r,on,1} = 0$), and the transcription term is reduced by dropping the basal leak ($k_{r,off} = 0$), so reaction 8 becomes $w_3 = k_{r,on,0}\, Gene_{\text{On}}\, I_{\text{TPL}}(t)$.

Models M1–M4 retain the low basal off-state transcription rate $k_{r,off}$; M5 is a reduction of M4 that removes both $k_{r,off}$ and $k_{r,on,1}$.

##### 1.6 Alternative models for cytoplasmic *DUSP1* mRNA degradation

**Species.** $DUSP1_{\text{cyt}}$ (cytoplasmic *DUSP1* mRNA). The cytoplasmic compartment modifies one reaction and adds one reaction (reactions 9 and 10, respectively in the graphical abstract, Fig. 1 (right)):

$$(9)\quad DUSP1_{\text{nuc}} \to DUSP1_{\text{cyt}} : \qquad w_1 = k_{Nuc2Cyt}\, DUSP1_{\text{nuc}}$$
$$(10)\quad DUSP1_{\text{cyt}} \to \varnothing : \qquad w_{\text{deg}}$$

Reaction 9 is nuclear export (the same $k_{Nuc2Cyt}$ process as in the nuclear model, now extended to track the cytoplasmic product). The four candidate models differ only in the degradation propensity $w_{\text{deg}}$ in reaction 10.

Free mRNA degrades at the basal rate $k_{degCyt,0}$ and TTP-bound mRNA at the enhanced rate $k_{degCyt,1}$, so

$$w_{\text{deg}} = k_{degCyt,0}\, r_{\text{cyt}} + (k_{degCyt,1} - k_{degCyt,0})\, r^*_{\text{cyt}}, \tag{4}$$

where (writing $r_{\text{cyt}} \equiv DUSP1_{\text{cyt}}$ for brevity) $r^*_{\text{cyt}}$ is the TTP-bound mRNA, obtained from a rapid-equilibrium binding assumption.

**Saturating TTP-bound term.** Total cytoplasmic mRNA $r_{\text{cyt}} = r_{\text{cyt,F}} + r^*_{\text{cyt}}$ splits into free and TTP-bound pools, and total TTP $TTP_{\text{tot}} = TTP_{\text{F}} + r^*_{\text{cyt}}$ into free and mRNA-bound pools. Assuming rapid reversible binding of free mRNA and free TTP,

$$r_{\text{cyt,F}} + TTP_{\text{F}} \underset{k_-}{\overset{k_+}{\rightleftharpoons}} r^*_{\text{cyt}}, \tag{5}$$

the equilibrium balance $k_+\, r_{\text{cyt,F}}\, TTP_{\text{F}} = k_-\, r^*_{\text{cyt}}$ defines the dissociation constant

$$k_{DTTP} \equiv \frac{k_-}{k_+} = \frac{r_{\text{cyt,F}}\, TTP_{\text{F}}}{r^*_{\text{cyt}}} = \frac{(r_{\text{cyt}} - r^*_{\text{cyt}})(TTP_{\text{tot}} - r^*_{\text{cyt}})}{r^*_{\text{cyt}}}. \tag{6}$$

Rearranging gives a quadratic in $r^*_{\text{cyt}}$,

$${r^*_{\text{cyt}}}^2 - (r_{\text{cyt}} + TTP_{\text{tot}} + k_{DTTP})\, r^*_{\text{cyt}} + r_{\text{cyt}}\, TTP_{\text{tot}} = 0, \tag{7}$$

whose physically meaningful (smaller) root is

$$r^*_{\text{cyt}}(r_{\text{cyt}}, TTP_{\text{tot}}) = \tfrac{1}{2}\left[(r_{\text{cyt}} + TTP_{\text{tot}} + k_{DTTP}) - \sqrt{(r_{\text{cyt}} + TTP_{\text{tot}} + k_{DTTP})^2 - 4\, r_{\text{cyt}}\, TTP_{\text{tot}}}\right]. \tag{8}$$

The total TTP is either held constant or follows the time-varying signal $I_{\text{TTP}}(t)$ (Section 1.2). The four models combine these choices with (4):

- **Model 1 – Basal first-order degradation:** no TTP binding ($r^*_{\text{cyt}} = 0$),

  $$w_{\text{deg}} = k_{degCyt,0}\, r_{\text{cyt}}.$$

- **Model 2 – Saturable, no time-varying induction:** (8) with $TTP_{\text{tot}}$ fixed at its basal value 1,

  $$w_{\text{deg}} = k_{degCyt,0}\, r_{\text{cyt}} + (k_{degCyt,1} - k_{degCyt,0})\, r^*_{\text{cyt}}(r_{\text{cyt}}, 1).$$

- **Model 3 – Time-varying input, no saturation:** bound mRNA approximated as $I_{\text{TTP}}(t)\, r_{\text{cyt}}$,

  $$w_{\text{deg}} = k_{degCyt,0}\, r_{\text{cyt}} + (k_{degCyt,1} - k_{degCyt,0})\, I_{\text{TTP}}(t)\, r_{\text{cyt}}.$$

- **Model 4 – Time-varying, saturable (best fit):** (8) with $TTP_{\text{tot}} = I_{\text{TTP}}(t)$,

  $$w_{\text{deg}} = k_{degCyt,0}\, r_{\text{cyt}} + (k_{degCyt,1} - k_{degCyt,0})\, r^*_{\text{cyt}}(r_{\text{cyt}}, I_{\text{TTP}}(t)).$$

##### 1.7 Extension to a fully mechanistic model

To extend the analysis to a fully mechanistic model, Reactions 1 to 9 were retained and reactions 10 - 17 were added as illustrated in Fig. S11. These additional reactions include:

$$(10_a) \quad DUSP1_{\text{cyt}} \rightarrow \varnothing : \qquad w_{10\text{a}} = k_{degCyt,0}\, DUSP1_{\text{cyt}}$$

$$(10_b) \quad DUSP1_{\text{cyt}} + TTP_{\text{Prot}} \rightarrow DUSP1.TTP_{\text{Prot}} : \qquad w_{10\text{b}} = k_b\, DUSP1_{\text{cyt}} \left(TTP_{\text{Prot}} + TTP_{\text{Prot}}^{\eta_{\text{TTP}}}\right)$$

$$(10_b^{-1}) \quad DUSP1.TTP_{\text{Prot}} \rightarrow DUSP1_{\text{cyt}} + TTP_{\text{Prot}} : \qquad w_{10\text{b}^{-1}} = k_u\, DUSP1.TTP_{\text{Prot}}$$

$$(10_c) \quad DUSP1.TTP_{\text{Prot}} \rightarrow TP_{\text{Prot}} : \qquad w_{10\text{c}} = k_{degCyt,1}\, DUSP1.TTP_{\text{Prot}}$$

$$(11) \quad gTTP_{\text{Off}} \rightarrow gTTP_{\text{On}} : \qquad w_{11} = \left(k_{on,0} + k_{on,1}^{\text{TTP}}\, GR_{\text{nuc}}\right) gTTP_{\text{Off}}$$

$$(12) \quad gTTP_{\text{On}} \rightarrow gTTP_{\text{Off}} : \qquad w_{12} = k_{\text{off}}^{\text{TTP}}\, gTTP_{\text{On}}$$

$$(13) \quad gTTP_{\text{On}} \rightarrow gTTP_{\text{On}} + TTP_{\text{nuc}} : \qquad w_{13} = k_{r,on}^{\text{TTP}}\, gTTP_{\text{On}}\, I_{\text{TPL}}(t)$$

$$(14) \quad TTP_{\text{nuc}} \rightarrow TTP_{\text{cyt}} : \qquad w_{14} = k_{Nuc2Cyt}\, TTP_{\text{nuc}}$$

$$(15) \quad TTP_{\text{cyt}} \rightarrow TTP_{\text{cyt}} + TTP_{Prot} : \qquad w_{15} = k_{\text{tl}}\, TTP_{\text{cyt}}$$

$$(16_a) \quad TTP_{\text{cyt}} \rightarrow \varnothing : \qquad w_{16\text{a}} = k_0^{\text{TTP}}\, TTP_{\text{cyt}}$$

$$(16_b) \quad TTP_{\text{cyt}} + TTP_{\text{Prot}} \rightarrow TTP.TTP : \qquad w_{16} = k_b\, TTP_{\text{cyt}}\, TTP_{\text{Prot}}$$

$$(16_b^{-1}) \quad TTP.TTP_{\text{Prot}} \rightarrow TTP\text{cyt} + TTP_{\text{Prot}} : \qquad w_{16^{-1}} = k_u\, TTP.TTP_{\text{Prot}}$$

$$(16_c) \quad TTP.TTP_{\text{Prot}} \rightarrow TP_{\text{Prot}} : \qquad w_{17} = k_{degCyt,1}\, TTP.TTP_{\text{Prot}}$$

$$(17) \quad TTP_{\text{Prot}} \rightarrow \varnothing : \qquad w_{18} = \gamma_0^{\text{TTP}}\, TTP_{\text{Prot}}.$$

Here, reactions 10a and 16a represent linear degradation for *DUSP1* and *TTP* mRNA, reversible reactions 10b/10b$^{-1}$ and 16b/16b$^{-1}$ correspond to binding and unbinding to the TTP protein, and reactions 10c and 16c correspond to degradation of the mRNA in the complex with the TTP protein; Reactions 11-14 correspond to the activation, deactivation, transcription and transport of the TTP mRNA; and Reactions 15 and 17 correspond to the translation and degradation of the TTP protein.

#### Supplementary Figures

**a** **Number of probes:** 21
**Probe length:** 20±5 nt (plus FLAP-Y)
**GC content:** 45–55%
**Melting T** (nearest neighbor): 66–70 °C
**Reference genome:** ENST00000239223.4
**Design:** Oligostan (Mueller et al., Nat. Methods 2013)

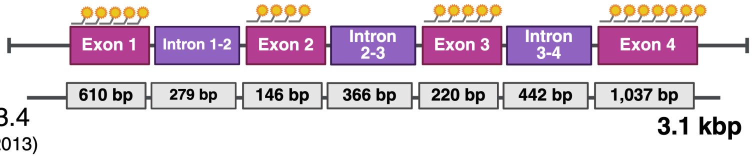

**b**

| Probe Label | Locus | Probe Sequence |
|---|---|---|
| DUSP1_P1_Exon1 | 101 - 127 | AGCCGGCGATGTGGCCGGCGTTGAAATTACACTCGGACCTCGTCGACATGCATT |
| DUSP1_P2_Exon1 | 132 - 158 | CGCACGATGGTGCTGAAGCGCACGTTTTACACTCGGACCTCGTCGACATGCATT |
| DUSP1_P3_Exon1 | 189 - 215 | CGGAGCTCGGCGTTGGGCACGATGTGTTACACTCGGACCTCGTCGACATGCATT |
| DUSP1_P4_Exon1 | 237 - 263 | TCGTCCAGCAACACCACGGCGTGGTATTACACTCGGACCTCGTCGACATGCATT |
| DUSP1_P5_Exon1 | 347 - 378 | TTCGTATCCTCCTTTGAGGAAGAAGACTTGCTTACACTCGGACCTCGTCGACATGCATT |
| DUSP1_P6_Exon2 | 380 - 407 | CACAGCTCCGGGCAGGAAGCCGAAAACTTACACTCGGACCTCGTCGACATGCATT |
| DUSP1_P7_Exon2 | 411 - 437 | CTGAGCCCCATGGGGGTCGACTGTTTTTACACTCGGACCTCGTCGACATGCATT |
| DUSP1_P8_Exon2 | 452 - 478 | CAGATTCCGCGCTGTCAGGGACGCTATTACACTCGGACCTCGTCGACATGCATT |
| DUSP1_P9_Exon2 | 490 - 517 | CTGATCGTAGAGTGGGGTACTGCTTACACTCGGACCTCGTCGACATGCATT |
| DUSP1_P10_Exon3 | 550 - 576 | CTTGCGGGAAGCGTGATACGCACTGCTTACACTCGGACCTCGTCGACATGCATT |
| DUSP1_P11_Exon3 | 609 - 641 | AAATGGTTGGGACAATTGGCTGAGACGTTGATTTACACTCGGACCTCGTCGACATGCATT |
| DUSP1_P12_Exon3 | 645 - 674 | ACAGGGATGCTCTTGTACTGGTAGTGACCTTACACTCGGACCTCGTCGACATGCATT |
| DUSP1_P13_Exon3 | 676 - 702 | GCTGATGTCTGCCTTGTGGTTGTCCTTTACACTCGGACCTCGTCGACATGCATT |
| DUSP1_P14_Exon3 | 704 - 734 | TCTATGAAGTCAATGGCCTCGTTGAACCAGTTACACTCGGACCTCGTCGACATGCATT |
| DUSP1_P15_Exon4 | 736 - 762 | CACCCTTCCTCCAGCATTCTTGATGGTTACACTCGGACCTCGTCGACATGCATT |
| DUSP1_P16_Exon4 | 764 - 790 | GGGAAATGCCTGCCTGGCAGTGGACATTACACTCGGACCTCGTCGACATGCATT |
| DUSP1_P17_Exon4 | 799 - 831 | ATTAGTCCTCATAAGGTAAGCAAGGCAGATGGTTACACTCGGACCTCGTCGACATGCATT |
| DUSP1_P18_Exon4 | 833 - 860 | AACTCAAAGGCCTCGTCCAGCTTGACTTTACACTCGGACCTCGTCGACATGCATT |
| DUSP1_P19_Exon4 | 862 - 888 | AGAGATGATGCTTCGCCTCTGCTTCATTACACTCGGACCTCGTCGACATGCATT |
| DUSP1_P20_Exon4 | 1046 - 1074 | CTGAAGGTAGCTCAGCGCACTGTTCGTGTTACACTCGGACCTCGTCGACATGCATT |
| DUSP1_P21_Exon4 | 1076 - 1104 | TCAGCAGCTGGGAGAGGTCGTAATGGGGTTACACTCGGACCTCGTCGACATGCATT |
| FLAP Y-Cy5 | | Cy5/AA TGC ATG TCG ACG AGG TCC GAG TGT AA/Cy5 |

**c**

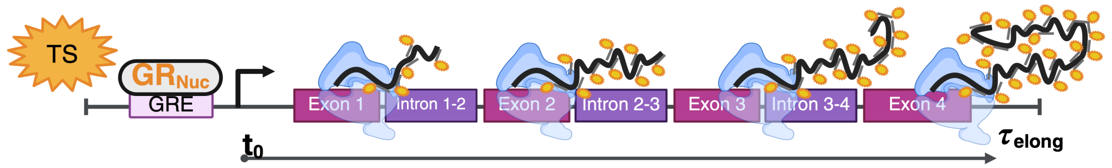

**d**

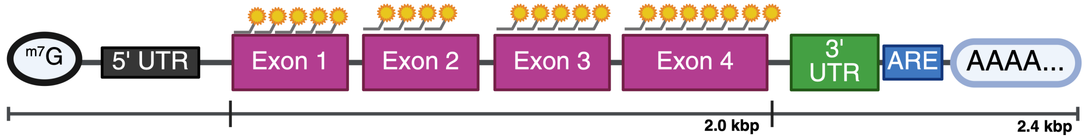

**Figure S1. Design and targeting scheme for the *DUSP1* smiFISH probe set.**
**(a)** Genomic organization of the *DUSP1* locus showing exons and introns with relative length scaling and the distribution of gene-specific smiFISH probes across exonic regions.
**(b)** Table listing the smiFISH probe labels, genomic loci, and sequences. Each probe consists of a gene-specific sequence extended with a FLAP-Y overhang for secondary probe binding.
**(c)** Cartoon of smiFISH detection at an active transcription site. As RNA polymerase progresses along the gene, nascent *DUSP1* transcripts are labeled by exon-targeting probes, producing a bright nuclear focus. The transcriptional elongation time, $\tau_{\text{elong}}$, defines the duration over which polymerases contribute to the TS signal and is used in TS-based model fits and transcriptional inhibition (triptolide) predictions.
**(d)** Cartoon of probe binding along a mature *DUSP1* mRNA, shown with a relative length scale. Probes tile the coding exons, resulting in a single fluorescent focus. The transcript includes a 5′ cap, coding exons, the 3′ UTR containing an ARE, and a poly(A) tail.

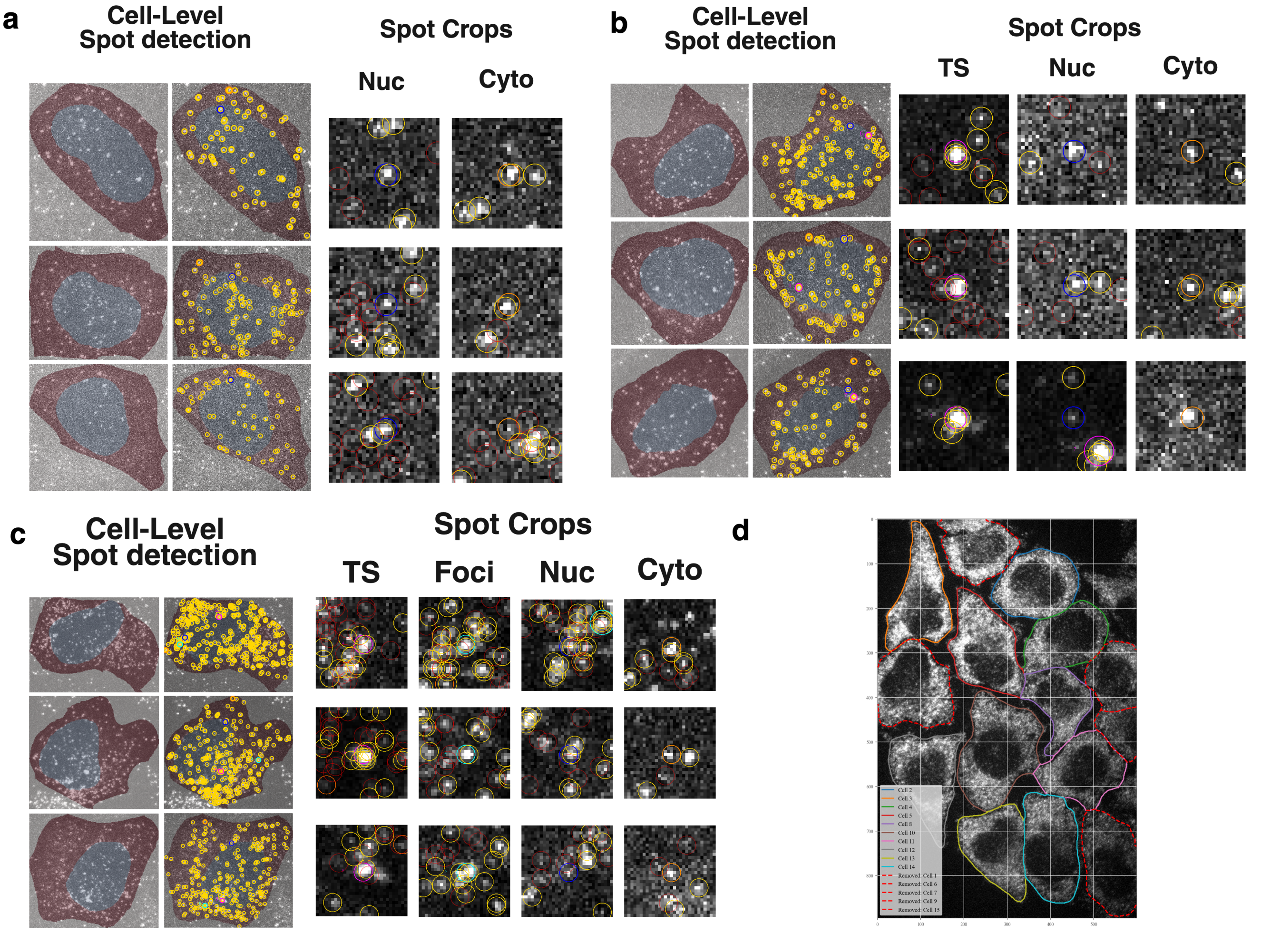

**Figure S2. Segmentation and smFISH spot detection for single-cell *DUSP1* quantification.**
**(a)** Representative *DUSP1* smiFISH images across selected times and Dex concentrations, shown with DAPI and 10 µm scale bars. These images illustrate the range of transcriptional responses used for single-cell quantification.
**(b)** Cell-level spot detection in cells without transcription sites or foci. For each example, both nuclear and cytoplasmic CellPose masks are displayed over the smiFISH image, with detected *DUSP1* mRNA spots shown in gold. The accompanying spot crops show nuclear and cytoplasmic detections, with red circles marking low-SNR candidates removed during filtering.
**(c)** Cell-level spot detection in cells containing transcription sites. Nuclear and cytoplasmic masks are displayed together with the detected *DUSP1* spots, including transcription-site detections. Spot crops illustrate transcription sites, nuclear mRNA spots, and cytoplasmic mRNA spots used for downstream quantification.
**(d)** Cell-level spot detection in cells containing both transcription sites and cytoplasmic foci. Nuclear and cytoplasmic masks are shown alongside the full set of detected spots. Spot crops include transcription sites, cytoplasmic foci, nuclear mRNA spots, and cytoplasmic mRNA spots, with removed low-SNR candidates indicated in red.
**(e)** Representative field of view with GAPDH-based cytoplasmic segmentation and removal of border-touching or partial cells. Each color denotes a unique cell used for downstream analysis.

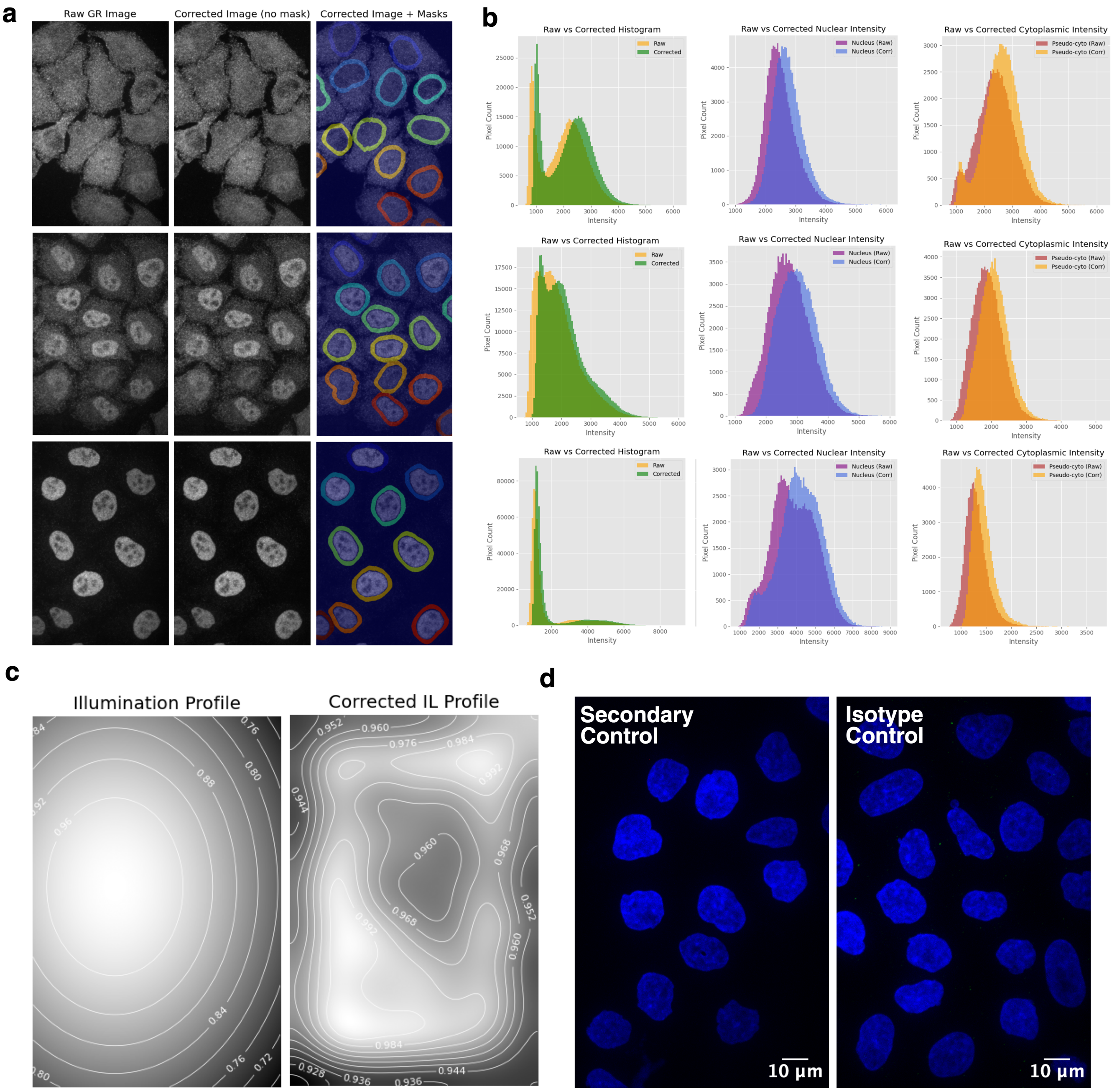

**Figure S3.** **Data-driven illumination correction and controls for GR images.**
**(a)** From left to right, raw GR images, illumination-corrected GR images, and corrected images with pseudo-cytoplasmic masks overlaid for three representative fields of view spanning low to high Dex stimulation from top to bottom.
**(b)** Pixel-intensity histograms corresponding to the fields of view in (a). From left to right, distributions of whole-image intensities (raw, yellow; corrected, green), nuclear GR intensities (raw, purple; corrected, blue), and pseudo-cytoplasmic GR intensities (raw, red; corrected, orange), showing improved uniformity after correction.
**(c)** Batch-derived illumination profile for the GR channel (left) and the corresponding profile after correction (right), both normalized to the range 0–1 and displayed as contour maps, demonstrating removal of the illumination gradient.
**(d)** Representative secondary-only and isotype control images of HeLa cells stained with DAPI, confirming minimal background signal in the GR channel. Scale bars, 10 µm.

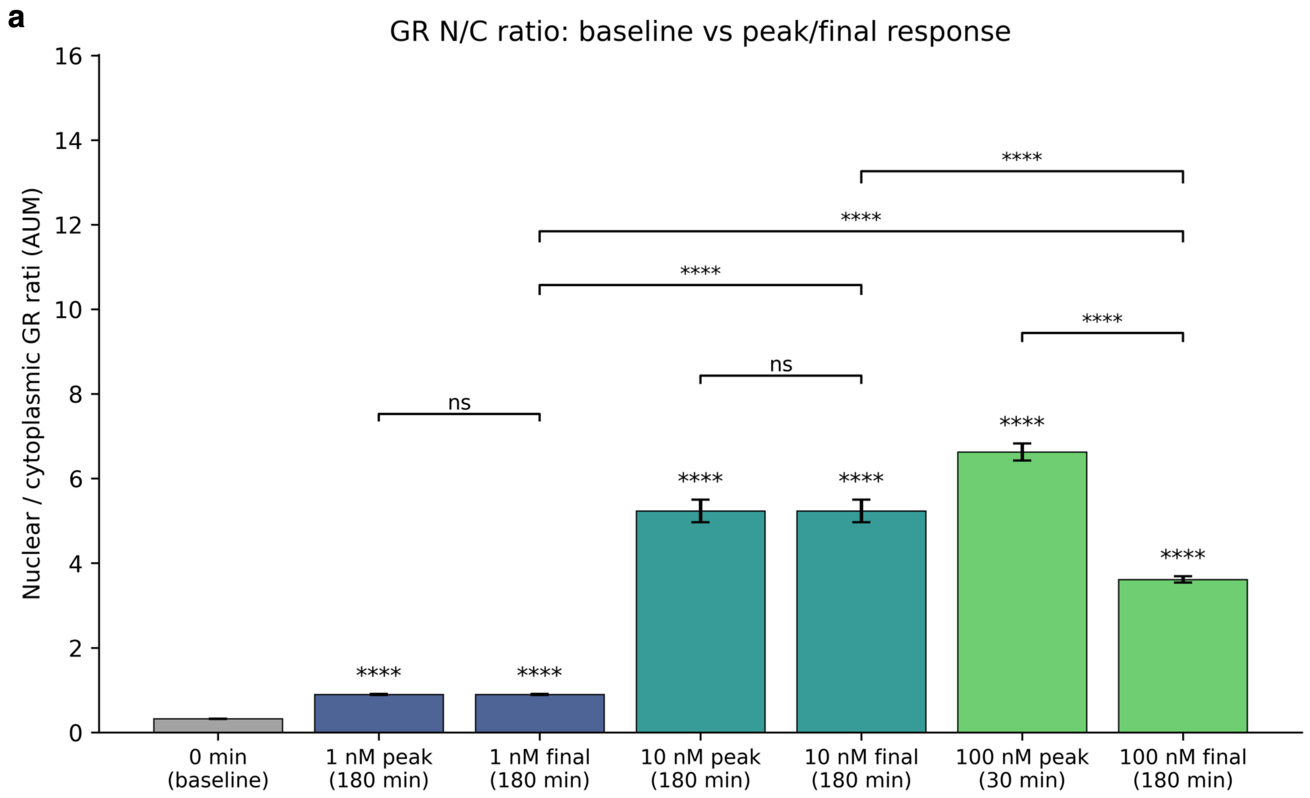

**Figure S4.** **Single-cell nuclear-to-cytoplasmic GR ratio across Dex concentrations and stimulation times.** Per-cell nuclear-to-cytoplasmic (N/C) GR mass ratio at baseline versus the peak and final response for each Dex concentration. Bars show the mean N/C ratio across single cells; error bars denote the standard error of the mean (SEM). "Peak" is the stimulation time at which the mean N/C ratio is maximal for that dose and "final" is the last measured time point (180 min); for 10 nM these coincide at 180 min. Stars above each bar denote Welch's two-sample $t$-tests against the unstimulated baseline; brackets denote peak-vs-final comparisons within each dose and endpoint (final-vs-final) comparisons across doses (****, $p < 10^{-4}$; ns, not significant).

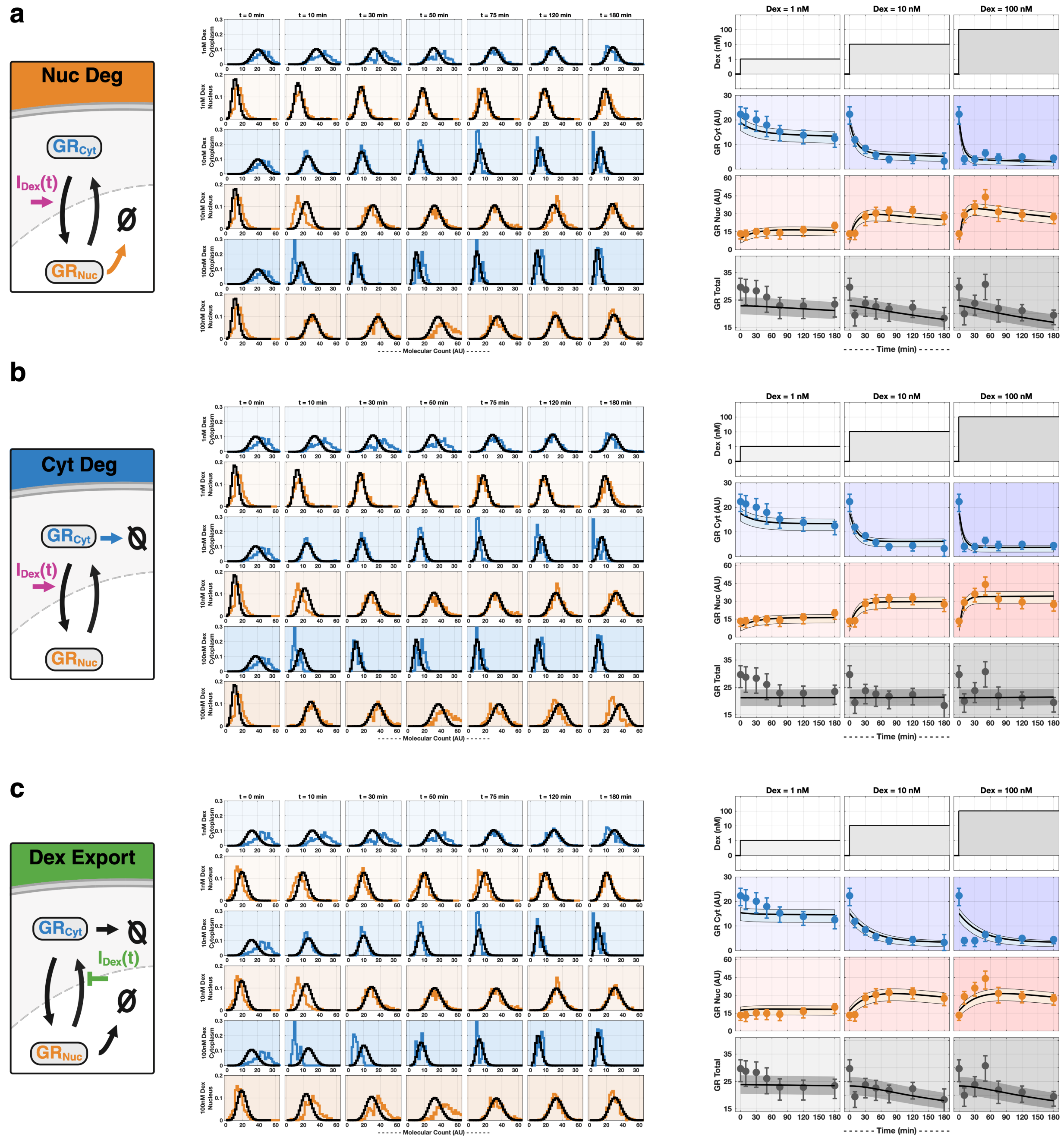

**Figure S5. Comparison of alternative GR transport and degradation models.**
**(a)** Nuclear degradation (Nuc Deg) model, in which GR is removed from the nuclear compartment. From left to right, the reaction schematic with import driven by $I_{\text{Dex}}(t)$; single-cell distributions of GR molecular count for cytoplasm and nucleus at 1, 10, and 100 nM Dex across time points ($t = 0$, 10, 30, 50, 75, 120, and 180 min), with measured distributions colored and shaded by compartment (cytoplasm, blue; nucleus, orange) and model predictions overlaid as black lines; and temporal trajectories at Dex = 1, 10, and 100 nM showing the imposed $I_{\text{Dex}}(t)$ input, $GR_{\text{Cyt}}$, $GR_{\text{Nuc}}$, and total GR.
**(b)** Cytoplasmic degradation (Cyt Deg) model, in which $GR_{\text{Cyt}}$ is degraded, shown with the same panel layout as (a).
**(c)** Dex-modulated export (Dex Export) model, in which $I_{\text{Dex}}(t)$ inhibits GR nuclear export with degradation from both compartments, shown with the same panel layout as (a).
In all distribution panels, data are colored and shaded by Dex concentration and model fits are black lines. In all trajectory panels, data points and bars denote the median ± IQR, and black lines with shading denote the model fit and its uncertainty.

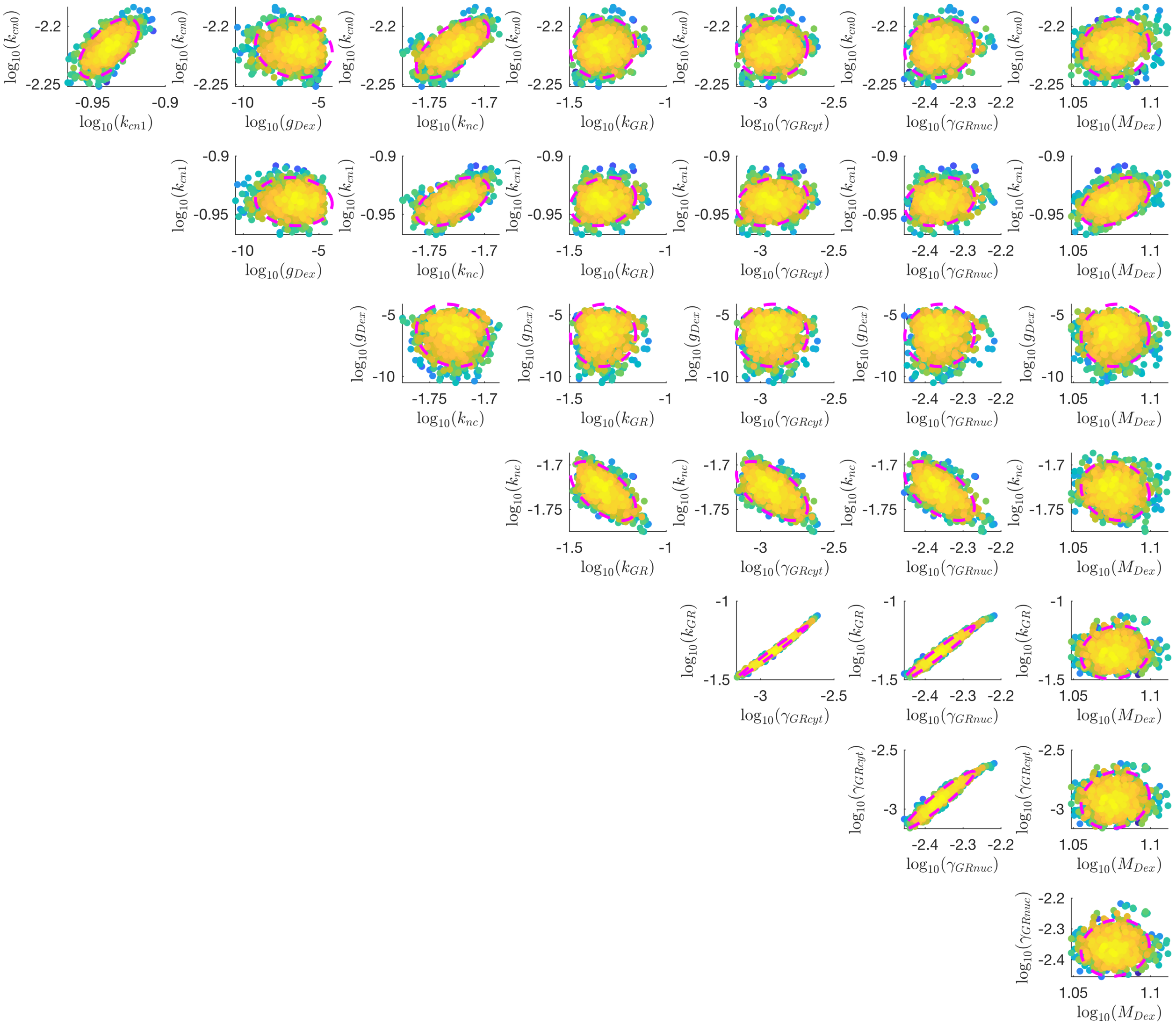

**Figure S6. Joint posterior distribution for the compartment-specific GR model.**
Uncertainty quantification for the chosen GR model, obtained by Metropolis–Hastings MCMC sampling. Each subplot shows the pairwise joint distribution of MCMC samples for two model parameters (in $\log_{10}$ space), with points colored by sample density. The magenta dashed ellipse marks the 95% confidence interval.

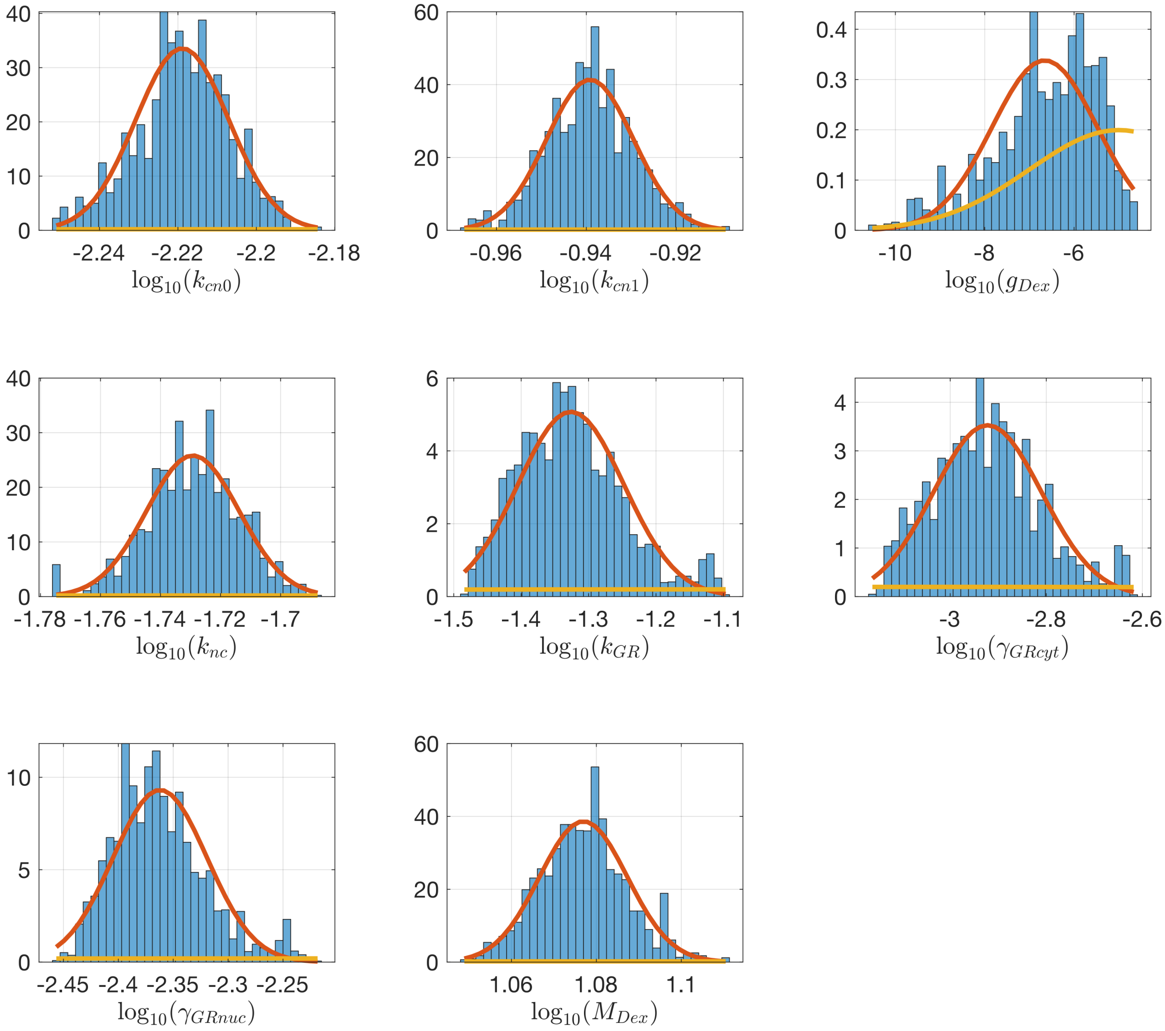

**Figure S7.** **Marginal posterior distributions for the compartment-specific GR model.**
Single-parameter marginal posteriors for the chosen GR model, obtained by Metropolis–Hastings MCMC sampling. Each panel shows one model parameter (in $\log_{10}$ space): the blue histogram is the distribution of MCMC samples, the red curve is a kernel density estimate of the marginal posterior, and the gold curve is the prior (lognormal with standard deviation of two orders of magnitude).

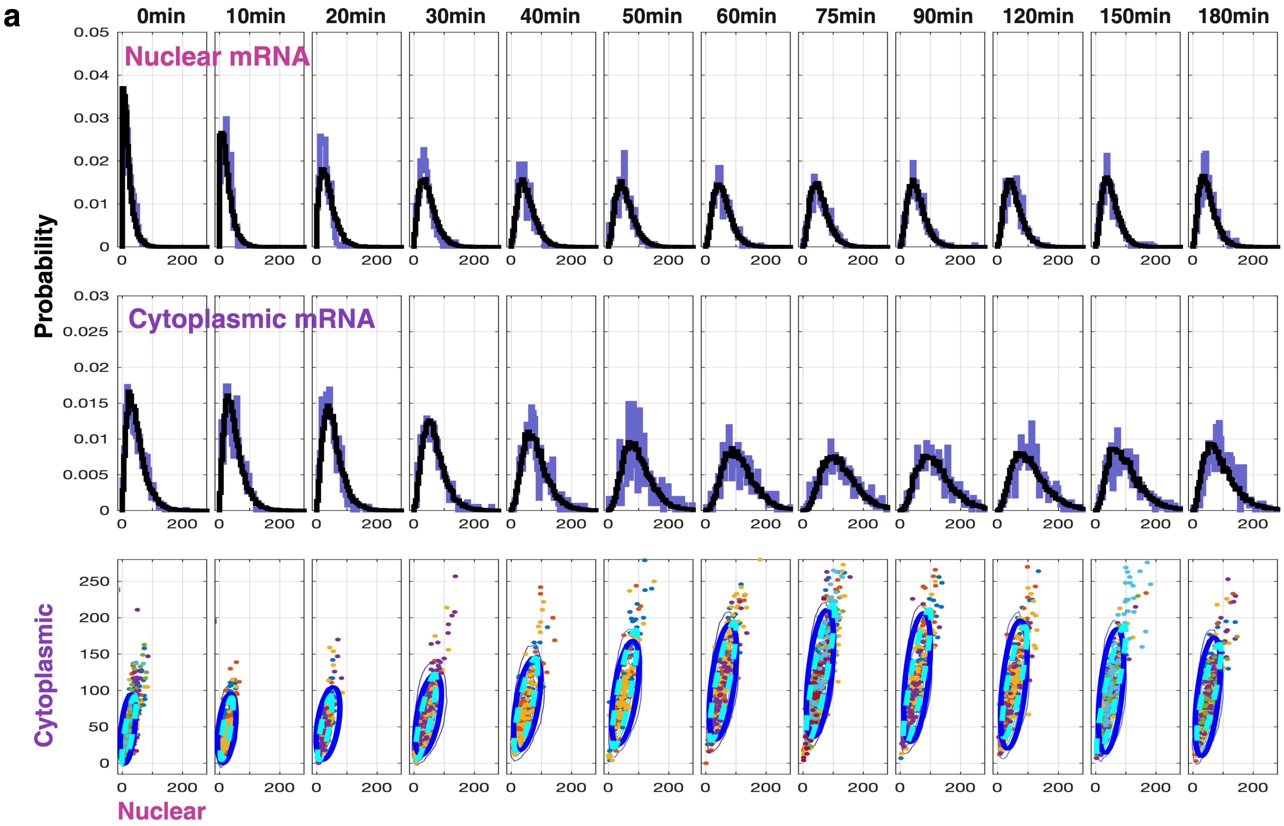

**Figure S8.** **Experimental *DUSP1* mRNA distributions used for model fitting at 100 nM Dex across all time points.**
**(a)** Distributions of nuclear and cytoplasmic *DUSP1* mRNA at all sampled time points following 100 nM Dex stimulation. Colored histograms show experimental measurements, with black curves indicating the corresponding model fits. Bottom row shows joint nuclear–cytoplasmic mRNA distributions, with data shown as points and model-fitted 95% confidence ellipses.

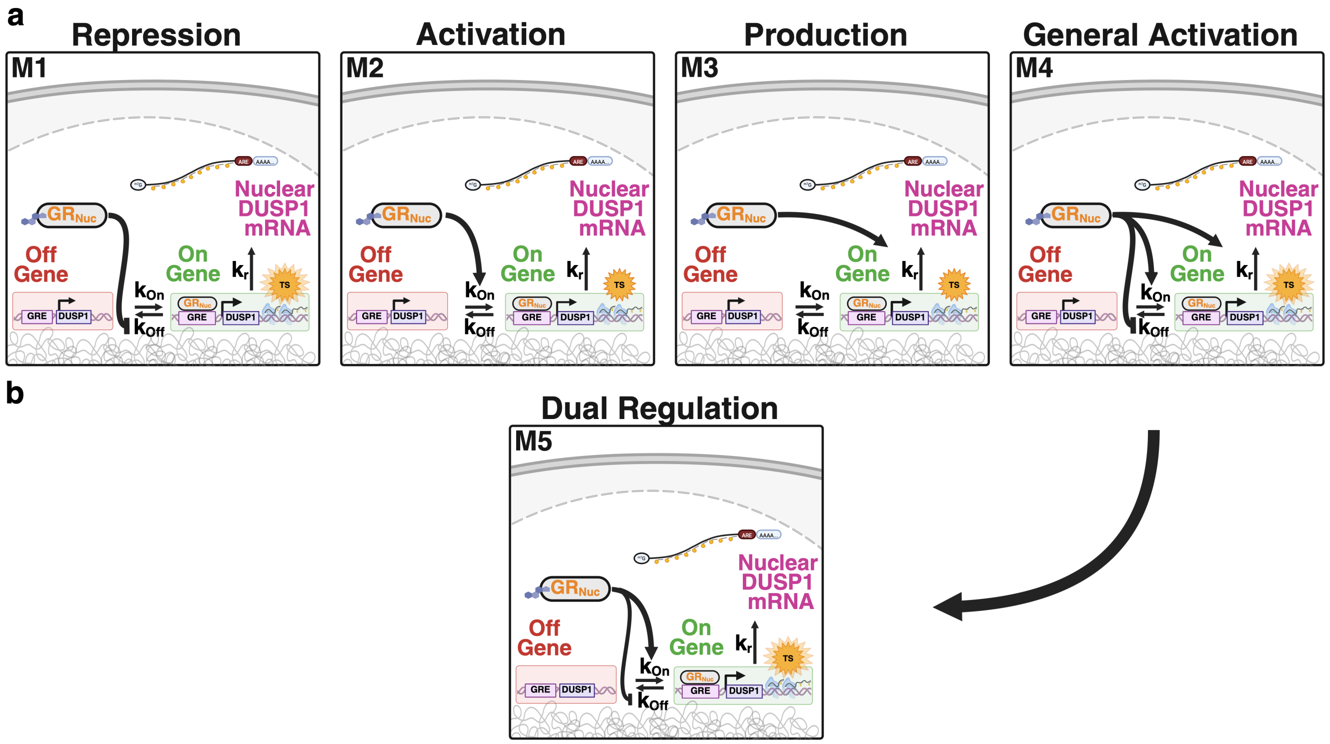

**Figure S9. Alternative models of GR-dependent *DUSP1* transcriptional regulation.**
Schematic representations of promoter-level models describing how nuclear GR ($GR_{Nuc}$) regulates *DUSP1* transcriptional bursting. The promoter transitions between OFF and ON states with rates $k_{on}$ and $k_{off}$, and produces nascent transcripts at rate $k_r$ at the transcription site (TS) when active. GR-dependent regulation is implemented through modulation of individual or combined parameters. **(a)** Candidate models M1–M4, each of which permits low-level transcription in the OFF state. In the repression model (M1), GR inhibits $k_{off}$; in the activation model (M2), GR stimulates $k_{on}$; in the production model (M3), GR increases $k_r$; and in the general activation model (M4), GR simultaneously modulates $k_{on}$, $k_{off}$, and $k_r$. **(b)** The reduced dual-regulation model (M5), in which GR stimulates $k_{on}$ and inhibits $k_{off}$ with no transcription in the OFF state. The thicker $k_{on}$ arrow relative to $k_{off}$ denotes the dominant contribution of burst-frequency activation. This reduced model provided the best agreement with the single-cell *DUSP1* data and was selected for subsequent analyses.

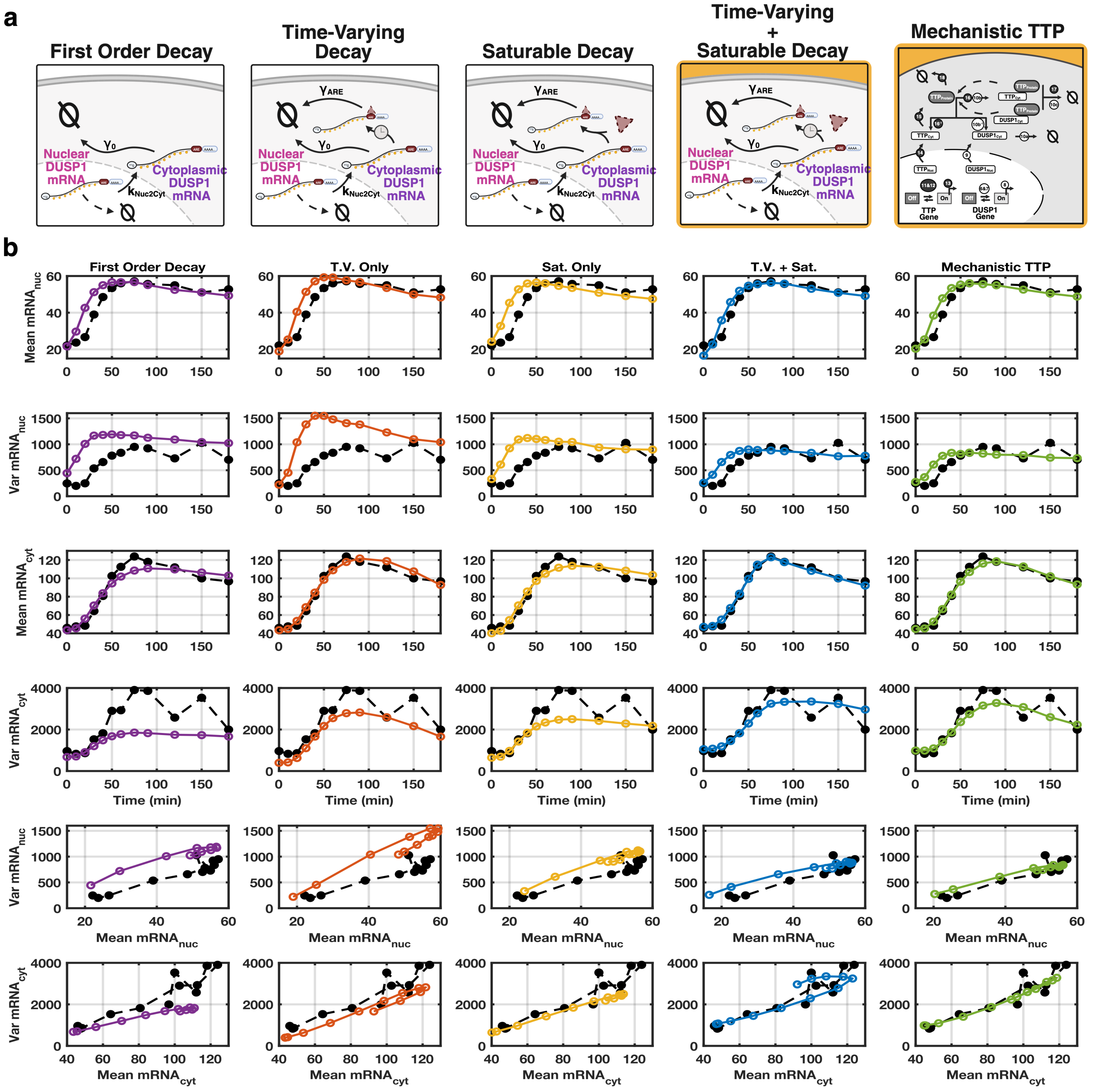

**Figure S10.** **Alternative models of cytoplasmic *DUSP1* mRNA degradation.** **(a)** Five candidate mechanisms for cytoplasmic *DUSP1* mRNA decay (left to right): **First Order Decay** (purple), a constant rate $\gamma_0$; **Time-Varying Decay** (orange) and **Saturable Decay** (yellow), each adding an ARE-mediated term $\gamma_{\mathrm{ARE}}$ with either explicit time dependence or abundance-dependent saturation; **Time-Varying + Saturable Decay** (blue), combining both into the phenomenological input $I_{\mathrm{TTP}}(t)$; and the full **Mechanistic TTP** model (green), which resolves TTP gene activation, transcription, translation, and TTP-catalyzed *DUSP1* mRNA degradation explicitly. Gold highlighting indicates the models retained for downstream analyses. **(b)** Model fits (colored open circles) versus single-cell smiFISH data (black dashed lines) for the 100 nM Dex time course. Rows, top to bottom: mean and variance of nuclear *DUSP1* mRNA versus time; mean and variance of cytoplasmic *DUSP1* mRNA versus time; and the variance–mean relationships for nuclear and cytoplasmic mRNA. All five models reproduce the mean trajectories comparably, so means alone do not discriminate among mechanisms. Only the models combining time-varying and saturable ARE-mediated decay (T.V. + Sat. and Mechanistic TTP) reproduce the variance dynamics and variance–mean relationships in both compartments; the First Order, T.V.-only, and Sat.-only models misestimate the nuclear and cytoplasmic variances. The agreement between the semi-mechanistic T.V. + Sat. model and the full Mechanistic TTP model supports $I_{\mathrm{TTP}}(t)$ as a compact, identifiable representation of TTP-driven ARE-dependent mRNA decay.

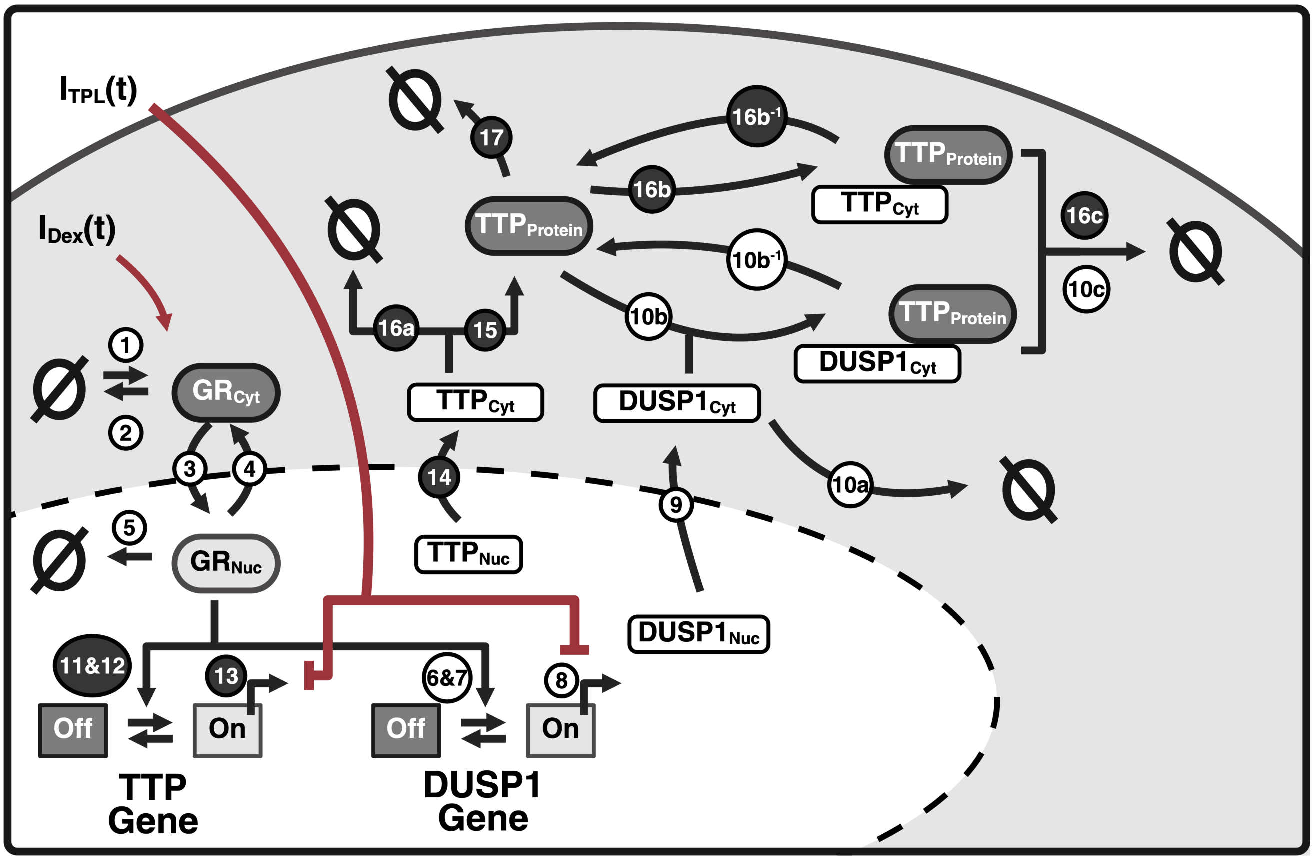

**Figure S11.** **Mechanistic model extension incorporating TTP-mediated post-transcriptional regulation.**
The semi-mechanistic model was extended to include explicit mechanisms for GR-driven TTP expression and control of mRNA degradation. New species include ON and OFF TTP gene states, nuclear and cytoplasmic *TTP* mRNA, TTP protein, and transient TTP–DUSP1 and TTP–TTP protein-mRNA complexes. Nuclear GR promotes *TTP* gene activation (**reaction 11**), followed by transcription (**reaction 13**), nuclear export (**reaction 14**), translation (**reaction 15**), and protein degradation (**reaction 17**), with basal gene deactivation (**reaction 12**). In the cytoplasm, TTP protein reversibly binds both *DUSP1* and *TTP* mRNA to form transient complexes (**reactions 10b/10b**$^{-1}$ and **16b/16b**$^{-1}$) and catalyzes their degradation (**reactions 10c** and **16c**); in both reactions the TTP protein acts catalytically and is not consumed. First-order decay of cytoplasmic *DUSP1* and *TTP* mRNA is also included (**reactions 10a** and **16a**). Reaction numbers shown in white circles denote processes retained from the semi-mechanistic model, whereas those in black circles denote the new reactions introduced in the full mechanistic extension. TPL input, $I_{\mathrm{TPL}}(t)$, inhibits transcription of both *DUSP1* and *TTP*. As *TTP* mRNA and protein were not directly measured, their dynamics are inferred through their effects on spatially resolved GR and *DUSP1* mRNA distributions.

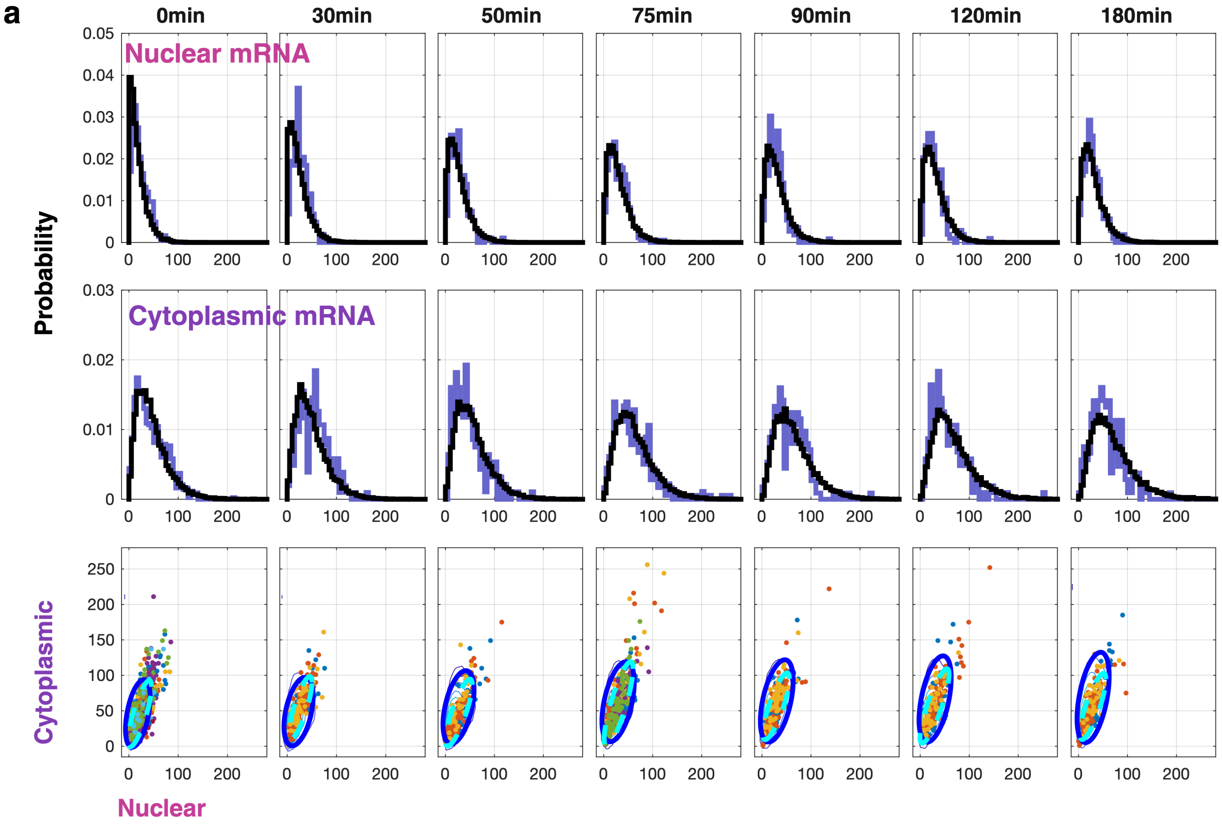

**Figure S12.** **Model fits and predictions for *DUSP1* mRNA dynamics at 1 nM Dex.**
**(a)** Distributions of nuclear and cytoplasmic *DUSP1* mRNA at the indicated time points following 1 nM Dex stimulation. Colored histograms show experimental measurements, with black curves indicating the corresponding model fits. Bottom row shows joint nuclear–cytoplasmic mRNA distributions, with data shown as points and model-predicted 95% confidence ellipses.

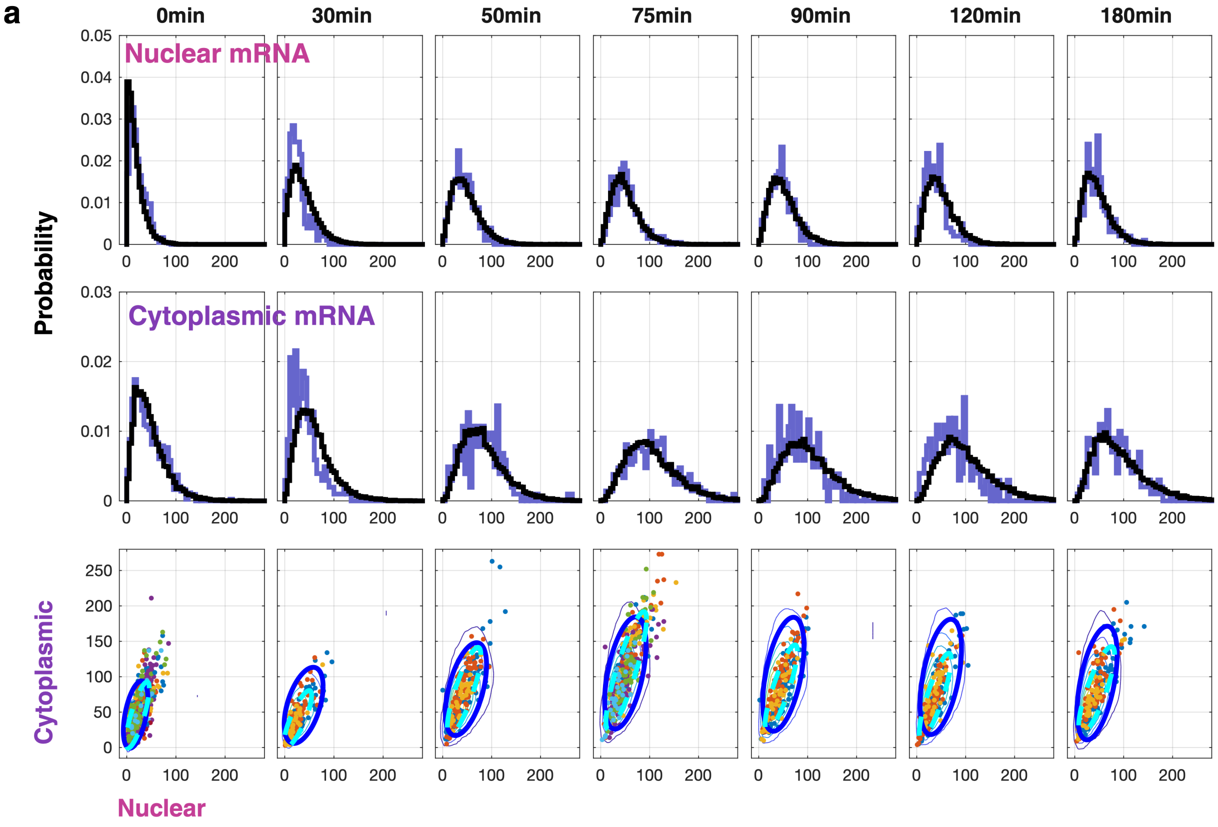

**Figure S13.** **Model fits and predictions for *DUSP1* mRNA dynamics at 10 nM Dex.**
**(a)** Distributions of nuclear and cytoplasmic *DUSP1* mRNA at the indicated time points following 10 nM Dex stimulation. Colored histograms show experimental measurements, with black curves indicating the corresponding model fits. Bottom row shows joint nuclear–cytoplasmic mRNA distributions, with data shown as points and model-predicted 95% confidence ellipses.

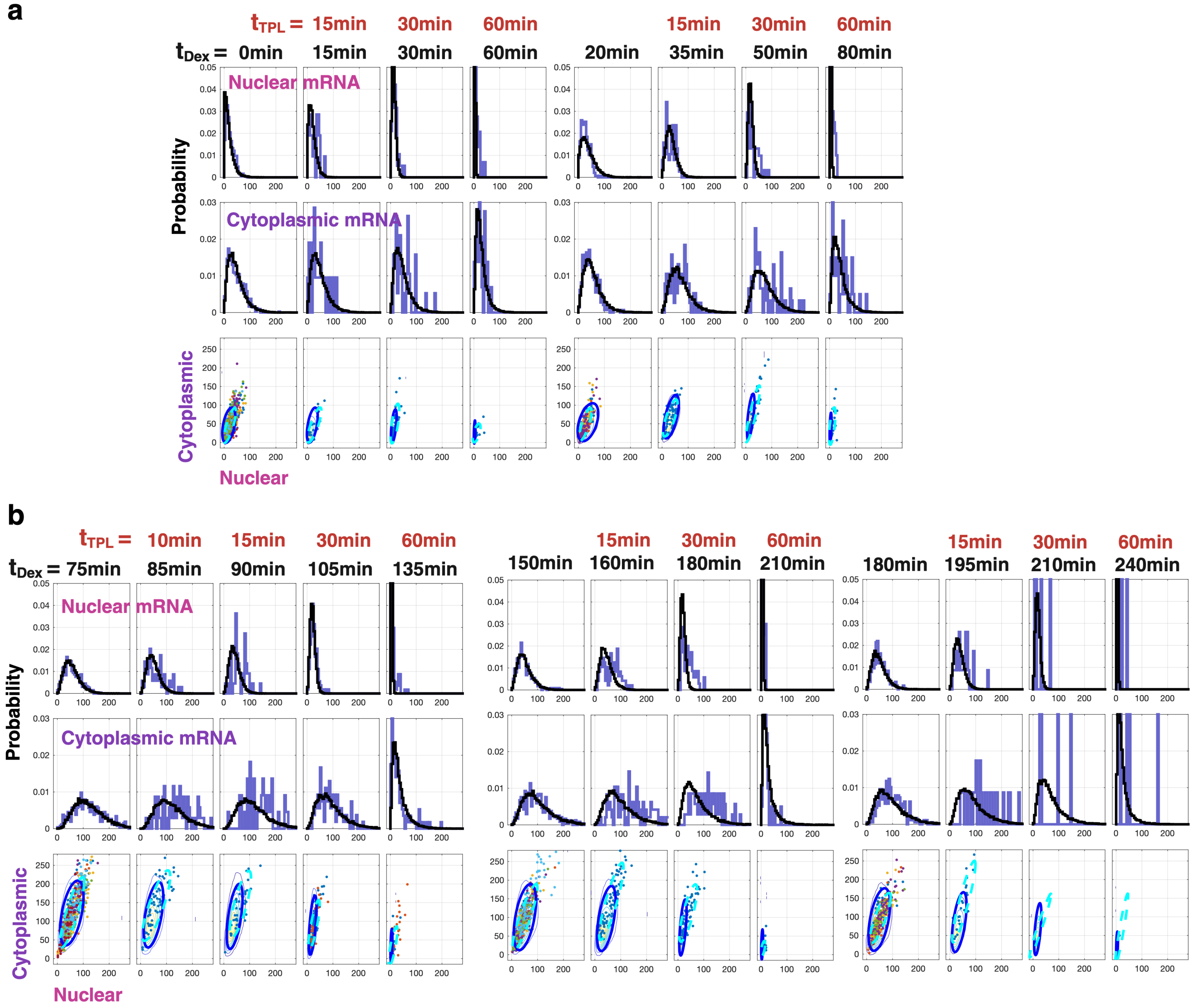

**Figure S14.** ***DUSP1*** **mRNA distributions under transcriptional inhibition by TPL.**

**(a)** Distributions of nuclear and cytoplasmic *DUSP1* mRNA and joint nuclear–cytoplasmic mRNA distributions across all sampled time points following transcriptional inhibition. Green-shaded panels correspond to control time points under 100 nM Dex stimulation without TPL (0, 20, 75, 150, and 180 min), while yellow-shaded panels correspond to time points following addition of 5 μM TPL. Colored histograms show experimental measurements, with black curves indicating the corresponding model predictions. Bottom rows show joint nuclear–cytoplasmic mRNA distributions, with data shown as points and model-predicted 95% confidence ellipses.

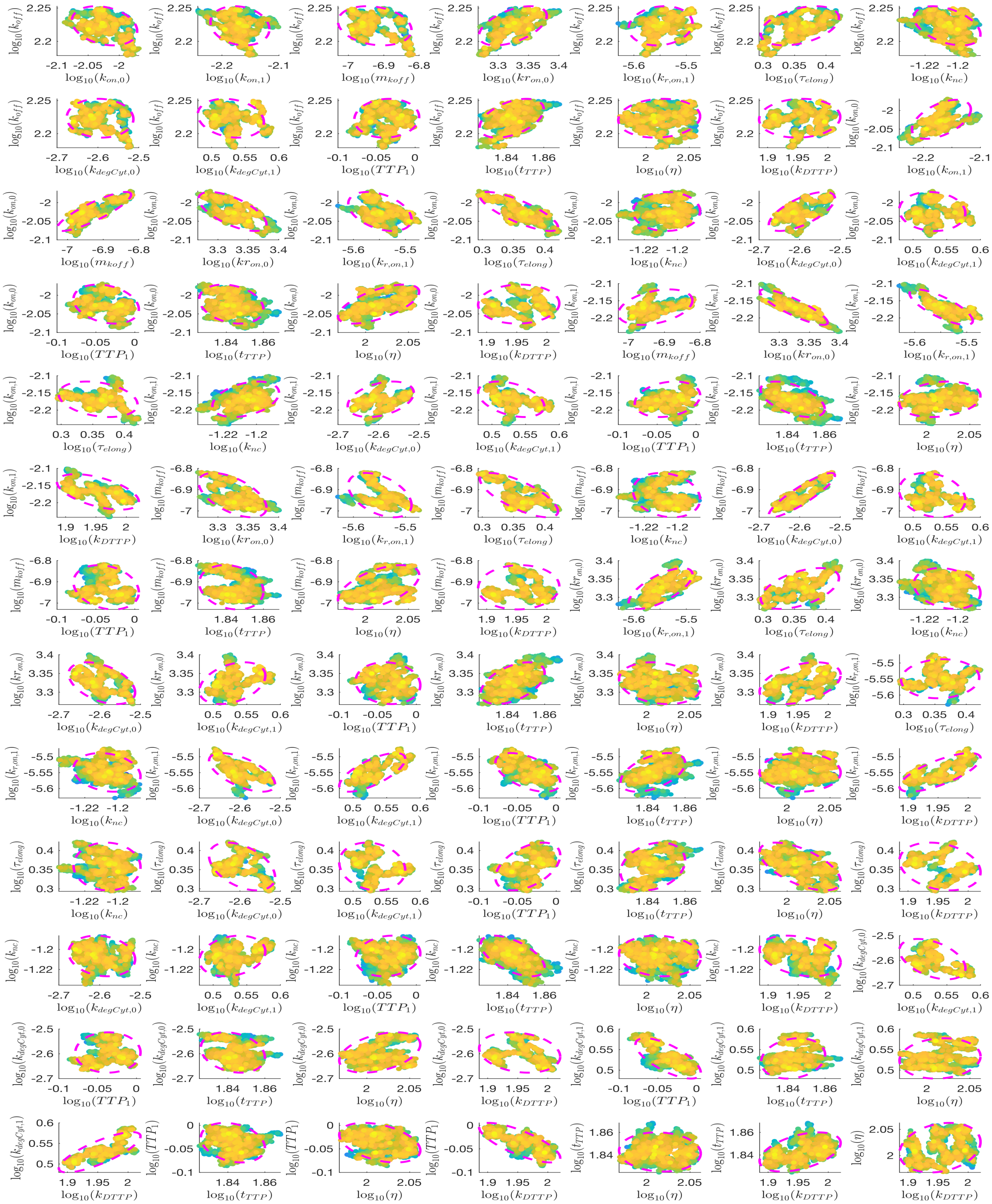

**Figure S15. Joint posterior distribution for the semi-mechanistic DUSP1 model.**
Uncertainty quantification for the semi-mechanistic DUSP1 model, obtained by Metropolis–Hastings MCMC sampling. Each subplot shows the pairwise joint distribution of MCMC samples for two model parameters (in $\log_{10}$ space), with points colored by sample density. The magenta dashed ellipse marks the 95% confidence interval.

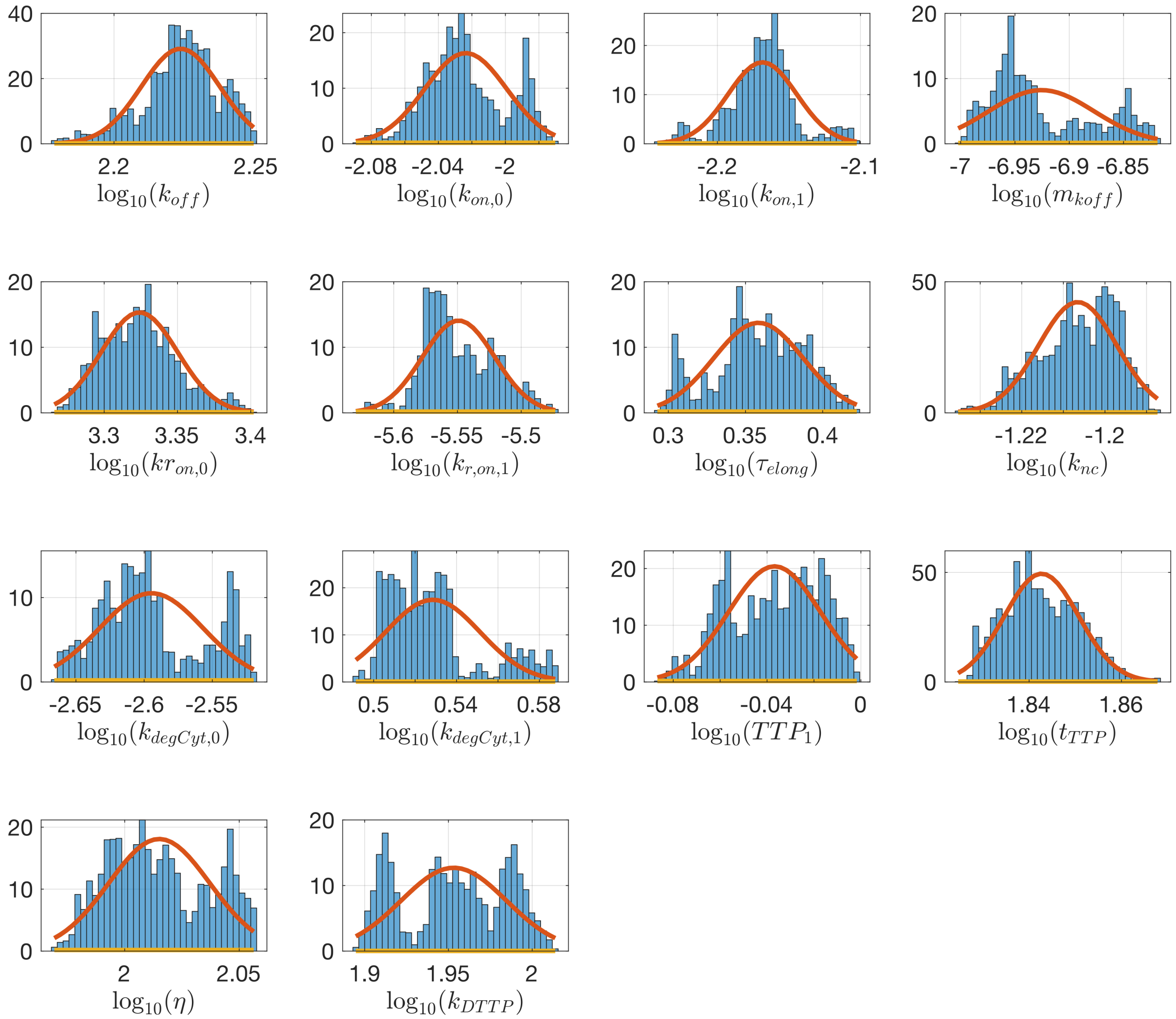

**Figure S16. Marginal posterior distributions for the semi-mechanistic DUSP1 model.**
Single-parameter marginal posteriors for the semi-mechanistic DUSP1 model, obtained by Metropolis–Hastings MCMC sampling. Each panel shows one model parameter (in $\log_{10}$ space): the blue histogram is the distribution of MCMC samples, the red curve is a kernel density estimate of the marginal posterior, and the gold curve is the prior.

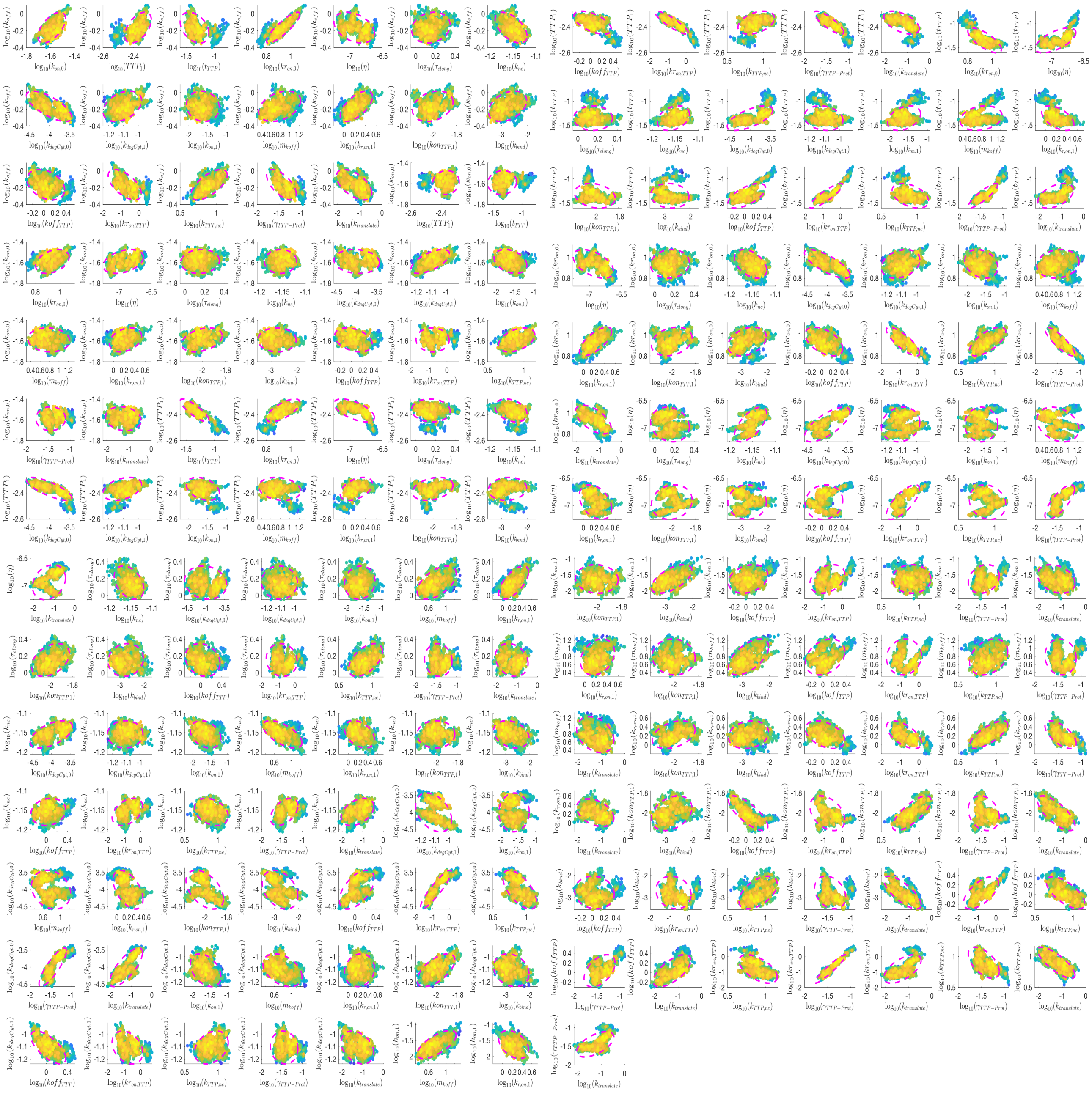

**Figure S17. Joint posterior distribution for the mechanistic DUSP1 model.**
Uncertainty quantification for the mechanistic DUSP1 model, obtained by Metropolis–Hastings MCMC sampling. Each subplot shows the pairwise joint distribution of MCMC samples for two model parameters (in $\log_{10}$ space), with points colored by sample density. The magenta dashed ellipse marks the 95% confidence interval.

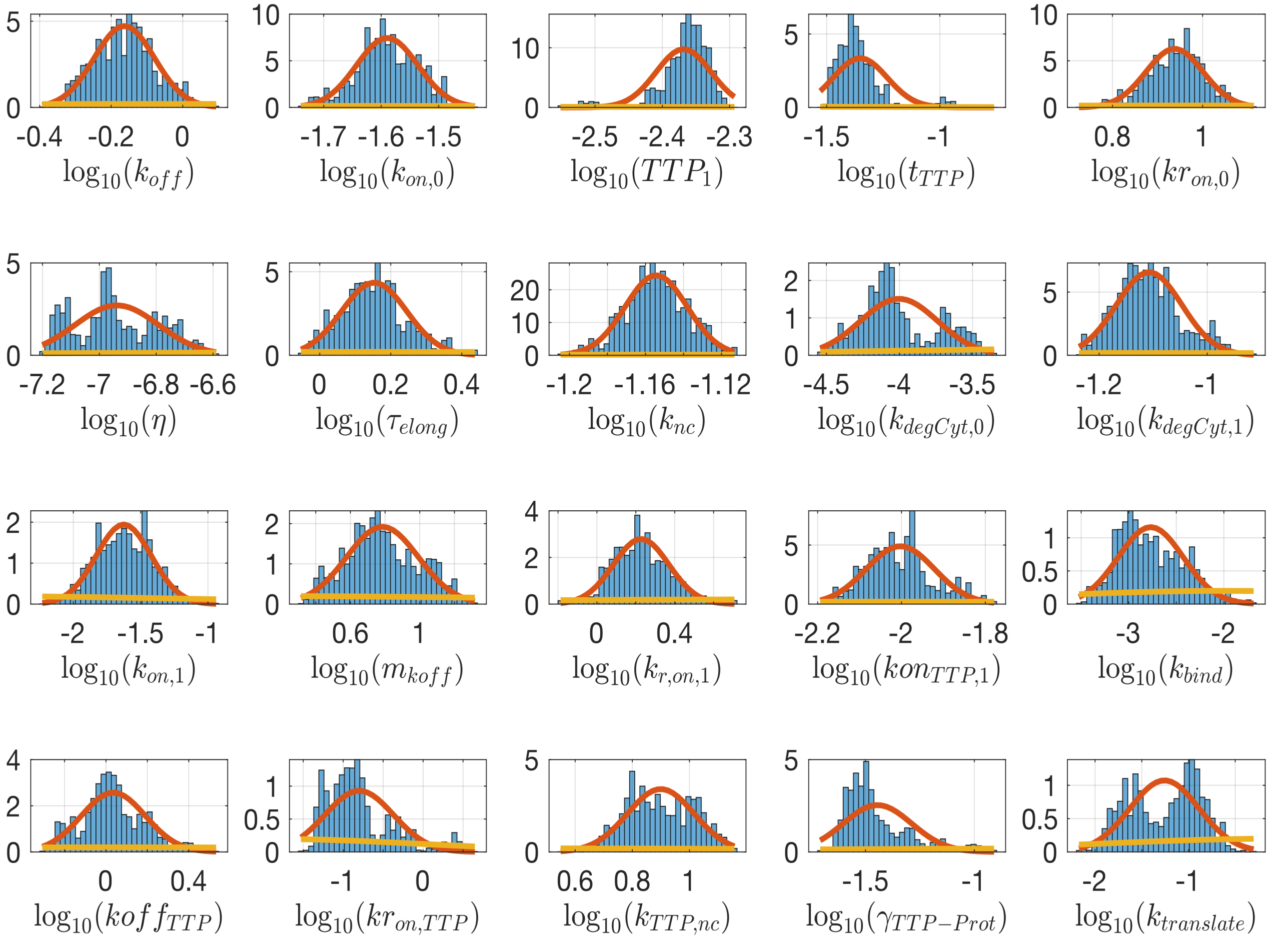

**Figure S18. Marginal posterior distributions for the mechanistic DUSP1 model.**
Single-parameter marginal posteriors for the mechanistic DUSP1 model, obtained by Metropolis–Hastings MCMC sampling. Each panel shows one model parameter (in $\log_{10}$ space): the blue histogram is the distribution of MCMC samples, the red curve is a kernel density estimate of the marginal posterior, and the gold curve is the prior.

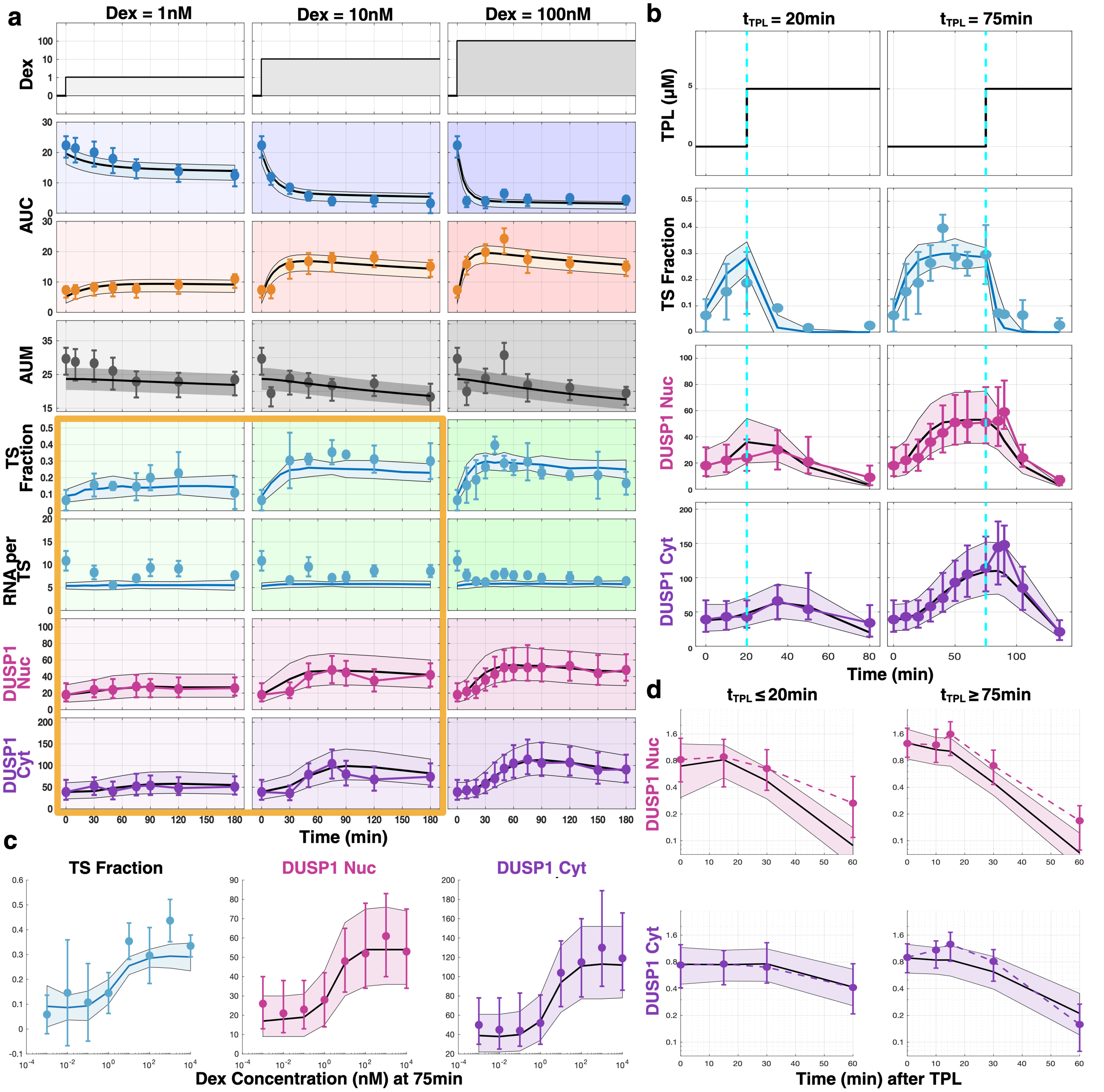

**Figure S19. Mechanistic TTP model predictions across Dex concentrations and transcriptional-inhibition conditions.** As in Fig. 4, but with all fits and predictions generated by the full mechanistic DUSP1-TTP model. **(a)** Joint GR and *DUSP1* time courses at 1, 10, and 100 nM Dex. The top three rows show fits to cytoplasmic GR (AUC, blue), nuclear GR (AUC, orange), and total GR mass (AUM, gray); the bottom four show *DUSP1* TS fraction, RNA per TS, nuclear mRNA, and cytoplasmic mRNA. The model was fit to the GR data and the 100 nM *DUSP1* column; the gold outline marks the 1 and 10 nM columns, which are predictions generated without refitting. **(b)** Dex–TPL dual-stimulation predictions for early ($t_{\mathrm{TPL}} = 20$ min) and late ($t_{\mathrm{TPL}} = 75$ min) transcriptional inhibition. Cells were stimulated with 100 nM Dex, then 5 µM TPL at $t_{\mathrm{TPL}}$ (cyan dashed lines). **(c)** Dex-titration predictions at $t = 75$ min spanning 1 pM to 10 µM. **(d)** Compartment-specific *DUSP1* mRNA decay after TPL, grouped into early ($t_{\mathrm{TPL}} \leq 20$ min) and late ($t_{\mathrm{TPL}} \geq 75$ min) inhibition; the y-axis is compartment-specific mRNA normalized for TPL onset, *DUSP1* elongation, and nuclear-to-cytoplasmic export. The mechanistic TTP model reproduces the data with accuracy comparable to the semi-mechanistic model (Fig. 4), with a slight reduction in predicted TS fraction across the Dex titration (c). In (a) and (c), TS fraction and RNA per TS are mean ± standard deviation; all other data are median ± interquartile range (25th/75th).

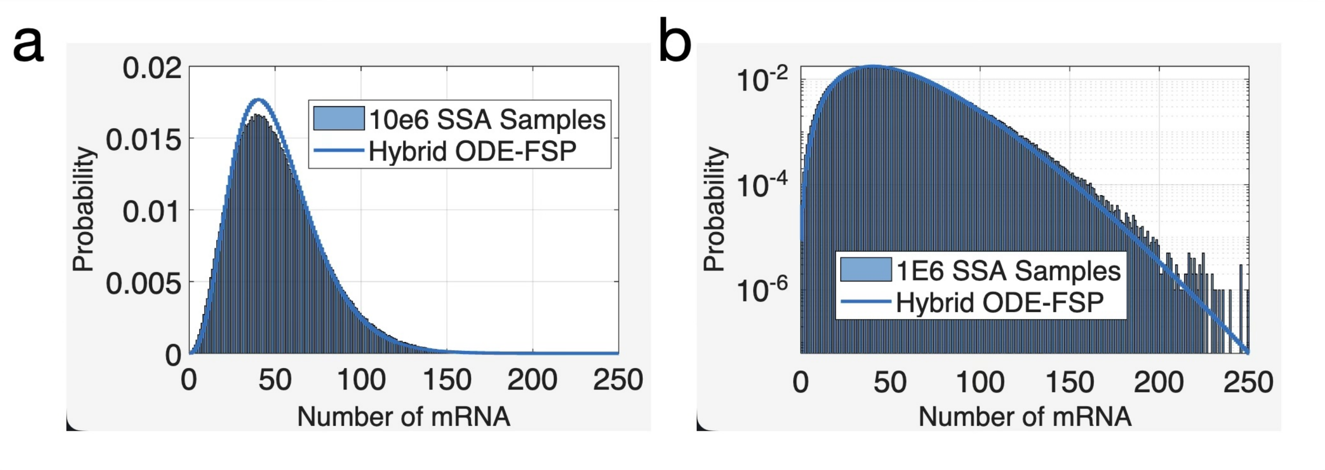

**Figure S20. Verification of the hybrid FSP solution.**
**(a)** Comparison of the marginal distribution for nuclear *DUSP1* mRNA at $t = 180$ min following 100 nM Dex for the hybrid ODE-FSP solution (line) and $10^6$ trajectories from Gillespie's Stochastic Simulation Algorithm (bars). **(b)** Same as (a) but with probabilities presented on a log scale.

#### Supplementary Tables

**Table S1.** Per-cell variability of GR fluorescence by Dex concentration and stimulation time.

| Dex (nM) | Time (min) | $n$ | Nuclear GR mean | Nuclear GR SD | Cytoplasmic GR mean | Cytoplasmic GR SD | N/C ratio mean | N/C ratio SD |
|---|---|---|---|---|---|---|---|---|
| 0 | 0 | 420 | 7 | 3 | 22 | 6 | 0.32 | 0.10 |
| 1 | 10 | 310 | 7 | 3 | 21 | 6 | 0.33 | 0.12 |
| 1 | 30 | 301 | 8 | 4 | 20 | 6 | 0.41 | 0.13 |
| 1 | 50 | 340 | 8 | 4 | 18 | 5 | 0.45 | 0.15 |
| 1 | 75 | 312 | 8 | 3 | 15 | 4 | 0.50 | 0.17 |
| 1 | 120 | 331 | 9 | 3 | 14 | 4 | 0.66 | 0.21 |
| 1 | 180 | 291 | 11 | 4 | 13 | 4 | 0.90 | 0.22 |
| 10 | 10 | 293 | 8 | 3 | 12 | 3 | 0.63 | 0.24 |
| 10 | 30 | 272 | 15 | 5 | 8 | 2 | 1.89 | 0.73 |
| 10 | 50 | 272 | 17 | 5 | 6 | 2 | 3.12 | 1.14 |
| 10 | 75 | 259 | 18 | 6 | 4 | 1 | 4.57 | 1.32 |
| 10 | 120 | 311 | 18 | 4 | 4 | 2 | 5.02 | 2.50 |
| 10 | 180 | 309 | 15 | 4 | 3 | 3 | 5.23 | 3.99 |
| 100 | 10 | 308 | 16 | 4 | 4 | 2 | 4.96 | 2.47 |
| 100 | 30 | 300 | 20 | 5 | 4 | 2 | 6.63 | 3.49 |
| 100 | 50 | 298 | 24 | 6 | 6 | 2 | 4.24 | 1.79 |
| 100 | 75 | 272 | 17 | 5 | 4 | 2 | 4.47 | 1.72 |
| 100 | 120 | 302 | 16 | 4 | 5 | 2 | 3.64 | 1.68 |
| 100 | 180 | 284 | 15 | 4 | 4 | 1 | 3.61 | 1.25 |

**Table S2.** Comparison of competing models to explain GR transport dynamics based on maximum a posterior estimate (MAP), AIC, BIC and approximate Bayes Factors based on BIC.

| | Import, $\gamma_{nuc}, \gamma_{cyt}$ | Import, $\gamma_{cyt}$ | Import, $\gamma_{nuc}$ | Export, $\gamma_{nuc}, \gamma_{cyt}$ |
|---|---|---|---|---|
| Num. Parameters | 8 | 7 | 7 | 8 |
| $\log(MAP/MAP_{best})$ | 0 | -913 | -128 | -19194 |
| AIC - $AIC_{best}$ | 0 | 1825 | 253 | 38388 |
| BIC - $BIC_{best}$ | 0 | 1818 | 246 | 38388 |
| $\log(aBF_{BIC})$ | 0 | -909 | -123 | -19194 |

**Table S3.** Inferred model parameter values and uncertainties for Dex-driven GR transport and degradation.

| Description (units) | Parameter | MLE Value | Posterior Mean ± STD |
|---|---|---|---|
| GR transport Cyt→Nuc, basal ($\min^{-1}$) | $k_{cn0}$ | 6.02e-03 | 6.04e-03 ± 1.65e-04 |
| GR transport Cyt→Nuc, Dex-dependent ($\min^{-1}$) | $k_{cn1}$ | 1.15e-01 | 1.15e-01 ± 2.56e-03 |
| Dex degradation rate, effectively zero ($\min^{-1}$) | $g_{Dex}$ | 6.76e-07 | 1.66e-06 ± 3.15e-06 |
| GR transport Nuc→Cyt ($\min^{-1}$) | $k_{nc}$ | 1.88e-02 | 1.87e-02 ± 6.61e-04 |
| GR synthesis rate, Cyt ($\min^{-1}$) | $k_{GR}$ | 4.82e-02 | 4.80e-02 ± 9.20e-03 |
| GR degradation, Cyt ($\min^{-1}$) | $\gamma_{GRcyt}$ | 1.24e-03 | 1.24e-03 ± 3.42e-04 |
| GR degradation, Nuc ($\min^{-1}$) | $\gamma_{GRnuc}$ | 4.40e-03 | 4.37e-03 ± 4.47e-04 |
| Michaelis constant for Dex-induced import (nM) | $M_{Dex}$ | 1.17e+01 | 1.19e+01 ± 2.84e-01 |

**Table S4.** Comparison of competing models to explain *DUSP1* TS and Nuclear mRNA dynamics based on MAP, AIC, BIC and approximate Bayes Factors.

| | $K_{OFF}$ | $K_{ON}$ | $K_R$ | $K_{OFF}, K_{ON}$ | $K_{OFF}, K_{ON}, K_R$ |
|---|---|---|---|---|---|
| Num. Parameters | 7 | 7 | 7 | 8 | 9 |
| $\log(MAP/MAP_{\text{best}})_{Nuc}$ | -54 | -16 | -218 | 0 | -0 |
| $\log(MAP/MAP_{\text{best}})_{TS}$ | -90 | 0 | -132 | -11 | -11 |
| $\log(MAP/MAP_{\text{best}})_{Tot}$ | -133 | -5 | -339 | 0 | -0 |
| AIC - $\text{AIC}_{\text{best}}$ | 264 | 9 | 676 | 0 | 2 |
| BIC - $\text{BIC}_{\text{best}}$ | 258 | 2 | 669 | 0 | 8 |
| $\log(BF_{\text{approx}})$ | -129 | -1 | -335 | 0 | -4 |

**Table S5.** Inferred model parameter values and uncertainties for semi-mechanistic model of *DUSP1* transcription regulation.

| Description (units) | Parameter | MLE Value | Posterior Mean ± STD |
|---|---|---|---|
| Basal gene inactivation rate ($\min^{-1}$) | $k_{off}$ | 6.99e+01 | 7.77e+01 ± 1.96e+01 |
| Basal gene activation rate ($\min^{-1}$) | $k_{on,0}$ | 4.06e-03 | 7.00e-03 ± 2.40e-03 |
| nucGR-dependent gene activation rate ($\min^{-1}$) | $k_{on,1}$ | 6.97e-03 | 6.82e-03 ± 2.79e-04 |
| nucGR modulation of inactivation ($\text{molecules}^{-1}$) | $m_{koff}$ | 2.25e-07 | 1.24e-07 ± 8.00e-08 |
| Basal DUSP1 transcription rate ($\min^{-1}$) | $kr_{on,0}$ | 8.80e+02 | 9.80e+02 ± 2.20e+02 |
| nucGR-dependent DUSP1 transcription rate ($\min^{-1}$) | $k_{r,on,1}$ | 4.42e-07 | 8.56e-07 ± 5.22e-07 |
| DUSP1 elongation time (min) | $\tau_{elong}$ | 1.76e+00 | 1.89e+00 ± 3.15e-01 |
| DUSP1 nuclear export rate ($\min^{-1}$) | $k_{nc}$ | 6.07e-02 | 6.16e-02 ± 2.65e-03 |
| Basal DUSP1 mRNA degradation rate ($\min^{-1}$) | $k_{degCyt,0}$ | 1.58e-03 | 9.30e-04 ± 2.82e-04 |
| Max TTP-dependent degradation rate ($\min^{-1}$) | $k_{degCyt,1}$ | 3.53e+00 | 4.05e+00 ± 6.15e-01 |
| Max fold-change in TTP-mediated degradation | $TTP_1$ | 1.02e+00 | 9.00e-01 ± 1.16e-01 |
| Half-max time for TTP activity (min) | $t_{TTP}$ | 6.82e+01 | 6.86e+01 ± 1.44e+00 |
| Hill coefficient, TTP time-dependence | $\eta$ | 2.68e+01 | 2.68e+01 ± 8.46e+00 |
| TTP-mRNA binding saturation (molecules) | $k_{DTTP}$ | 1.07e+02 | 1.15e+02 ± 1.26e+01 |

**Table S6.** Summary of model fit and prediction errors for TS and nuclear and cytoplasmic *DUSP1* at various Dex induction levels and transcriptional inhibition times. For each condition, prediction errors are relative to the best performing model for that condition.

| Data Set | Linear Constant | Nonlinear Constant | Linear Time-Varying | Nonlinear Time-Varying | Detailed Mechanistic |
|---|---|---|---|---|---|
| Nuc/Cyt, 100 nM Dex (Fit) | -124 | -214 | -103 | -4 | 0 |
| TS, 100 nM Dex (Fit) | -4 | -2 | -3 | 0 | -2 |
| Nuc/Cyt, 0 min Tpl (Fit) | -19 | -28 | -32 | -16 | 0 |
| **Total Fit Error** | -147 | -245 | -139 | -20 | -2 |
| Nuc/Cyt, 0.3 nM Dex (Predict) | -27 | -29 | -48 | -26 | 0 |
| Nuc/Cyt, 1.0 nM Dex (Predict) | -40 | -64 | -41 | -31 | 0 |
| Nuc/Cyt, 10 nM Dex (Predict) | -39 | -77 | -44 | -6 | 0 |
| TS, 0.3 nM Dex (Predict) | -1 | -1 | -0 | 0 | -4 |
| TS, 1.0 nM Dex (Predict) | -1 | -0 | -0 | 0 | -2 |
| TS, 10 nM Dex (Predict) | -2 | -2 | -1 | 0 | -0 |
| Nuc/Cyt, 20 min Tpl (Predict) | -10 | -36 | -5 | 0 | -2 |
| Nuc/Cyt, 75 min Tpl (Predict) | 0 | -8 | -4 | -0 | -2 |
| Nuc/Cyt, 150 min Tpl (Predict) | -21 | -36 | -16 | 0 | -9 |
| Nuc/Cyt, 180 min Tpl (Predict) | -7 | -28 | -8 | -9 | 0 |
| **Total Prediction Error** | -147 | -282 | -169 | -73 | -19 |

**Table S7.** Inferred DUSP1- and TTP-related parameter values across the three model variants. Em-dashes (—) indicate parameters not present in that model. Units shown in parentheses; dimensionless parameters are unmarked.

| Description (units) | Parameter | TS/Nuc ($k_{on}, k_{off}, k_r$) | TS/Nuc ($k_{on}$ only) | TS/Nuc/Cyt | Extended |
|---|---|---|---|---|---|
| Basal gene inactivation rate ($min^{-1}$) | $k_{off}$ | 1.83e+00 | 1.73e+00 | 6.99e+01 | 6.11e-01 |
| Basal gene activation rate ($min^{-1}$) | $k_{on,0}$ | 1.63e-02 | 1.00e-02 | 4.06e-03 | 3.55e-02 |
| nucGR-dependent gene activation rate ($min^{-1}$) | $k_{on,1}$ | 3.11e-03 | 3.63e-03 | 6.97e-03 | 3.48e-03 |
| nucGR modulation of inactivation ($molecules^{-1}$) | $m_{koff}$ | 1.74e-02 | - | 2.25e-07 | 7.86e-02 |
| Basal DUSP1 transcription rate ($min^{-1}$) | $kr_{on,0}$ | 1.77e+01 | 2.08e+01 | 8.80e+02 | 6.79e+00 |
| nucGR-dependent DUSP1 transcription rate ($min^{-1}$) | $kr_{on,1}$ | 9.97e-06 | - | 4.42e-07 | 2.44e-07 |
| DUSP1 elongation time (min) | $\tau_{elong}$ | 2.67e+00 | 2.67e+00 | 1.76e+00 | 1.36e+00 |
| DUSP1 nuclear export rate ($min^{-1}$) | $k_{nc}$ | 3.17e-02 | 3.16e-02 | 6.07e-02 | 7.12e-02 |
| Basal DUSP1 mRNA degradation rate ($min^{-1}$) | $\gamma_{Cyt,0}$ | - | - | 1.58e-03 | 5.07e-04 |
| Max TTP-dependent degradation rate ($min^{-1}$) | $\gamma_{Cyt,1}$ | - | - | 3.53e+00 | 8.74e-02 |
| Max fold-change in TTP-mediated degradation | $TTP1$ | - | - | 1.02e+00 | - |
| Half-max time for TTP activity (min) | $t_{TTP}$ | - | - | 6.82e+01 | - |
| Hill coefficient, TTP time-dependence | $\eta$ | - | - | 2.68e+01 | - |
| TTP-mRNA binding saturation (molecules) | $kD_{TTP}$ | - | - | 1.07e+02 | - |
| nucGR-dependent TTP gene activation rate ($min^{-1}$) | $kon_{TTP,1}$ | - | - | - | 2.10e-02 |
| TTP gene inactivation rate ($min^{-1}$) | $koff_{TTP}$ | - | - | - | 2.26e+00 |
| TTP transcription rate ($min^{-1}$) | $kr_{on,TTP}$ | - | - | - | 1.02e+00 |
| TTP nuclear export rate ($min^{-1}$) | $k_{TTP,nc}$ | - | - | - | 1.17e-02 |
| Basal TTP mRNA degradation rate ($min^{-1}$) | $\gamma_{TTP,Cyt}$ | - | - | - | 2.09e-03 |
| TTP translation rate ($min^{-1}$) | $k_{translate}$ | - | - | - | 6.65e-01 |
| TTP protein degradation rate ($min^{-1}$) | $\gamma_{CTT,Prot}$ | - | - | - | 1.42e+00 |
| TTP–mRNA binding rate ($min^{-1}$ $mol^{-1}$) | $k_{bind}$ | - | - | - | 4.21e+00 |
| TTP–mRNA unbinding rate ($min^{-1}$) | $k_{unbind}$ | - | - | - | 4.11e-02 |
| Hill coefficient, TTP–mRNA binding | $\eta_{TTP}$ | - | - | - | 7.48e-02 |
